# Targeted Magnetic Nanodiscs for Wireless Causal Manipulation of Gut-Brain Circuits

**DOI:** 10.64898/2026.03.26.714579

**Authors:** Ye Ji Kim, Nasim Biglari, Taylor M. Cannon, Cameron Forbrigger, Scott Machen, Emmanuel Vargas Paniagua, Karen K. L. Pang, Jessica Slaughter, Jacob L. Beckham, Keisuke Nagao, Elizabeth Whittier, Florian Koehler, Rebecca Leomi, Polina Anikeeva

## Abstract

Causal manipulation of gut-brain neural circuits empowers studies of metabolism and interoception. However, the anatomy and cytoarchitecture of peripheral ganglia relaying gut-brain circuits pose challenges to deployment of optical or electrical stimulation probes. To enable implant-free, cell-type specific, and temporally precise control of defined gut-brain pathways, we develop a neuromodulation platform based on magnetic nanodiscs (MNDs) targeted to peripheral neurons via genetically delivered anchoring moieties. The anchored MNDs selectively transduce externally applied weak magnetic fields to mechanical torque, thereby activating endogenous mechanosensitive pathways in specified cell types with sub-second latency. When targeted to nodose ganglia neurons expressing oxytocin or glucagon-like peptide 1 receptors, MND-mediated stimulation enables robust and reversible activation of gut-brain signaling, which engages hindbrain satiety circuits and regulates feeding behavior. These findings establish MND-mediated stimulation as a genetically targetable, implant-free strategy for modulating gut-brain neural circuits and highlight its potential in studies of brain-body physiology and bioelectronic medicines.

## Introduction

Bidirectional communication between the gastrointestinal (GI) tract and the central nervous system underlies energy balance and homeostasis, and its contributions to motivation and reward behaviors are increasingly recognized, inspiring the development of technologies for causal manipulation of gut-brain neural circuits ^1-5^. However, the anatomy and cellular architecture of peripheral ganglia that relay gut-brain neural signals present obstacles to deploying implantable electrical and optical hardware commonly used in causal studies of brain function ^6-8^. Electrical neuromodulation offers millisecond temporal precision but does not discriminate between distinct cell types, and chronic implantation of electrodes into miniscule peripheral nerves and ganglia is challenging in rodent models. Optogenetic neuromodulation, deployed in genetically modified model organisms, permits bidirectional control of specific cell types but also requires implanted or wearable optical hardware ^9-11^. Chemogenetics using Designer Receptors Exclusively Activated by Designer Drugs (DREADDs) remains a dominant tool for bidirectional interrogation of genetically identified gut-brain pathways as it relies on systemic delivery of designer ligands (e.g. clozapine N-oxide, CNO) eliminating the need for implants ^12-14^. However, chemogenetic manipulation unfolds over minutes, limiting one’s ability to distinguish between millisecond-scale neuronal and minute-scale endocrine contributions to gut-brain communication ^15,16^. Consequently, there remains a need for implant-free, cell-type specific and temporally precise neuromodulation approaches deployable in peripheral nerves and ganglia relaying signals from the gut and other peripheral organs to the brain.

Magnetic fields (MFs) offer unprecedented access to deep tissues ^17^, and magnetic nanoparticles have emerged as minimally invasive actuators locally converting biologically benign MFs into thermal ^18-21^, electrical ^20-22^, or mechanical ^23-25^ cues that trigger neuronal activity with sub-second precision ^22^. Solubility of magnetic nanomaterials in physiological media permits their delivery via injections and enables distributed stimulation, which is uniquely suited to multiscale architecture of the gut-brain circuits spanning centimeter-long nerves, millimeter-scale ganglia, ^26^ and micrometer-thick enteric (gut) plexuses ^27^.

Magnetomechanical stimulation, which harnesses magnetic nanomaterials to transduce weak (<100 mT), slow-varying (<100 Hz) MFs into torque, offers a practical neuromodulation strategy deployable across in vitro and in vivo assays due to the simplicity of the required magnetic instrumentation ^23-25^. Although this approach has been shown to elicit calcium ion (Ca^2+^) influxes in subpopulations of diverse cell types ^23,28^, achieving temporally precise and targeted magnetomechanical neural stimulation has so far demanded exogenous expression of known mechanoreceptor transgenes ^25,28,29^, potentially reflecting limited transduction efficiency at the chosen conditions.

Here, we evaluate magnetite nanodiscs (MNDs) as transducers of magnetomechanical stimulation across a broad range of MF conditions and identify regimes where forces generated by MNDs anchored to cell membranes are sufficient to engage mechanoreceptors with pN-scale thresholds ^30^. We then leverage covalent bonding between SNAP-tag protein and a benzyl guanine (BG) moiety to target MNDs to specific neurons and achieve their selective excitation with sub-second precision. This cell-type specific MND stimulation is then evaluated in the mouse nodose ganglia - critical relays of vagal gut-brain circuits anatomically inaccessible to implanted hardware (**Fig. 1a**). Wireless magnetomechanical stimulation of oxytocin receptor (Oxtr^+^) or glucagon-like peptide receptor (Glp1r^+^) expressing neurons in the left nodose ganglia suppresses feeding and yields regional upregulation of hindbrain satiety circuits. Together these observations establish targeted MNDs as injectable tools for temporally precise causal interrogation of genetically defined body-brain circuits.

**Fig. 1.**
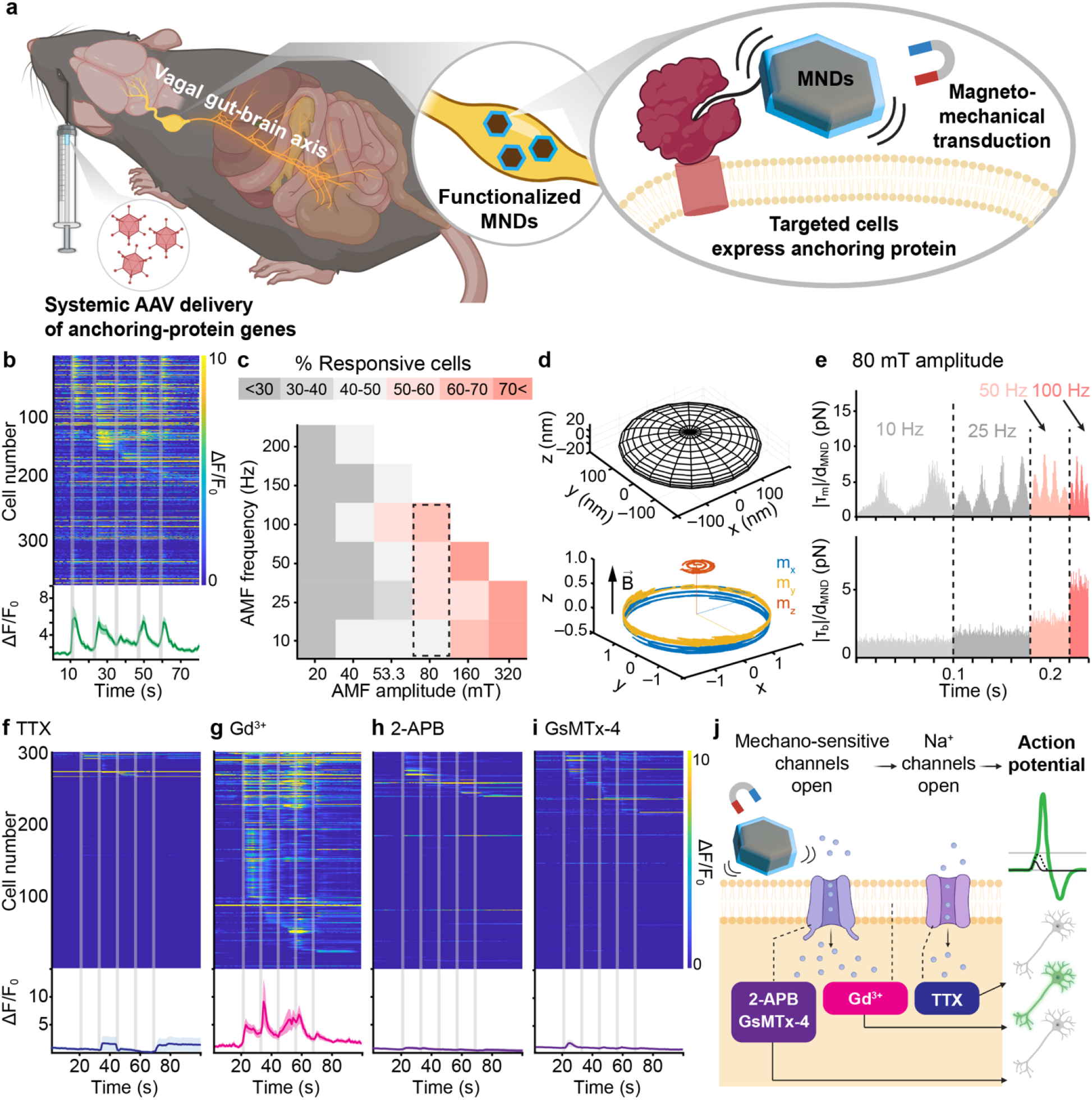
Magnetomechanical neuromodulation with magnetic nanodiscs (MNDs). **a**, Magnetomechanical neuromodulation platform comprising: (1) systemic viral delivery of SNAP-tag gene to defined cell types; (2) MNDs functionalized with benzylguanine (BG) that covalently binds to SNAP-tag (inset); and (3) Cre-driver mouse lines to direct SNAP expression to specific vagal neurons transducing gut-brain signaling. **b**, Relative fluorescence traces of GCaMP6s of individual neurons (top) and their average (bottom) in response to 2 s magnetic field (MF, 80 mT, 50 Hz). **c**, A diagram showing the percentage of neurons responding to various MF amplitude and frequency combinations, assessed by GCaMP6s imaging. **d**, Geometry (top) of simulated MNDs (230 nm diameter, 30 nm thickness) and simulated magnetization trajectories during four MF cycles (bottom). **e**, Simulated magnetic torque (T_m_) and torque generated by Brownian motion (T_b_), normalized by diameter of MNDs (d_MND_) across varying MF frequencies at a fixed amplitude (80 mT). **f-i** Dynamic GCaMP6s fluorescence imaging of MND-decorated hippocampal neurons treated with: (**f**) tetrodotoxin (TTX; sodium channel blocker), (**g**) gadolinium (Gd^3+^; membrane stiffening agent), (**h**) 2-APB (TRP channel and internal calcium store inhibitor), and (**i**) GsMTx-4 (selective mechanosensitive channel blocker), revealing the contribution of specific pathways. (**b**,**f-i**) Lines and shaded areas represent mean ±standard error of the mean (s.e.m.) **j**, Schematic summary of the proposed mechanism: MND-generated mechanical forces activate mechanosensitive ion channels, and subsequent depolarization triggers opening of voltage-gated sodium channels to initiate action potentials.

## Results and Discussion

### Endogenous mechanoreceptors mediate neuronal responses to magnetomechanical stimulation

To identify MF regimes for effective magnetomechanical neuronal excitation, we employed magnetite nanodiscs MNDs (229±34 nm in diameter, 30±9 nm thickness, Fig. S1, Methods) that were previously shown to elicit Ca^2+^ influxes in multiple cell classes in response to weak MFs ^25,31,32^. These particles were functionalized with an amphiphilic coating of poly(ethylene glycol) - poly(maleic anhydride-alt-1-octadecene) (PEG-PMAO, Methods) that enables their electrostatic association with cell membranes ^33,34^. The MNDs were then applied at a density of 26.3 *µ*g cm^-2^ to primary hippocampal neurons virally transduced to express a fluorescent Ca^2+^ indicator GCaMP6s, as a proxy for neural activity. By varying MF amplitude between 20-320 mT and frequency between 20-200 Hz, we found that at amplitudes ≥80 mT, MFs with frequencies ≥25 Hz evoked Ca^2+^ influxes in >50% neurons (**Fig. 1b,c**, Fig. S2-4, Methods). The fraction of responsive neurons increased with the MF frequency and amplitude (Fig. S3, S4), and further depended on the MF duration saturating at epochs ≥ 2s (Fig. S5 a-e). In contrast, the latency of the response did not depend on the epoch duration and was found to be 433 ± 244 ms at our camera rate of 20 fps (Fig. S5f).

Given that prior work on magnetomechanical neuromodulation concentrated on the effects of MF amplitude alone, we sought to mechanistically probe the observed synergy between the frequency and amplitude in driving the neuronal responses. We developed a computational model to describe dynamic magnetization behavior and hydrodynamic drag experienced by a MND (230 nm in diameter, 30 nm in thickness) at our experimental MF conditions (**Fig. 1d, Supplementary Note 1**, Fig. S6a, Supplementary Video S1). Here, the sum of magnetic (***T***_***m***_), hydrodynamic/ fluidic (***T***_***f***_), and Brownian motion (***T***_***b***_) torques experienced by a MND is assumed to be zero, as the particle rotational inertia is negligible compared to the magnetic and fluidic torques. Therefore, the dynamic equation of a MND constrained to rotate about a fixed axis can be written as:

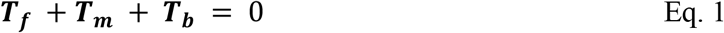

The force applied to the cell membrane due to MND movement is directly coupled to the viscous flow, captured by ***T***_***f***_, whose dependence on MF amplitude and frequency can be obtained by analyzing those of the ***T***_***m***_ and ***T***_***b***_.

Magnetic torque, ***T***_***m***_, scales with MF amplitude but does not explicitly depend on frequency (Eq. 2, **Fig. 1e**, Fig. S6).

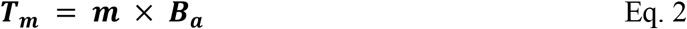

where ***m*** is the MND magnetic moment vector and ***B***_***a***_ is the applied magnetic flux density. Here ***m*** is varying along the probability density determined by the magnetic energy of an MND.

To account for the Brownian torque, ***T***_***b***_, in our dynamic model, we introduce a white noise term (**Supplementary Note 1**) and show that its intensity increases at higher frequencies manifesting in ***T***_***b***_ scaling with MF frequency (**Fig. 1e**, Fig. S6b). This scaling in part explains the weaker neuronal responses to low MF frequencies. Note that all MF frequencies examined here are 2-3 orders of magnitude below those necessary to observe inductive effects on neuronal activity ^35,36^.

Using our dynamic model we found that at 80 mT and 50 Hz, a 230 nm MND produces an average force of ‖***T***_***m***_ + ***T***_***b***_‖ / (MND diameter) = 2.5 Nm / 230 nm = 10.9 pN, which substantially exceeds the activation thresholds of known mechanosensitive ion channels (∼0.3 pN) ^30^ (**Fig. 1e**). Notably, not accounting for MND Brownian motion, leads to significant overestimation of magnetic forces (438 pN at 80 mT for 230 nm MNDs, Supplementary Note 2), masking the observed, but previously unexplained effects of MF frequency. Although the theoretical force values exceed mechanoreceptor activation thresholds at every amplitude and frequency simulated in this study, under experimental conditions magnetomechanical forces are likely partially dissipated by the surrounding matrix. Nonetheless, the simulation qualitatively explains the dependence of the cellular responses on the MF amplitude and frequency.

Given that our experiments (Fig. 1b,c) were conducted in hippocampal neurons without exogenous mechanoreceptor transgenes, we employed pharmacology to further probe the molecular mechanisms underlying the observed responses. In these experiments, hippocampal neurons expressing GCaMP6s and decorated with MNDs at a density of 26.3 *µ*g cm^-2^ were exposed to four 2s epochs of MF (80 mT, 50 Hz). We found that blocking voltage-gated sodium channels (VGSCs) with tetrodotoxin (TTX, 1 µM) was sufficient to abolish the observed magnetomechanical responses (**Fig. 1f, Fig. S7**). Given that VGSCs do not exhibit significant mechanosensitivity, we then evaluated contributions of cellular machinery known to respond to mechanical stimuli.

First, we found that gadolinium ions (Gd^3+^, 10 µM), which stiffen the plasma membrane without affecting the neuronal refractory period ^37,38^ did not reduce neuronal responsiveness to MND-mediated stimulation (**Fig. 1g, Fig. S7**), suggesting that membrane stiffness alone is not a limiting factor for the observed activation. We then evaluated the role of mechanosensitive Ca^2+^ transport machinery. Administration of 2-aminoethoxydiphenyl borate (2-APB, 100 µM), which inhibits Ca^2+^ release from intracellular stores and blocks the activity of mechanosensitive ion channels from the transient receptor potential (TRP) family ^39-43^ resulted in a significant reduction in neuronal responses to MF (**Fig. 1h, Fig. S7**). The addition of Grammostola mechanotoxin #4 (GsMTx-4, 5 µM), a selective inhibitor of cationic mechanosensitive channels, including those from the Piezo and TRP channel families ^44,45^, similarly led to a marked reduction in neuronal responses (**Fig. 1i, Fig. S7**). These findings support the role of endogenous mechanosensitive ion channels in initiating MND-mediated neuronal activation.

To assess whether intracellular Ca^2+^ release alone could account for the observed responses, we applied thapsigargin (Tg), an inhibitor of the sarcoplasmic/endoplasmic reticulum Ca^2+^ ATPase (SERCA), which depletes internal calcium stores and blocks store-operated calcium entry^46-48^. Treatment with 1 µM Tg had negligible effects on Ca^2+^ dependent GCaMP6s fluorescence change in response to MF (**Fig. S7**), suggesting that Ca^2+^ release from intracellular stores is not a major contributor to the observed response.

Taken together, these observations suggest that magnetomechanical forces exerted by MNDs activate nearby endogenous mechanoreceptors, and the response is amplified by the VGSCs to yield broad neuronal depolarization (**Fig. 1j**).

### Membrane targeting of MNDs enhances neuronal responses to magnetic stimulation

Although electrostatic association between MNDs and cell membranes is sufficient to mediate Ca^2+^ influxes into neurons in vitro, this interaction lacks cell-type selectivity of magnetomechanical stimulation and provides limited control over binding strength. To enable cell-type specific targeting and robust membrane anchoring of MNDs, we leveraged covalent bonding between SNAP-tag protein and a synthetic benzyl guanine (BG) probe (**Fig. 2a**). The reaction between SNAP-tag and BG is ubiquitously used for specific labeling of cells and molecules in vitro and in vivo ^49^. MNDs were functionalized with BG by employing BG-conjugated PEG combined with PMAO conjugated with Cy7 dye (**Fig. S8-10**, Methods).

**Fig. 2.**
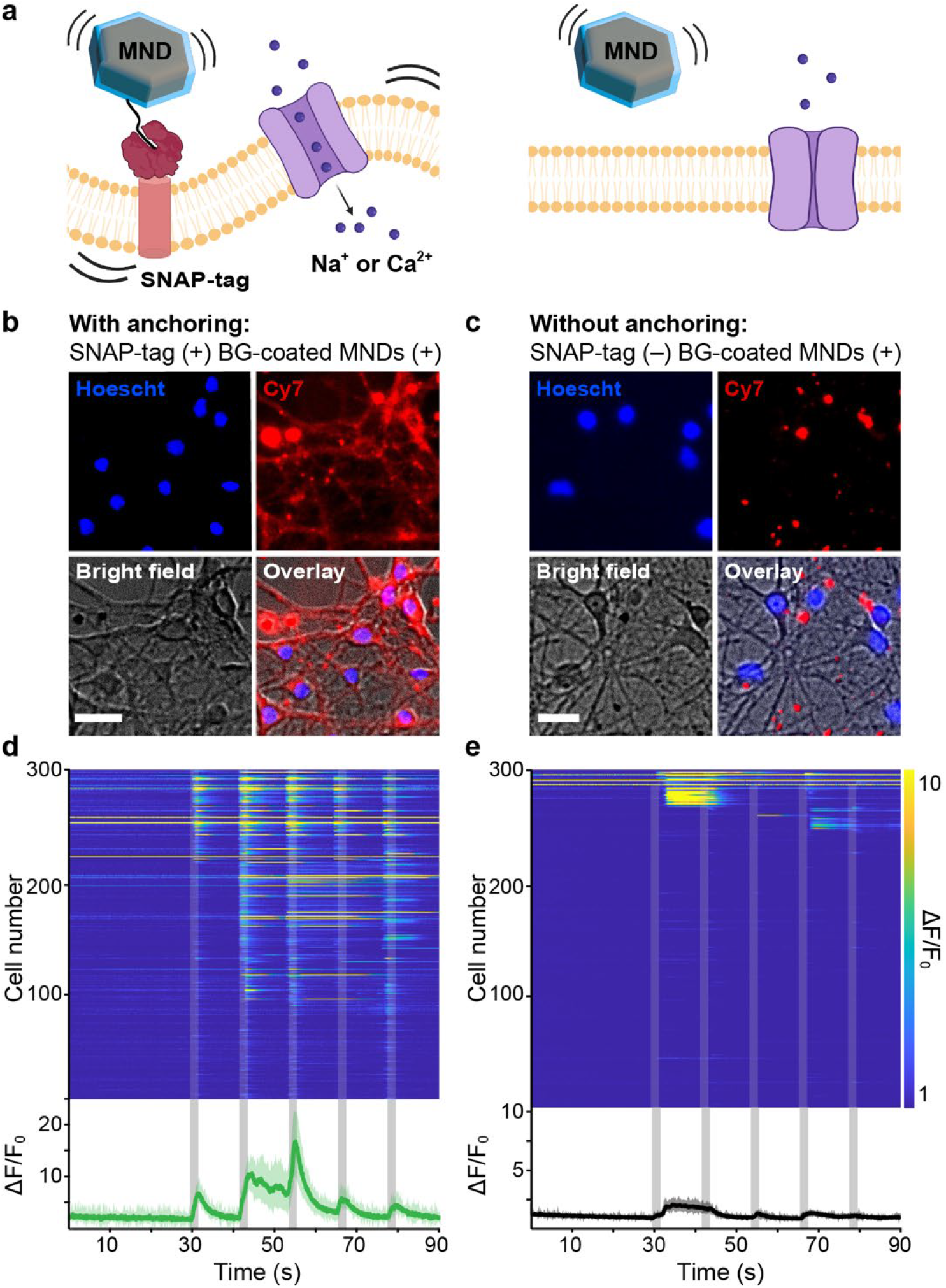
Genetic targeting of MNDs to neuronal membranes. **a**, Schematic illustrating cell-specific magnetomechanical neuromodulation, where membrane-anchored MNDs efficiently transmit mechanical torque to the cell surface under MFs. **b**, Fluorescence image of primary hippocampal neurons transduced with AAV9-Ef1α::SNAP-PDGFR and incubated with BG-coated MNDs (15.8 μg cm^−2^), showing uniform membrane localization. **c**, Fluorescence image of non-transduced neurons under identical conditions, showing MND aggregation and poor membrane localization. **d**, Dynamic GCaMP6s fluorescence imaging of SNAP-tag-expressing neurons exposed to MF (80 mT, 100 Hz, indicated by grey bars), showing time-locked activation. **e**, Dynamic GCaMP6s fluorescence in control neurons lacking SNAP-tag expression under identical MF conditions. The heatmap (top) shows the relative fluorescence traces of individual neurons, and the bottom plot shows the average. Scale bars, 25 μm. Lines and shaded areas represent mean ±s.e.m.

The attachment of BG- and Cy7-functionalized MNDs to the membranes was first assessed in neurons expressing SNAP-tag fused to a PDGFR transmembrane domain (Methods). Following incubation with functionalized MNDs (5.26 µg cm^−2^), Cy7 fluorescence was observed uniformly along the cell membrane (**Fig. 2b, Fig. S11**). In contrast, in neurons lacking SNAP-tag expression, MNDs (as denoted by Cy7) failed to localize to the membrane (**Fig. 2c, Fig. S12**).

Next, we evaluated whether covalent anchoring of MNDs to the membrane improves neuromodulation efficiency as compared to non-specific electrostatic association. The neurons were virally co-transduced to express GCaMP6s and SNAP-tag (Methods), and neurons expressing GCaMP6s alone were employed as controls. In both conditions, neurons were decorated with functionalized MNDs at a low surface density of 15.8 µg cm^-2^ (as compared to 26.3 µg cm^−2^ used for non-specific association in Fig. 1) and exposed to five epochs of MF (80 mT, 100 Hz, 2 s duration, 10 s interval). A significantly greater fraction of neurons expressing SNAP-tag responded to MF stimulation as compared to controls lacking SNAP-tag expression (50.5±31.7% vs 20.2±13.6%), while the response latency was comparable (740 ±33 ms vs 783±257 ms, **Fig. 2d,e** and **Fig. S13**, Supplementary Video S2). At a lower MND density (7.9 µg cm^-2^), MF application activated 34.8±18.9% of SNAP-expressing neurons, while neurons lacking SNAP-tag showed only a spontaneous level of response (5.7±3.6%). Further reducing MNDs to 4.0 µg cm^-2^ diminished responses, even in SNAP-tag-expressing neurons, to spontaneous levels (4.7±3.1%) (**Fig. S14**).

These observations indicate that covalent attachment of BG-functionalized MNDs to SNAP-tag expressing neurons enhances their response to MF stimulation allowing for selective activation of targeted cells while minimizing off-target effects.

### MND-mediated stimulation activates mechanosensory nodose ganglion neurons in vivo

Nodose ganglia (NGs) serve as key relays of gut-brain communication harboring cell bodies of bipolar mechanosensory and chemosensory neurons that transmit stretch and nutrient signals from the GI tract to the hind brain ^50,51^. Given the miniature dimensions and proximity to critical blood vessels ^26^, NGs are currently inaccessible to implantable neuromodulation hardware making them an important target for wireless neuromodulation with MNDs.

We first assessed the ability of targeted MNDs to activate mechanosensory NG neurons via Ca^2+^ imaging with GCaMP6s in mice expressing Cre recombinase in oxytocin receptor (Oxtr+) neurons (Oxtr-Cre^+/–^, heterozygous, Methods) (**Fig. 3a, Fig. S15**,**16**). Mechanosensory Oxtr+ vagal neurons with endings distributed throughout the small intestine transmit stretch signals associated with food ingestion to the brain, and their selective activation has been shown to reduce food intake making them a robust biological model for mechanical stimulation ^52^.

**Fig. 3.**
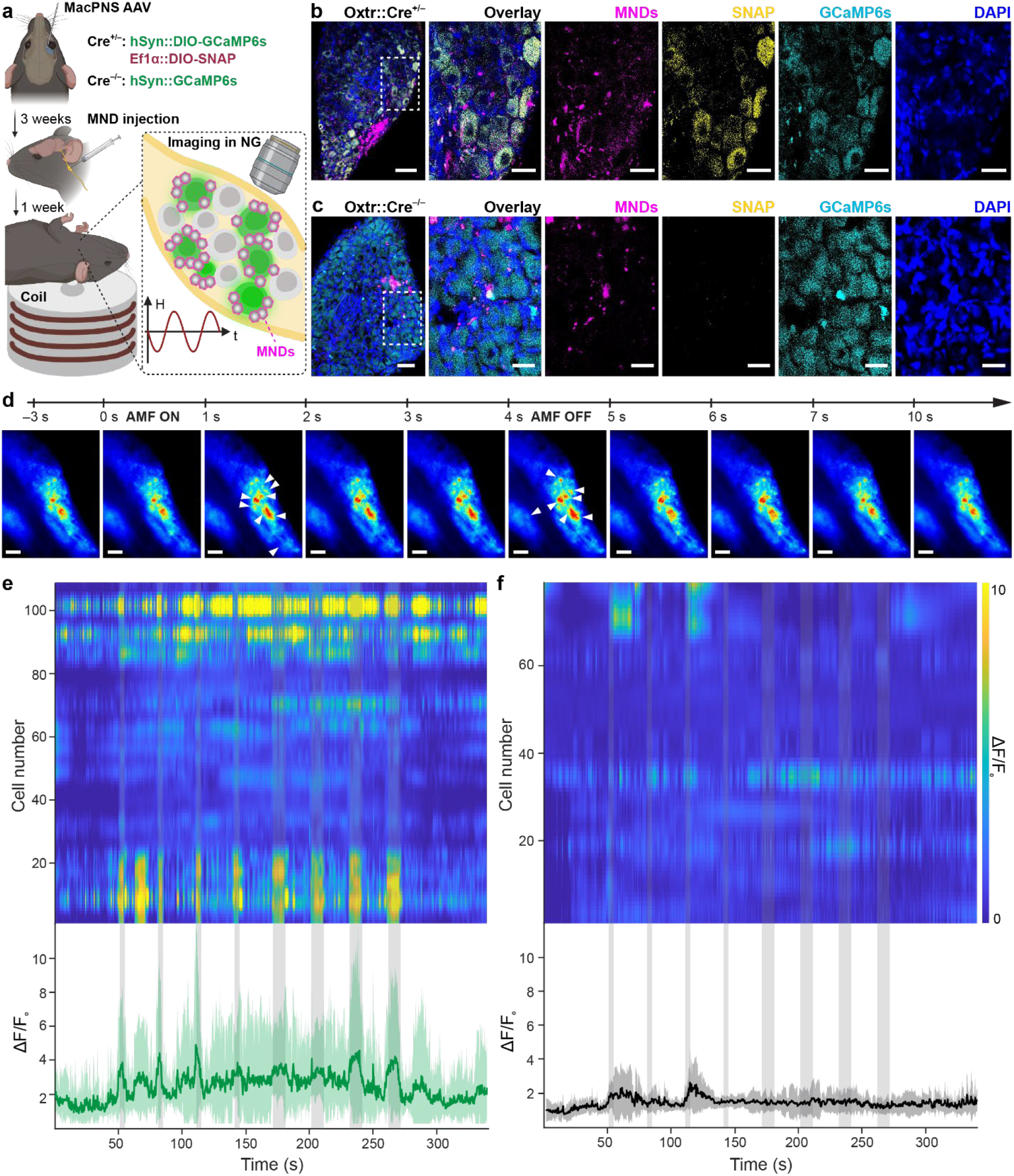
MND-mediated activation of targeted nodose ganglion neurons. **a**, Simultaneous calcium imaging and magnetic stimulation of nodose ganglia (NG) neurons expressing GCaMP6s and targeted with MNDs. **b**, Fluorescence image of the *left* NG in an Oxtr-Cre^+^/^−^ mouse showing expression of GCaMP6s and Cy7-labeled MNDs localized to SNAP-tag-expressing cells. **c**, Fluorescence image of the left NG in a Cre-negative (Oxtr-Cre^−^/^−^) control mouse, lacking SNAP expression and MND anchoring. (**b**,**c**) Scale bars in the left-most panels are 100 μm, and others are 25 μm. **d**, Representative trace of bulk GCaMP6s fluorescence during four 4-second MF pulses (80 mT, 50 Hz) in Oxtr-Cre^+^/^−^ mice, showing robust, time-locked activation (regions exhibiting prominent fluorescence changes at during MF epochs are indicated by white arrows). Scale bars, 100 μm. **e-f**, Single-cell GCaMP6s ΔF/F_0_ calcium traces and population averages in (**e**), Oxtr-Cre^+^/^−^ (male n=4) mice and (**f**) Oxtr-Cre^−^/^−^ (male n=3) mice during four 4-second and four 10-second MF pulses (80 mT, 50 Hz). Lines and shaded areas represent mean ±s.e.m.

To achieve SNAP-tag and GCaMP6s expression in Oxtr+ NG neurons, Oxtr-Cre^+/–^ mice were systemically (retro-orbitally) injected with a mixture of MaCPNS2 AAVs carrying the genes within Cre-dependent constructs (0.1 mL of a 1:1 cocktail of MacPNS2-DIO-EF1α::SNAP-tag-PDGFR, 7×10^12^ vg/mL and MaCPNS2-DIO-hSyn::GCaMP6s, 7×10^12^ vg/mL, Methods). MaCPNS2 was chosen for its efficient and preferential targeting of peripheral neurons as compared to a parent serotype AAV9 ^53^. To assess the role of MND targeting to specific neurons we additionally performed imaging experiments in Cre-negative littermate controls (Oxtr-Cre^−/–^). These mice were retro-orbitally injected with MaCPNS2-hSyn-GCaMP6s (0.1 mL, 7×10^12^ vg/mL) to achieve broad expression of GCaMP6s in all peripheral neurons (including those in NGs). Following a 3-week incubation period to allow for transgene expression, BG- and Cy7-functionalized MNDs (0.5 μL, 1 mg/mL) were injected into the left NG in both groups of mice. Robust colocalization between the MNDs (Cy7), SNAP-tag (SNAP dye), and GCaMP6s was observed in Oxtr-Cre^+/–^ mice (**Fig. 3b**), while minimal and punctal Cy7 fluorescence indicative of microscopic MND precipitates was observed in Cre-negative controls (**Fig. 3c**).

Oxtr-Cre^+/–^ mice (and Oxtr-Cre^−/–^ controls) expressing GCaMP6s in NG neurons and injected with MNDs in the left NG and were then anesthetized (1% isoflurane) and exposed to four 4 s epochs and four 10 s epochs of MF stimulation (80 mT, 50 Hz, separated by 30 s rest epochs) during dynamic Ca^2+^ imaging of their left NGs via a custom-built setup (**Fig. 3a**, Methods). Robust and time-locked (latency 500±35 ms for 4s epochs and 650±41 ms for 10s epochs) GCaMP6s fluorescence transients in response to MF epochs were observed in Oxtr-Cre^+/–^ mice (**Fig. 3d,e**, Supplementary Video S3). In contrast, in control Oxtr-Cre^−/–^ mice MF stimulation did not trigger detectable synchronized Ca^2+^ responses (**Fig. 3f**), suggesting that neuronal activation in the NGs was dependent on MND targeting to neurons.

### Magnetomechanical stimulation activates vagal gut-brain satiety pathways in vivo

Following our observation of MF-evoked activity in mechanosensory Oxtr^+^ NG neurons targeted with MNDs, we hypothesized that magnetomechanical activation can be applied to vagal gut-brain pathways that mediate satiety (**Fig. 4a**).

**Fig. 4.**
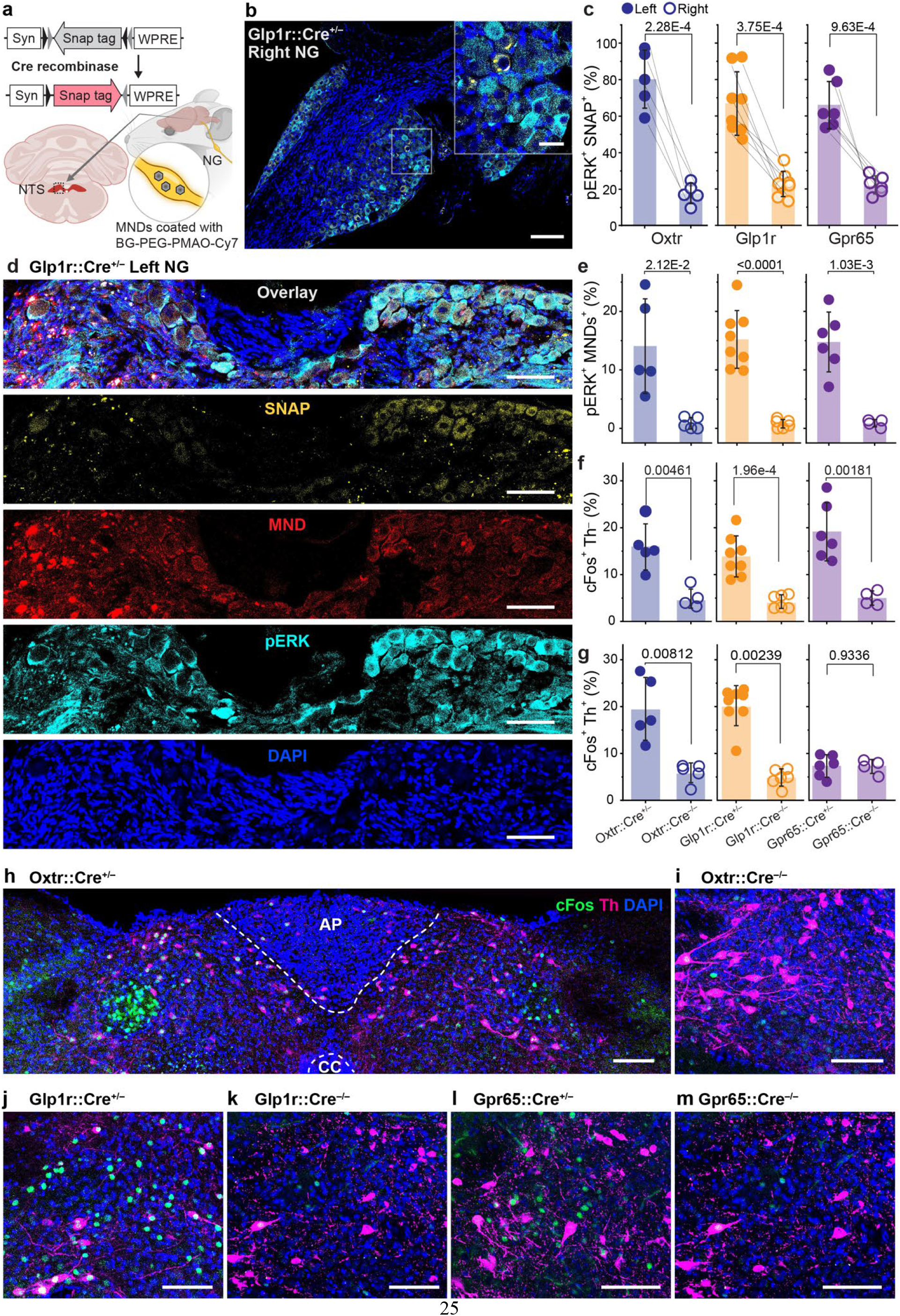
MND-mediated activation of nodose neurons and downstream hindbrain structures. **a**, Schematic illustration of the left and right NGs and their sensory projections to the caudal nucleus tractus solitarius (NTS) in the brainstem. **b**, Immunofluorescence image of the *right* NG in a Glp1r-Cre^+^/^−^ mouse (control side without MND injection). The scale bar is 100 μm. The scale bar of inset is 25 μm. **c**, Quantification of NG cells with overlapping expression of SNAP-tag (yellow) and phosphorylated ERK (pERK, cyan), indicating cell-specific activation; blue indicates DAPI-labeled nuclei, and red shows Cy7 fluorescence marking MND localization. As the distribution follows normality, a paired sample t-test was performed; Oxtr-Cre^+^/^−^ t=12.61, DF=4; Glp1r-Cre^+^/^−^ t=6.377, DF=7; Gpr65-Cre^+^/^−^ t=6.93, DF=5. **d**, Immunofluorescence image of the *left* NG from the same animal, injected with SNAP-targeted MNDs and exposed to MF (80 mT, 50 Hz). Scale bar is 50 μm. **e**, Quantification of NG cells co-expressing pERK and Cy7 (MNDs), confirming selective activation of MND-bound neurons. As the distribution follows normality, a two-sample t-test was performed; Oxtr line t=3.62, DF=4.12; Glp1r t=8.14, DF=7.40; Gpr65 line t=6.57, DF=5.22. **f**, Proportion of c-Fos–positive, tyrosine hydroxylase (TH)–negative cells relative to total DAPI-stained cells. For the data following a normal distribution, a two-sample t-test was performed; Oxtr line t=4.36, DF=4.78; Gpr65 line t=0.086, DF=7.99. For the data not following a normal distribution, a Mann-Whitney U-test was performed; Glp1r line U=48, Z=3.04. **g**, Proportion of c-Fos–positive, TH–negative cells relative to total DAPI-stained cells. For the data following a normal distribution, a two-sample t-test was performed; Oxtr line t=4.65, DF=5.4; Gpr65 line t=5.38, DF=5.88. For the data not following a normal distribution, a Mann-Whitney U-test was performed; Glp1r line U=48, Z=3.04. **c-g** Oxtr-Cre^+^/^−^ n_female_=2, n_male_=3; Oxtr-Cre^−^/^−^ n_female_=3, n_male_=2; Glp1r-Cre^+^/^−^ n_female_=4, n_male_=4; Glp1r-Cre^+^/^−^ n_female_=3, n_male_=3; Gpr65-Cre^+^/^−^ n_female_=3, n_male_=3; Gpr65-Cre^+^/^−^ n_female_=2, n_male_=2. **h-m**, Immunofluorescence images of NTS sections from: (**h**) Oxtr-Cre^+^/^−^ ; (**i**) Oxtr-Cre^−^/^−^; (**j**) Glp1r-Cre^+^/^−^; (**k**) Glp1r-Cre^−^/^−^; (**l**) Gpr65-Cre^+^/^−^; and (**m**) Gpr65-Cre^−^/^−^ mice. Green, c-Fos; magenta, Th; blue, DAPI. Scale bars, 100 μm.

The bifurcated axons of NG neurons simultaneously project to the brain stem and the visceral organs ^50,51^. In the gut, these vagal afferent form specialized endings that detect either chemical (mucosal endings) or mechanical (intraganglionic laminar endings, IGLEs) stimuli ^54^. These mechanosensory and chemosensory circuits that help orchestrate food consumption have been recently shown to have distinct genetic identities and anatomical distributions. For instance, Oxtr+ IGLEs densely innervate the small intestine ^52^, while mechanosensory glucagon-like peptide 1 receptor (Glp1r) expressing IGLEs are enriched in the stomach, and chemosensory G-protein coupled receptor 65 (Gpr65) vagal afferents have mucosal endings primarily in the duodenum ^52^.

We sought to evaluate the ability of MND-mediated stimulation to selectively activate these genetically and functionally distinct gut-brain vagal circuits. Akin to the imaging experiments described above, we employed Oxtr-Cre^+/–^ (n=10) as well as Glp1r-Cre^+/–^ (n=8, heterozygous) and Gpr65-Cre^+/–^ (n=6, heterozygous) mice to achieve cell-type specific expression of SNAP-tag protein in NG neurons. The construct for SNAP-tag expression was systemically delivered in a Cre-dependent construct (DIO-EF1α::SNAP-tag-PDGFR) via MacPNS2 AAV (retro-orbital route, 0.1 mL, 7×10^12^ vg/mL, **Fig. 4a**, Methods). Cre-negative littermates were employed as controls (Oxtr-Cre^−/–^, n=8; Glp1r-Cre^−/–^, n=6; Gpr65-Cre^−/–^, n=4). Following a 3-week incubation period BG- and Cy7-functionalized MNDs (0.5 μL, 1 mg/mL) were injected into the left NGs due to the their stronger relevance to feeding regulation than the right vagal circuits that additionally relay food reward signals ^55,56^.

To assess whether MND-mediated stimulation selectively activates targeted vagal circuits, we quantified the activity of the left NG neurons (right NG neurons serving as controls) as well the activity of neurons in the caudal nucleus tractus solitarius (NTS), a brainstem region critical for control of satiety that receives direct vagal sensory inputs and ^57,58^ (**Fig. 4a**). All mice were exposed to MF stimulation (80 mT, 50 Hz; 5 s epochs separated by 30 s rests for 30 min), and immunofluorescence analysis was employed to quantify expression of immediate early genes linked to neuronal activation, phosphorylated extracellular signal-regulated kinase 1/2 (pERK) in the NGs and c-Fos in the NTS ^59-61^.

In the left NGs injected with functionalized MNDs in all three types of mice, robust pERK expression with substantial SNAP-tag co-localization was observed (**Fig. 4b**,**c**, and **Fig. S17-19**). In contrast, only baseline pERK expression and significantly lower co-localization with SNAP-tag was observed in the right (control) NGs devoid of MNDs (**Fig. 4d** and **Fig. S20-22**). Similarly, in Cre-negative controls, where SNAP-tag was not expressed (**Fig. S23-25**), MNDs did not target NG neurons, and significantly lower pERK activation was observed in response to MF. Quantitative analysis of the overlap between MNDs (visualized via Cy7 labeling) and pERK expression confirmed significantly higher activation of left NG neurons in Cre-positive mice as compared to Cre-negative controls (**Fig. 4e**). This is consistent with the imaging experiments that revealed that MND targeting was a prerequisite for MF-triggered neuronal activity in NGs.

We then examined downstream activation in the NTS, as marked by c-Fos expression and its overlap with tyrosine hydroxylase (Th), a marker of catecholaminergic neurons projecting to hypothalamus and parabrachial nucleus (PBN) that contribute to regulation of hunger and feeding ^62-65^. MND-mediated MF stimulation of Oxtr+ and Glp1r+ left NG neurons yielded robust c-Fos expression in Th+ neurons in the NTS, as compared to stimulation of Gpr65+ left NG neurons or non-specific stimulation (MNDs injected in left NGs of Cre-negative mice without SNAP-tag expression) (**Fig. 4f-m** and **Fig. S26-31**). This is consistent with prior observations of enhanced c-Fos expression in a subset of Th+ NTS neurons following selective activation of Oxtr+ and Glp1r+ vagal afferents (**Fig. 4f,g**) ^52^.

Notably, both Th^+^ and Th^−^ cells exhibited increased c-Fos expression in Oxtr-Cre^+/–^ and Glp1r-Cre^+/–^ mice following stimulation mediated by targeted MNDs in the left NGs (**Fig. 4f-m**). This suggests activation of divergent downstream populations either through direct vagal inputs or through second-order activation of neurons projecting to Th^+^ populations. While NTS neurons activated by Glp1r^+^ vagal inputs are largely distinct from Th^+^ populations neurons, some overlap has been reported ^66^. Similarly, Oxtr^+^ vagal afferents also modulate activity of non-catecholaminergic (Th^−^) neurons in the NTS ^52,67^.

In contrast, MND-mediated stimulation of Gpr65^+^ left NG neurons led to increased c-Fos expression exclusively in Th^−^ neurons, with no significant increase in activation of Th+ neurons (**Fig. 4f,g**). Gpr65^+^ vagal afferents detect intestinal nutrient signals and primarily project to the commissural subnucleus of the NTS, which is anatomically distinct from Th+ neuron-enriched regions. Accordingly, MND-mediated stimulation of Gpr65^+^ left NG neurons did not activate Th^+^ neurons that regulate satiety. Instead, the elevated c-Fos in Th^−^ neurons suggests recruitment of a functionally distinct NTS circuit, likely involved in metabolic regulation rather than feeding suppression ^68,69^.

These findings suggest that magnetic stimulation via MNDs enables selective activation of targeted genetically and functionally distinct vagal gut-brain pathways with an added benefit of anatomical restriction (e.g. left vs. right NGs) afforded by the choice of the MND injection site.

### Magnetomechanical activation of targeted gut-brain satiety pathways suppresses feeding

Selective optogenetic and chemogenic activation of Oxtr^+^ and Glp1r^+^ vagal afferents in the NTS was previously shown to suppress feeding in fasted mice, while activation of Gpr65^+^ afferents did not significantly influence food intake ^52^. We examined whether direct activation of the neuron cell bodies in the left NGs via targeted MNDs would offer a precise yet wireless tool to manipulate feeding behaviors by engaging the vagal pathways involved in gut-brain communication (**Fig. 5a**).

**Fig. 5.**
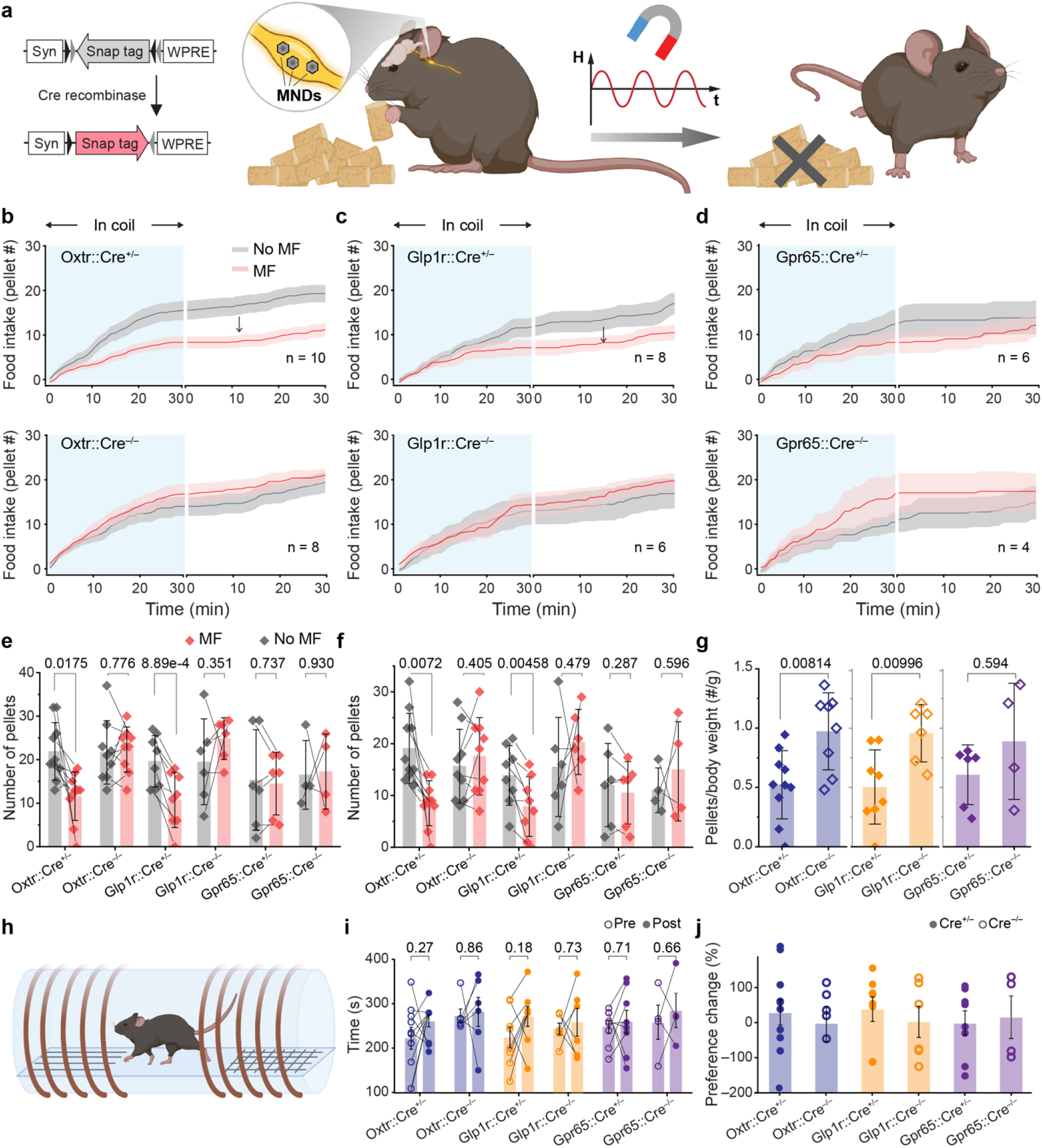
Wireless magnetic control of feeding behavior mediated by targeted MNDs. **a**, An illustration of feeding assays in mice expressing Cre-recombinase in specific vagal mechanoreceptors targeted with SNAP-tag and injected with MNDs in the left NGs. **b-d**, Time-course of food intake recorded using FED3.0 during magnetic field (MF) stimulation (blue shaded area; 80 mT, 50 Hz; 5 s pulses every 25 s for 30 min) followed by 30 min without stimulation in: (**b**) Oxtr-Cre^+^/^−^ (n_female_=5, n_male_=5) and Cre^−^/^−^ (n_female_=4, n_male_=4) mice; (**c**) Glp1r-Cre^+^/^−^ (n_female_=4, n_male_=4) and Cre^−^/^−^ (n_female_=3, n_male_=3) mice; (**d**) Gpr65-Cre^+^/^−^ (n_female_=3, n_male_=3) and Cre^−^/^−^ (n_female_=2, n_male_=2) mice. Lines and shaded areas represent mean ±s.e.m. (**e**) Total number of food pellets consumed over the 60-min session (30 min with MF + 30 min post-stimulation). As the data distribution follows normality, a paired sample t-test was performed; Oxtr-Cre^+^/^−^ t=2.90, DF=9;Glp1r-Cre^+^/^−^ t=5.52, DF=7; Gpr65-Cre^+^/^−^ t=–0.355, DF=5; Oxtr-Cre^−^/^−^ t=-0.29, DF=8; Glp1r-Cre^−^/^−^ t=-1.03, DF=5; Gpr65-Cre^−^/^−^ t= -0.096, DF=3. **f**, Number of pellets consumed during the 30-min MF stimulation period only. As the data distribution follows normality, a paired sample t-test was performed; Oxtr-Cre^+^/^−^ t=3.47, DF=9; Glp1r-Cre^+^/^−^ t=4.10, DF=7; Gpr65-Cre^+^/^−^ t=1.19, DF=5; Oxtr-Cre^−^/^−^ t=–0.88, DF=8; Glp1r-Cre^−^/^−^ t=-0.76, DF=5; Gpr65-Cre^−^/^−^ t=-0.59, DF=3. **g**, Food intake over 60 min, normalized to body weight, comparing Cre^+^/^−^ and Cre^−^/^−^ mice across Oxtr, Glp1r, and Gpr65 lines. As the distribution of Oxtr and Glp1r lines follows the normality, a two-sample t-test was performed; Oxtr line t=-3.07, DF = 14.2; Glp1r line t=–3.06, DF=12.0. For the Gpr65 line, whose intake data does not follow a normal distribution, the Mann-Whitney U test was performed; U=9, Z = –0.533. Bars and error bars represent mean ±standard deviation (s.d.) **h**, Schematic of place preference assay, where mice receive MF stimulation only upon entering the less-preferred chamber; baseline (pre) and test phases (post) are performed without MF. **i**, Time spent in the stimulation chamber before and after three days of training with closed-loop MF stimulation. **j**, Percent change in time spent in the MF stimulation chamber. Bars and error bars represent mean ±s.d.

Akin to the immunohistology analyses described above (Fig. 4), Oxtr-Cre^+/–^ (n=10), Glp1r-Cre^+/–^ (n=8), and Gpr65-Cre^+/–^ (n=6) mice (and Cre-negative controls: Oxtr-Cre^−/–^, n=8; Glp1r-Cre^−/–^, n=6; Gpr65-Cre^−/–^, n=4) were systemically transduced to express SNAP-tag in targeted neurons, and then injected with functionalized MNDs in their left NGs after a 3-week incubation period. Following post-surgical recovery, all mice have undergone food intake assays using a custom apparatus equipped with an electromagnet and a FED3.0 device (**Fig. 5a, Fig. S32**). In these assays, mice habituated to the apparatus, were fasted for 18-21 hrs during the active phase (Methods) and then given access to chow pellets for 30 min while being exposed to MF (80 mT, 50 Hz; 5 s epochs separated by 30 s rests; 30 min). The mice were then allowed to feed for an additional 30 min without MF application in another FED3.0-equipped arena.

Consistent with prior observations, we found significantly reduced food intake during MF stimulation in Oxtr-Cre^+/–^ and Glp1r-Cre^+/–^ mice (**Fig. 5b,c**). During the post-stimulation period, diminished food intake was observed in the Glp1r-Cre^+/–^, whereas in Oxtr-Cre^+/–^ food intake was comparable to non-stimulation conditions with no observable rebound feeding. In contrast, Gpr65-Cre^+/–^ mice exhibited no change in food intake during or after MF stimulation, suggesting that mucosal afferents have limited involvement in the control of satiety (**Fig. 5d, S33**). Cre-negative littermates injected with MNDs, but lacking SNAP-tag expression showed no significant changes in feeding behavior upon MF stimulation (**Fig. 5b-f, S33**). Notably, even when controlling for baseline food intake and body mass differences, stimulation of Oxtr^+^ and Glp1r^+^ but not Gpr65^+^ left NG neurons significantly suppressed feeding as compared to MF exposure in Cre-negative litter mates (**Fig. 5g**).

To rule out off-target effects such as malaise or stress, we conducted a place preference/ aversion assay, during which the mice that have previously undergone feeding assays explored a two-chamber arena externally equipped with electromagnets (**Fig. 5h, Fig. S34**, Methods). Following habituation when mice typically showed a bias toward one chamber of the arena, the mice received MF stimulation only when entering their less-preferred chamber. No significant changes in preference were observed for any group (Oxtr-Cre^+/–^, Glp1r-Cre^+/–^, Gpr65-Cre^+/–^, or Cre^−/–^ controls), suggesting that MND-mediated stimulation lacked detectable positive or negative valence (**Fig. 5h-j**).

These findings demonstrate the ability of targeted MNDs to mediate wireless magnetic control of genetically identifiable vagal gut-brain circuits involved in the regulation of satiety and suggest applications of this technology to studies of other vagal brain-body circuits.

## Conclusion

By leveraging magnetic nanodiscs (MNDs) targeted to specific cells via genetically encoded SNAP-tag moieties, this study establishes a wireless magnetomechanical paradigm for controlling vagal gut-brain pathways. Our approach affords cell-type specificity, while delivering sub-second temporal precision inaccessible with chemogenetics or pharmacology and eliminating the need for implantable devices employed for optogenetics or electrical neuromodulation. Selective magnetic stimulation of mechanosensory Oxtr^+^ and Glp1r^+^ neurons in the left nodose ganglia activated vagal gut-brain pathways previously demonstrated to drive satiety and suppressed feeding without inducing negative valence. Peripheral ganglia are critical relays of communication between the central and peripheral neural circuits, yet their miniature dimensions and often challenging anatomical locations make them inaccessible with chronically implantable neuromodulation devices. By demonstrating a targeted and implant-free peripheral neuromodulation, this work empowers studies of brain-body communication and informs the development of future injectable bioelectronic medicines.

## Acknowledgments

The authors thank Dr. Michael Christiansen at the Swiss Federal Institute of Technology (ETH Zurich) for insightful discussions and facilitating collaboration in simulation of mechanical forces generated by the nanodiscs. This work was funded in part by the Director’s Pioneer Award from the National Institutes of Health and National Center for Complementary and Integrative Health (DP1-AT011991), K. Lisa Yang and Hock E. Tan Center for Molecular Therapeutics at MIT, K. Lisa Yang Brain-Body Center at MIT, and the McGovern Institute for Brain Research at MIT. Y.J.K. is a recipient of the Mathworks Fellowship. N.B. is a recipient of the Human Frontier Science Program: Long-term Postdoctoral Fellowship. K.N. is a recipient of the Honjo International Scholarship Foundation and Eva Tan Fellowship from the K. Lisa Yang and Hock E. Tan Center for Molecular Therapeutics at MIT. E.W. is a recipient of the NIH Neurobiological Engineering Training Program Grant. J.L.B. is a recipient of the Schmidt Science Fellows program, in partnership with the Rhodes Trust. K.P. is a recipient of the Friends of the McGovern Institute Student Fellow, Irene T. Cheng Fellowship, and Henry E. Singleton Fellowship. T.M.C. is a recipient of the NIH NRSA F32 postdoctoral fellowship (1F32MH139162-01).

## Author contributions

Y.J.K. and P.A. designed the study. YJ.K. synthesized and characterized the materials, performed all in vitro and in vivo experiments. Y.J.K. and F.K. developed targeting approach and magnetic instrumentation. Y.J.K. and N.B. designed and executed food intake behavior assays. Y.J.K. and C.F. performed computational simulations for mechanical force generated by MNDs. Y.J.K. and T.M.C. performed calcium imaging on nodose ganglia in vivo. Y.J.K., E.V.P., R. L. and K.N. prepared plasmids and vectors for SNAP-tag expression. Y.J.K and J.S. developed behavior analyses tools. S.M. conducted mouse colony management, breeding, and phenotyping. Y.J.K. and K.P. developed protocols for surgery and extraction of mouse nodose ganglia. Y.J.K. and J.L.B. performed FTIR analysis on polymer coatings. Y.J.K. and E.W. simulated the MFs within electromagnets used for calcium imaging in vivo. Y.J.K., N.B., and R.L. prepared figures. All authors contributed to writing of the manuscript.

## Supplementary Materials

Materials and Methods

Supplementary Notes 1 and 2

Figs. S1 to S34

References (*1-11*)

Supplementary Videos S1 to S3

