## Supplementary information for "Targeted Magnetic Nanodiscs for Wireless Causal Manipulation of Gut-Brain Circuits"

**The PDF file includes:**

Materials and Methods
Supplementary Notes 1 and 2
Figs. S1 to S34
References 1-11

**Other Supplementary Materials for this manuscript include the following:**

Supplementary Videos S1 to S3

### Materials and Methods

Synthesis and Characterization of BG-PEG-PMAO-Cy7 Coated Magnetite Nanodiscs (MNDs)

Hematite nanodiscs were first synthesized by heating a homogeneous mixture of 0.273 g $FeCl_3 \cdot 6H_2O$ (Fluka), 0.8 g anhydrous sodium acetate (Sigma-Aldrich), 10 mL ethanol, and 0.8 mL deionized (DI) water in a sealed Teflon-lined steel autoclave at 180 °C for 18 hours. The resulting red hematite nanodiscs were washed 3–5 times with ethanol and DI water, then redispersed in 20 mL trioctylamine (Sigma-Aldrich) and 1 g oleic acid (Alfa Aesar).

Reduction of hematite to magnetite was conducted under a 5% $H_2$ / 95% $N_2$ atmosphere in a three-neck flask connected to a Schlenk line. The system was evacuated at room temperature for 20 minutes, then heated to 370 °C at 20 °C/min for 30 minutes to yield hydrophobic oleic acid-coated MNDs.

For SNAP-tag targeting, the MNDs were rendered hydrophilic by coating with a functionalized PMAO-based polymer. First, 8.75 mg of poly (maleic anhydride-alt-1-octadecene) (PMAO, Mn 30,000–50,000, Aldrich) was mixed with 0.2 mg $NH_2$-Cy7 (Lumiprobe) in 2.5 mL tetrahydrofuran (THF) overnight. After solvent evaporation, the polymer was redissolved in 2 mL chloroform and incubated with 2 mg BG-PEG-$NH_2$ (New England Biolabs) for 8 hours.

Next, 1 g of MNDs was added to the polymer solution, followed by 10 minutes of sonication. Chloroform was evaporated overnight, and the coated MNDs were washed 3–5 times in 5 mL TEA buffer and twice with 5 mL DI water.

Polymer composition was confirmed via Fourier Transform Infrared Spectroscopy (FTIR) using a ThermoFisher FTIR6700. Morphology was examined by transmission electron microscopy (TEM; FEI Tecnai G2 Spirit TWIN), and disc dimensions (diameter and thickness) were measured from TEM images. Room-temperature hysteresis loops were obtained using a vibrating sample magnetometer (VSM) on a Quantum Design MPMS-3 system. Elemental analysis for saturation magnetization was performed using an Agilent 5100 ICP-OES. Samples for ICP-OES were digested overnight in 37% HCl and diluted in 2 wt% $HNO_3$.

Design and Fabrication of Electromagnets

A C-shaped magnetic core (1018 steel) was wound with 14 AWG copper wire (TEMCo) to generate a uniform magnetic field across an 8 mm gap. Coverslips containing cultured rat hippocampal neurons were placed in the field center under an inverted microscope. For food intake assays, 11 AWG copper wire was wound around an acrylic cylinder (8 cm diameter, 5 cm height, 2 mm wall thickness) and mounted to the FED3.0 system.

An acrylic cylinder (75 mm inner diameter, 2 mm wall thickness) was divided into three chambers: a 10 cm neutral middle chamber with white flooring and two 11 cm stimulation chambers with stripe- and check-patterned floors. Each stimulation chamber was wound with 11 AWG copper wire.

The coil used for in vivo nodose ganglia (NG) imaging had a 12 mm diameter central core made of 1018 steel with a conical upper part (12 mm top, 18.3 mm base diameter, 4 mm height) and a cylindrical lower part (18.3 mm diameter, 45.75 mm height). The coil was wound with 19 AWG copper wire, and the assembly was encased in 1018 steel (6.4 mm thick base and 12.9 mm thick side walls).

To generate magnetic fields (MF; 20–320 mT, 10–200 Hz), the coils were connected to a Crown DC-300A Series II amplifier and a Picoscope 2204A signal generator.

<u>In-vitro Calcium imaging in cultured hippocampal neurons</u>

All animal procedures were approved by the Massachusetts Institute of Technology Committees on Animal Care (Protocol #2506000814). The neonatal rat hippocampal neurons were extracted from Sprague-Dawley, 001 pups (P1) and dissociated with Papin (Worthington Biochemical). On glass slides (5 mm diameter, Bellco Glass 1943-00005), the neurons were seeded and incubated in 24-well plates at a density of 112,500 cells $ml^{-1}$. The glass slides were sterilized by evaporating ethanol with an alcohol lamp and then coated with Matrigel (Corning). In each well, 3-5 cover slips were incubated in 1 ml Neurobasal medium (Invitrogen), and 3 days after seeding, 5-fluoro-2′-deoxyuridine (F0503 Sigma) was treated for glial inhibition. Four days after seeding, the neurons were transduced with 1 µl of AAV9-hSyn::GCaMP6s (Addgene viral prep #100843-AAV9, >1 × $10^{13}$ IU $ml^{-1}$) or AAV9−EF1α::SNAP-tag−PDGFR (3 × $10^{13}$ vg $ml^{-1}$)

The neurons were incubated for 1 h in 37 °C MND solution dispersed in Tyrode solution to allow the BG on the surface of MNDs to precipitate onto the cells expressing SNAP tags. The concentration and volume of the MND incubation solution were controlled for the experiment, with variation in MND density on neurons. For Hoechst staining, Thermo Scientific, Hoechst 33342 Solution (20 mM) was added to the incubation solution 20 minutes prior to imaging. After the incubation of neurons in the MND solution, they were transferred to the calcium imaging holder containing 200 µl Tyrode's solution. This holder is placed in the gap of the C-shaped coil described above. GCaMP6s fluorescence was recorded at 1 fps for phase diagram experiments or 20 fps for others, using a fluorescent microscope (Olympus IX73, 20× objective lens). Fluorescence recordings were analyzed using ImageJ. Baseline ($F_0$) was defined as the average intensity 20 s before initial MF onset. From the $\Delta F/F_0$ of each cell, the cells were defined as responsive when the $\Delta F/F_0$ value exceeded three times the standard deviation of the baseline. Each experiment was repeated on three independent coverslips. Peak response latency was determined using MATLAB's peakfinder function.

<u>Surgeries</u>

All surgical procedures were approved by the MIT Committee on Animal Care (Protocol #2506000815). Three weeks following retro-orbital injection (*1*) of MaCPNS2 AAVs in 6–8-week-old mice, they underwent aseptic surgery for MND injection into the nodose ganglia (NGs). Mice were anaesthetized with isoflurane (0.5-2.5% in $O_2$) using an anaesthesia machine (VET EQUIP) with their eyes covered with ophthalmic ointment and a heat pad to maintain the animals' body temperature. Subcutaneous injections of 0.5 mL of sterile Ringer's solution for hydration and extended-release buprenorphine (1 mg $kg^{-1}$) for analgesia were provided to the mice at the start of the surgery. To expose the NG, animals were fixed in a supine position using tape and the hair removal cream was applied from the chin to the thorax and then wiped with saline. Then, the surgical area was disinfected with povidone-iodine and 70% ethanol, each applied three times. Cutting a midline incision exposed the submandibular glands, and then the outwards bilateral retraction of them exposed the trachea. Subsequently, the sternocleidomastoid muscle was retracted laterally, and the omohyoid muscle was retracted towards the midline to expose the carotid sheath. This then grants visibility of the vagus trunk next to the carotid artery, and the NGs correspond to the crossings with the hypoglossal nerve immediately adjacent to the base of the skull. The NG was carefully detached from the hypoglossal nerve. A glass pipette connected to NanojectIII (Drummond 3-000-207) was loaded with 1 mg $mL^{-1}$ MNDs solution, and 500 nL of the MND solution was injected into the NG. At the end of surgery, the skin incision was sutured and post-operative analgesia, carprofen 5 mg $kg^{-1}$was applied. The animals were transferred to a clean cage, part of which was positioned on a heating pad with ad libitum access to water and diet gel (Bio-serve NutraGel).

Behavior assays

All behavioral assays were approved by the MIT Committee on Animal Care (Protocol #2506000815). For behavioral experiments, the mice were tested during the dark phase of the 12 h-reverse light/dark cycle.

Feeding experiments were performed using a pellet dispending system (FED3.0(*2*)). Food pellets (ScottPharma, 5TUM) were dispensed at the beginning of trials or after pellet removal with intervals shorter than 3 s. Prior to start of experiments, mice were habituated for three days to the pellet dispending system (3 h per day) and also together with the magnetic coil attached to the device (30 min per day). Prior to the test, mice were fasted for overnight (18-21 h). The MF application was performed for the initial 30 min of the refeeding duration.

The place preference experiment was performed in the stimulation chambers equipped with custom-made electromagnets described in the previous section. The daily 15 min habituation for the setup was performed for 3 days prior to the initial stimulation on the less preferred chamber, and during the following 3 days, the MF (80 mT 50 Hz; 5s duration 25 intervals) was applied daily for 10 min, when mice entered the less preferred area. At the end of the three habituation days and subsequential 3 days of training, the mice explored the entire arena without stimulation (post), which was compared with the baseline preference (pre) determined by the average of the last two habituation days. The mouse location in the arena was recorded with Logitech HD Pro Webcam C920, and analysis of the place preference assays was done with a custom-made pipeline (*3*).

Immunohistochemical quantification of biomarkers

Following the stimulation during the behavior experiments, after a 90-minute c-Fos induction period, the mice were sacrificed by a lethal intraperitoneal injection of sodium pentobarbital (Fatal-plus, 50 mg $ml^{-1}$, dose 100 mg $kg^{-1}$). Transcardial perfusion was then performed with PBS and 4% paraformaldehyde (PFA) solution, postfixed in 4% PFA overnight. The NGs and brain were isolated from the skull and the NGs and brains were stored in PBS. Brain tissue was sectioned into 50 μm coronal slices using a vibrating blade microtome (Leica VT1000S), and the NGs were cryoprotected with 30% sucrose in PBS overnight at 4°C, embedded in OCT, frozen and stored at −20°C and then sectioned at 20 um thickness using a cryostat (Leica CM3050S).

The NG slices were washed 3x10 min with PBS, incubated with 5 μM SNAP-Surface® Alexa Fluor® 647 (NEB #S9136) for 30 min at room temperature, washed 10 min with PBS, washed 2x10 min with 0.3% v/v Triton X-100 solution in PBS, blocked for 1 h with blocking solution (0.3% v/v Triton X-100 and normal donkey serum 5% v/v blocking serum solution in PBS), incubated with primary antibodies (1:500 diluted in blocking solution) overnight at 4°C, washed 3x10 min with blocking solution, and then incubated with secondary antibodies (1:1000 diluted in blocking solution) for 2 hours at room temperature. After 3x10 min washing with PBS, the slices were incubated in 1:20000 4′,6-diamidino-2-phenylindole (DAPI) solution for 30 min, washed 3x10 min with PBS, and then covered with mounting medium (Fluoromount G, Southern Biotech) and cover glass.

The brain slices were incubated 3 × 10 min with 0.3% v/v Triton X-100 in PBS, blocked 3% NDS in 0.3% PBST for 60 min at RT, and incubated with primary antibodies (1:1000 diluted in blocking solution) overnight at 4°C, washed 3x10 min with PBS, incubated with secondary antibodies (1:1000 diluted in blocking solution) for 2 h at RT, washed again 3 × 10 min with PBS, and then stained 1:20000 4′,6-diamidino-2-phenylindole (DAPI) solution for 30 min. After final washing with PBS, the brain slices were mounted with the mounting medium.

For tyrosine hydroxylase (TH) staining, sheep-anti-tyrosine hydroxylase antibody (Novus Biologicals NB300-110SS) and donkey anti-sheep IgG (H + L), Alexa Fluor 568 (Thermo Scientific A-21099) were used as primary and secondary antibodies. Rabbit anti-c-Fos (Cell Signaling Technology, 2250s) primary antibodies and donkey anti-rabbit Alexa Fluor 488 (Invitrogen, A-21206) secondary antibodies were used for the c-Fos expression analysis. Phospho-p44/42 MAPK (Erk1/2) (Thr202/Tyr204) (Cell Signaling Technology, 9101s) primary antibodies and donkey anti-rabbit Alexa Fluor 488 (Invitrogen, A-21206) secondary antibodies were used for pERK expression analysis.

<u>*In vivo* $Ca^{2+}$ imaging in nodose ganglia</u>

*In vivo* NG $Ca^{2+}$ imaging with GCaMP6s was conducted under isoflurane (0.5-2.5% in $O_2$) anesthesia following a NG exposure according to the procedure described in the surgery section above. After NG exposure, imaging was conducted using a FLIR Blackfly monochrome camera

(BFS-U3-200S6M-C USB 3.1) with a frame rate of 5 Hz. To enable the MF application, the entire procedure was performed on top of the electromagnet, such that the mouse neck region was positioned in the center of the coil (Fig. S32).

Statistical Analyses

OriginPro 2025b was used for assessing the statistical significance of all comparisons. Although sample sizes were not determined with power analysis, the group size for behavior tests and immunohistochemistry was determined to be similar to previous research associated with similar neural circuits. To test the normality of data distribution, the Shapiro-Wilk test was performed. For the data following normal distribution, statistical significance among groups of three or more was analyzed with analysis of variance (ANOVA) followed by Tukey's post-hoc comparison test. For non-normally distributed data, comparisons among three or more groups were performed using a Kruskal–Wallis test followed by Dunn's post hoc test. To compare independent groups with normally distributed data, a two-sample t-test was performed, while Mann-Whitney U-test was performed for the groups not following a normal distribution. For the dependent groups, a paired t-test was done for the data following a normal distribution, or a Wilcoxon signed-rank test was performed for the non-normally distributed data. P-values are not indicated on plots for inter-group comparisons where no significant differences were found.

### Supplementary Text

#### Supplementary Note 1

#### 1.1 Dynamic Model

The dynamic model describes changes in magnetization direction driven by physical particle rotation and internal magnetization processes, allowing for estimation of torques applied by the magnetic nanodisc (MND) particle to its surroundings. The MND rotational inertia is assumed to be negligible compared to the magnetic and fluidic torques. This assumption is reasonable for nanoparticles due to the $1/L^5$ (where $L$ is a particle linear dimension) scaling of rotational inertia. Therefore, the dynamic equation of a single magnetic nanoparticle constrained to rotate about a fixed axis is:

$$\mathbf{0} = \boldsymbol{T}_m + \boldsymbol{T}_f + \boldsymbol{T}_b \qquad \text{eq.1}$$

Where $\boldsymbol{T}_m$ is the magnetic torque applied to the particle in the presence of an external magnetic field, $\boldsymbol{T}_f$ is the hydrodynamic drag force acting on the particle, and $\boldsymbol{T}_b$ is a torque that accounts for the Brownian motion of the particle. The fluidic torque on the MND can be expressed as:

$$\boldsymbol{T}_f = -\eta\Omega_O\boldsymbol{\omega_r} \qquad \text{eq.2}$$

where $\eta$ is the dynamic viscosity of the fluid, $\Omega_O$ is the hydrodynamic rotation tensor about a point $O$ that lies on an instantaneous fixed axis of rotation and is rigidly attached to the particle, and $\boldsymbol{\omega_r}$ is the instantaneous angular velocity of the particle about the instantaneous axis of rotation (*4*). After substituting Eq. (2) into Eq. (1), the instantaneous angular velocity of an MND can be expressed as:

$$\boldsymbol{\omega_r} = \eta\Omega_O^{-1}\,(\boldsymbol{T}_m + \boldsymbol{T}_b) \qquad \text{eq.3}$$

This angular velocity describes the rate of physical rotation of the nanoparticle.

Assuming magnetic gradients produce negligible force couples on the MND about the point $O$, the applied torque on the particle due to a magnetic field, $\boldsymbol{B}_a$, is

$$\boldsymbol{T}_m = \boldsymbol{m} \times \boldsymbol{B}_a \qquad \text{eq.4}$$

where $\boldsymbol{m}$ is the MND magnetic dipole moment. We assume that the externally applied flux density $\boldsymbol{B}_a$ is uniform over the volume of the nanoparticle. The MND dipole moment is also a function of the applied field:

$$\boldsymbol{m} = V_m \boldsymbol{M}(\boldsymbol{H}_a) \qquad \text{eq.5}$$

where $V_m$ is the volume of magnetized material, $\boldsymbol{M}(\boldsymbol{H}_a)$ is the volume magnetization (dipole density) as a function of $\boldsymbol{H}_a = \boldsymbol{B}_a/\mu_0$, and $\mu_0 = 4\pi \times 10^{-7}$ T m A$^{-1}$ is the vacuum permeability.

**1.2 Magnetization Model**

For MNDs, the magnetization $\boldsymbol{M}$ varies stochastically and is not a constant value over very short time scales (e.g. ns). We will assume that the particle is effectively always saturated $\| \boldsymbol{M} \| = M_s$, so that the only variation is in the direction of magnetization $\boldsymbol{w}$, where $\boldsymbol{M} = M_S \boldsymbol{w}$. Let this stochastic process be represented by the random variable $\boldsymbol{W}$. Observations of the random variable are distributed according to the following probability density:

$$\boldsymbol{W} \sim p(\boldsymbol{w}) = \frac{1}{C} \exp\left(-\frac{U(\boldsymbol{w})}{k_B T}\right) \qquad \text{eq.6}$$

where $U(\boldsymbol{w})$ is the energy of the particle (in Joules) as a function of $\boldsymbol{w}$, $k_B = 1.380649 \times 10^{-23}$ J K$^{-1}$ is the Boltzmann constant, $T$ is the temperature (in Kelvin), and $C$ is a unitless constant of proportionality needed to make the total probability equal to one.

Here, we consider three components of the particle energy: magnetostatic anisotropy energy $U_S$, magnetocrystalline anisotropy energy $U_C$, and Zeeman energy $U_Z$.

The magnetostatic (shape) anisotropy energy is

$$U_s = -\frac{\mu_0}{2} \int_V \boldsymbol{M} \cdot \boldsymbol{H}_d \, dV \qquad \text{eq.7}$$

where $\boldsymbol{H}_d$ is the demagnetizing field and $V$ is the magnetic volume of the particle (*5*). If we approximate the magnetic volume of the disc as an oblate ellipsoid with major radii $\rho_1 = \rho_2$ and minor radius $\rho_3 < \rho_1$, the demagnetizing field in the body frame is constant throughout the magnetic volume and can be determined using:

$$H_d = -\begin{bmatrix} n_{d1} & 0 & 0 \\ 0 & n_{d2} & 0 \\ 0 & 0 & n_{d3} \end{bmatrix} \boldsymbol{M} \qquad \text{eq.8}$$

where $n_{d1} = n_{d2}$ are the demagnetizing factors of the major axes of the ellipsoid and $n_{d3}$ is the demagnetizing factor of the minor axis of the ellipsoid. The demagnetizing factors of oblate spheroids can be directly calculated from elementary functions in Ref. (*6*):

$$n_{d3} = \frac{a^2}{a^2-1}\left(1 - \frac{1}{\sqrt{a^2-1}} arcsin\left(\frac{\sqrt{a^2-1}}{a}\right)\right) \qquad \text{eq.9}$$

where $a = \rho_1/\rho_3$ is the aspect ratio of the spheroid. The remaining demagnetization factors can be determined from $n_{d1} = n_{d2} = 0.5\,(1 - n_{d3})$.

Then the magnetostatic potential energy can be expressed as:

$$U_S = \frac{\mu_0 V M_{sat}^2}{2}(n_{d1}w_1^2 + n_{d2}w_2^2 + n_{d3}w_3^2) \qquad \text{eq.10}$$

Magnetocrystalline anisotropy energy $U_C$, for a nanoparticle material with cubic anisotropy such as magnetite, can be determined from the coordinate-dependent expression:

$$U_C(\boldsymbol{w}) = k_c V(w_1^2 w_2^2 + w_1^2 w_3^2 + w_2^2 w_3^2) \qquad \text{eq.11}$$

where the $w_i$ are the direction cosines of the magnetization vector along the $\langle 100 \rangle$ axes of the crystal and $k_C$ is the first-order cubic anisotropy coefficient (in J $m^{-3}$). The crystalline anisotropy energy landscape of a magnetite particle with negative cubic anisotropy coefficient $k_c = -9$ kJ $m^{-3}$ is shown in **Fig. S1** mapped onto the unit sphere of magnetization directions. Negative cubic anisotropy results in high energy along the $\langle 100 \rangle$ axes and low energy along the $\langle 111 \rangle$ axes. This anisotropy results in eight easy axes and six hard axes.

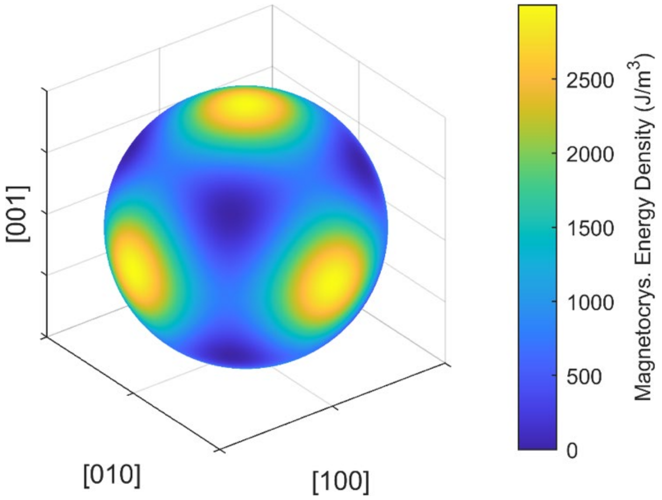


**Fig. SN1** Magnetocrystalline anisotropy energy per unit volume mapped onto the unit sphere of magnetization shown with the axes representing the ⟨100⟩ direction of the crystal structure.

The Zeeman energy $U_Z$ can be determined from

$$U_Z(\boldsymbol{w}) = -\int_V \boldsymbol{M} \cdot \boldsymbol{B}_a dV \qquad \text{eq.12}$$

which can be simplified for a uniformly magnetized particle in a uniform field to

$$U_Z(\boldsymbol{w}) = -\boldsymbol{m} \cdot \boldsymbol{B}_a = -M_S V(w_1 B_1 + w_2 B_2 + w_3 B_3) \qquad \text{eq.13}$$

Given the preceding model assumptions, the anisotropy energy landscape of MNDs under a MF contains multiple energy minima separated by energy barriers. Consequently, the probability density $p(\boldsymbol{w})$ in Eq. (6) is multimodal. We expect the magnetization to be trapped near each energy minimum with a finite likelihood of transition between the energy minima.

For example, in **Fig. SN2**, we simulated the magnetic energy landscape and relative likelihood of magnetization states of a hexagonal disc-shaped particle (230 nm in diameter and 30 nm thick - closely resembling our experimental MNDs) with 80 mT applied field along the short axis of the disc. The relative probability was calculated using Eq. (6) without the normalization term and letting the minimum energy equal to zero (the global minimum energy states have a relative probability of 1). **Fig. S2NA** shows the energy associated with magnetocrystalline anisotropy and **Fig. SN2B** shows the relative likelihood of a given magnetization state based on the magnetocrystalline anisotropy alone. The magnetocrystalline anisotropy energy results in eight sharp likelihood peaks (four visible in **Fig. SN2B**) that correspond to the corners of the cubic crystal structure (⟨111⟩ directions). Similarly, **Fig. SN2C** shows the energy associated with the shape anisotropy of the disc-shaped particle, and **Fig. SN2D** shows the corresponding relative likelihood. The shape anisotropy energy results in a sharp likelihood band around the equator (long axes) of the disc-shaped particle. **Fig. SN2E** shows the energy associated with the 80 mT applied field (the Zeeman energy), and **Fig. SN2F** shows the corresponding relative likelihood. The applied field biases the magnetization likelihood in its direction (here along the vertical axis). The sum of all energy contributions is shown in **Fig. S2G**, and the corresponding relative likelihood is shown in **Fig. SN2H**. The total energy landscape is dominated by the shape anisotropy energy, and the total energy density exhibits six distinct minima, which correspond to sharp maxima in the probability distribution derived from the square of the energy density function. The equally probable peaks in the magnetization likelihood are seen as small bright regions along the equator in Fig. **SN2H**. We assume that these localized regions are sufficiently narrow to approximate six discrete magnetization states.

We assume that transitions between these discrete states can be modeled as a continuous-time Markov Chain (CTMC) whose transition rates are related to the Néel relaxation time (*7*). The Néel relaxation time is the expected length of time between transitions between two magnetization states, usually defined in terms of uniaxial anisotropy as the time between magnetization reversals. In the one-dimensional case, given two magnetization states A and C with local minimum energies $U_A$ and $U_C$ separated by a maximum energy $U_B$, we can estimate the Néel relaxation time from state A to state C from the energy barrier between them using

$$\tau_{N,A\to C} = \frac{1}{v_0}\exp\left(-\frac{U_B - U_A}{k_B T}\right) \qquad \text{eq.14}$$

$$\tau_{N,C\to A} = \frac{1}{v_0}\exp\left(-\frac{U_B - U_C}{k_B T}\right) \qquad \text{eq.15}$$

where the attempt rate $v_0 \approx 10^9$ Hz is estimated to an order of magnitude (*7*). If there are multiple paths between states A and C (as is the case for our MNDs, where $\boldsymbol{w}$ can transition through a continuum of paths across the unit sphere), the exponential term in Eqs. (14) and (15) increases the likelihood of the minimum energy path. We can therefore reduce the continuous 2D state space (unit sphere) shown in **Fig SN2** to a discrete graph of states connected by relative transition probabilities. That is, the evolution of the magnetization state over a single time step of our simulation can be represented as a continuous time Markov chain (*8*).

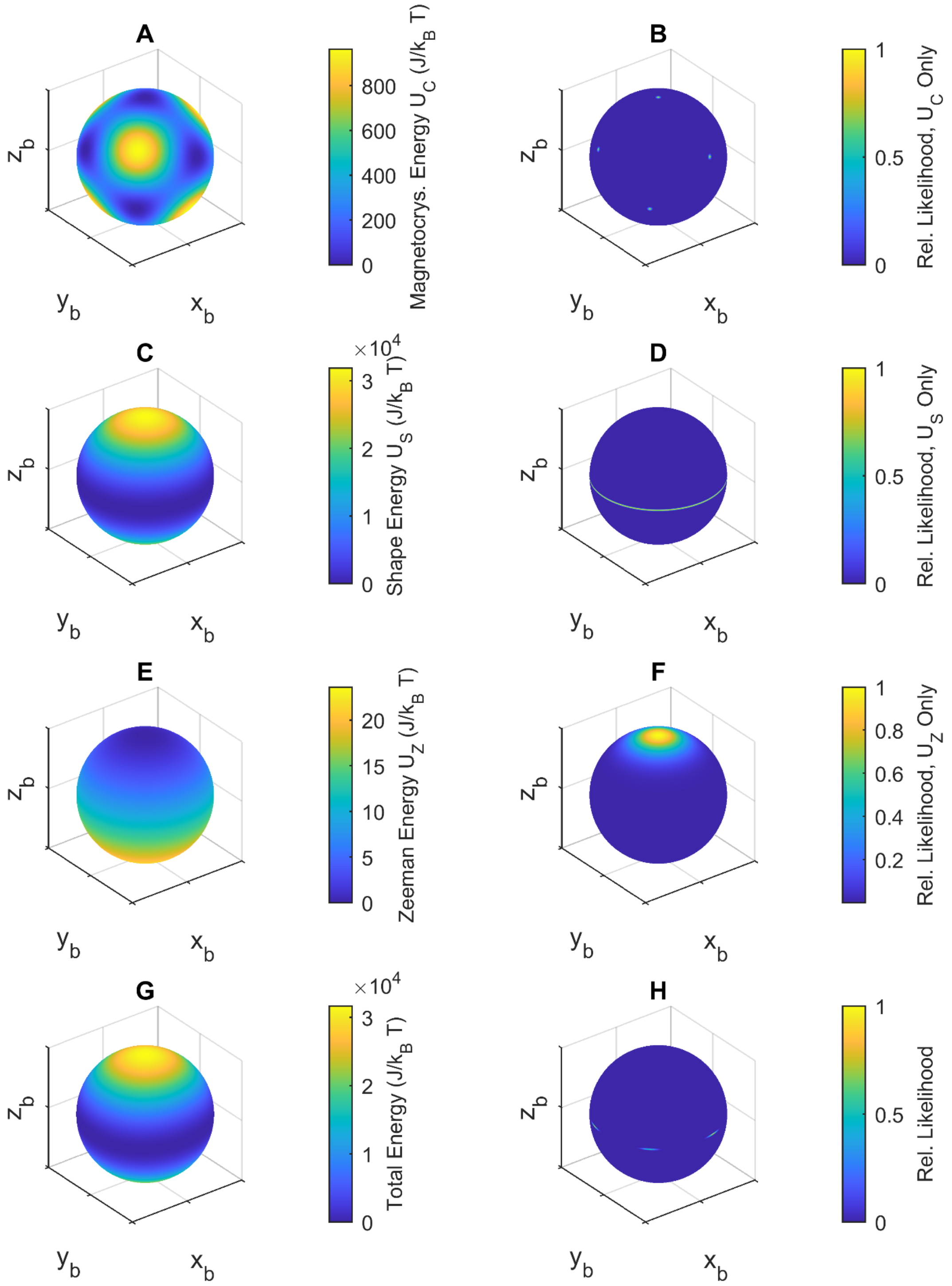


**Fig. SN2** Magnetic energy and relative likelihood plotted on the unit sphere for a simulated disc-shaped particle with 80 mT applied field in the positive Z-direction (the short axis of the disc). The vertical axis represents the short axis of the hexagonal disc (230 nm diameter and 30 nm thickness). **A,B** Magnetocrystalline anisotropy energy, **C,D** Shape anisotropy energy, **E,F** Zeeman energy, and **G,H** Total energy (A,C,E,G correspond to magnetic energy and B,D,F,H show probability function).

The generator matrix of the continuous time Markov chain for $n$ discrete states is:

$$G = \begin{bmatrix} \sum_{j\neq 1} g_{1j} & \cdots & g_{1n} \\ \vdots & \ddots & \vdots \\ g_{n1} & \cdots & \sum_{j\neq 1} g_{nj} \end{bmatrix} \text{a} \qquad \text{eq.16}$$

where $g_{ij}$ are the transition rates from state i to state j. In our simulation, the transition rates are the reciprocal of the Néel relaxation times between each state. Given a known probability density $\boldsymbol{P}(t)$ at some time $t$, the probability density at a later time $\boldsymbol{P}(t+\tau)$ is

$$\boldsymbol{P}(t+\tau) = \exp(\tau G)\,\boldsymbol{P}(t) \qquad \text{eq.17}$$

The minimum energy path and energy barriers between the discrete states shown in Fig. S2 H can be found using the nudged elastic band (NEB) method (*9*). The results of the NEB method for the energy landscape of a disc-shaped nanoparticle with an 80 mT applied field in the positive Z direction (short axis of the disc) are shown in **Fig. SN2H**. Here, we obtained the energy minima (states) and energy maxima (energy barriers).

We applied the magnetic model described in this section to develop an algorithm for calculating dynamic changes in the magnetization direction of a MND due to an applied field. Given an initial distribution $\boldsymbol{P}(t)$ (an impulse distribution centered on the previous state), an applied field $\boldsymbol{B}_a$ and the magnetic properties of MNDs, the algorithm uses the NEB method to find the local energy minima and the energy barriers between them. Then it calculates the generator matrix and applies Eq. (17) to calculate the final distribution $\boldsymbol{P}(t+\tau)$. Finally, it randomly chooses the final state with the likelihood of each state being chosen proportional to $\boldsymbol{P}(t+\tau)$.

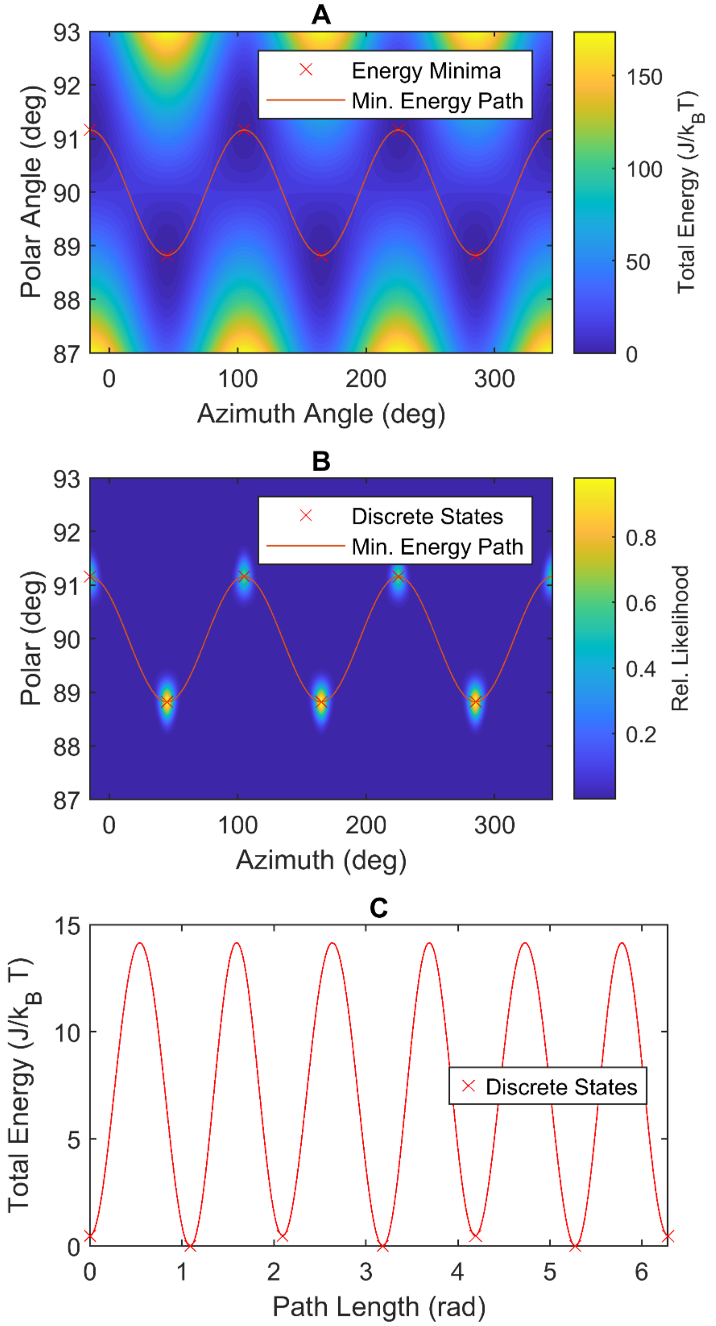


**Fig. SN3** **A** Energy landscape of the magnetization state parameterized by the azimuth angle (angle from the x-axis to the projection of $\boldsymbol{w}$) and the polar angle (angle from the $z$-axis to $\boldsymbol{w}$) for a simulated disc-shaped particle with 80 mT applied field at a polar angle of zero degrees (the short axis of the disc). The local energy minima are indicated by red x marks, and the minimum energy path connecting them is a red line. **B** Relative likelihood of the magnetization state parameterized by the azimuth and polar angles. The discrete states corresponding to each local energy minimum are indicated by red x marks. **C** The energy plotted as a function of path length (in radians) along the minimum energy path between the energy minima (red x marks). The energy barriers between states are the energy maxima of this function.

346

#### 1.3 Fluidic Resistance Model

The hydrodynamic resistance force $\boldsymbol{F}$ and torque $\boldsymbol{T}_f$ (about any point $O$ rigidly attached to the particle) for a translating and rotating particle in a fluid medium with Reynolds number $Re \ll 1$ and with the fluid at rest at infinity is given by:

$$\begin{pmatrix} \boldsymbol{F} \\ \boldsymbol{T}_f \end{pmatrix} = \eta \begin{pmatrix} K & C_O^T \\ C_O & \Omega_O \end{pmatrix} \begin{pmatrix} \boldsymbol{U}_O \\ \boldsymbol{\omega} \end{pmatrix} \qquad \text{eq.18}$$

where $\eta$ is the dynamic viscosity of the fluid, $K$ is the hydrodynamic translation tensor, $C_O$ is the hydrodynamic coupling tensor, $\Omega_O$ is the hydrodynamic rotation tensor, $\boldsymbol{U}_O$ is the velocity of the point on the particle, and $\boldsymbol{\omega}$ is the angular velocity of the particle. The expressions for these tensors are described by Gappel and Brenner (1983) (*10*).

All particles have a unique point $R$ where the coupling tensor $C_R$ is symmetric. For any body with three mutually perpendicular planes of symmetry, such as a hexagonal MND, $R$ lies at the intersection of the planes of symmetry, $C_R = (0)$, and $K$ and $\Omega_R$ are diagonal if their components are defined along the lines of intersection of the three planes of symmetry (the body frame). In this case, equation (18) becomes:

$$\begin{pmatrix} \boldsymbol{F} \\ \boldsymbol{T}_f \end{pmatrix} = \eta \begin{pmatrix} K & -(\boldsymbol{r}_{RO} \times K)^T \\ -\boldsymbol{r}_{RO} \times K & \Omega_R - \boldsymbol{r}_{RO} \times K \times \boldsymbol{r}_{RO} \end{pmatrix} \begin{pmatrix} \boldsymbol{U}_O \\ \boldsymbol{\omega} \end{pmatrix} \qquad \text{eq.19}$$

where $\boldsymbol{r}_{RO}$ is the vector from the point $O$ to the point $R$. Then

$$\Omega_O = \Omega_R - \boldsymbol{r}_{RO} \times K \times \boldsymbol{r}_{RO}\boldsymbol{\omega} \qquad \text{eq.20}$$

This equation for $\Omega_O$ is used in Eq. (3) to find the rate of MND rotation.

There are no general analytical expressions for the resistance tensors $\Omega_R$ and $K$ of a hexagonal disc. However, the dynamic force and torque on a particle at low Reynolds numbers are bounded from above and below by the force and torque on a circumscribing body. We therefore take the minimum spheroid of the same aspect ratio that circumscribes the hexagonal MND to calculate the hydrodynamic resistance to its motion. We used the expressions for $\Omega_R$ and $K$ given by Happel and Brenner (1983) (*4*) for a spheroidal particle.

### 1.4 Combined Dynamic Model

We implemented the dynamic model described in **Section 1.1** in MATLAB Simulink R2024b. The dynamic model generates an applied MF on the MND. This applied MF is used to calculate the magnetization of the particle using the magnetization model in **Section 1.2**. The magnetization is then used to calculate the magnetic torque on the MND. A white noise term is added to account for the Brownian motion of the nanoparticle. These torques are then used in Eq. (3) to find the angular velocity of the MND, which is used to update the MND orientation for the next time step. The MATLAB code is available within the data repository.

### Supplementary Note 2

**Theoretical maximum torque per particle at 80 mT MF**

In a uniform magnetic field, the torque on a magnetic dipole is calculated as:

$$\boldsymbol{T}_m = \boldsymbol{m} \times \boldsymbol{B}_a$$

where $\boldsymbol{m}$ is the dipole moment and $\boldsymbol{B}_a$ is the applied MF (*11*).

When the magnetic moment and the MF are orthogonal, the maximum torque is equal to:

$$|\boldsymbol{T}_{\boldsymbol{m}}| = |\boldsymbol{m}||\boldsymbol{B}_a|$$

An MND uniformly magnetized under a magnetic field of 80 mT can generate a maximum theoretical torque:

$$|\boldsymbol{T}_{\boldsymbol{m}}| = 6.3 \times 10^{-16} Am^2 \times 0.080\, T = 5.04 \times 10^{-17} Nm$$

As the force on the edge of a disk generated by this magnetic torque is scales inversely with the disk radius, the mechanical force from 230 nm-diameter MND is

$$F = 5.04 \times 10^{-17} Nm \div (115 \times 10^{-9}\, m) = 438\, pN$$

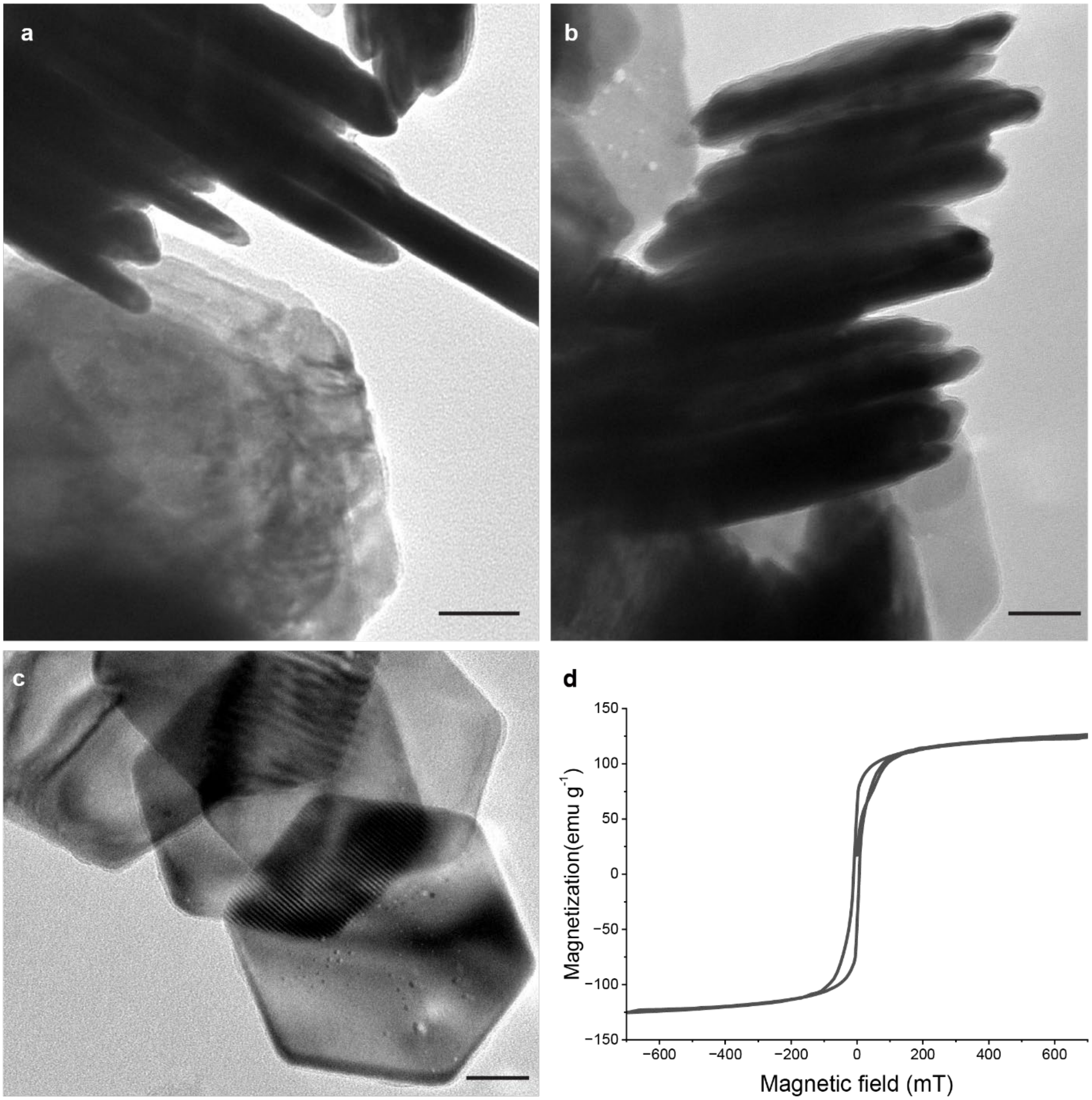


**Fig. S1 Synthesis and characterization of MNDs. a,** MNDs before coating with BG-PEG-PMAO-Cy7 polymer. **b,** Side view of MNDs after polymer coating, showing the surface morphology of the MNDs being more irregular than uncoated MNDs. **c,** Top view of MNDs after polymer coating, demonstrating a uniform thickness of BG-PEG-PMAO-Cy7 polymer coating (5 nm). Scale bars = 50 nm. **d,** Hysteresis loop measured from a vibrating-sample magnetometer on polymer-coated MNDs dispersed in 0.6% w/v agarose gel to mimic brain mechanical properties.

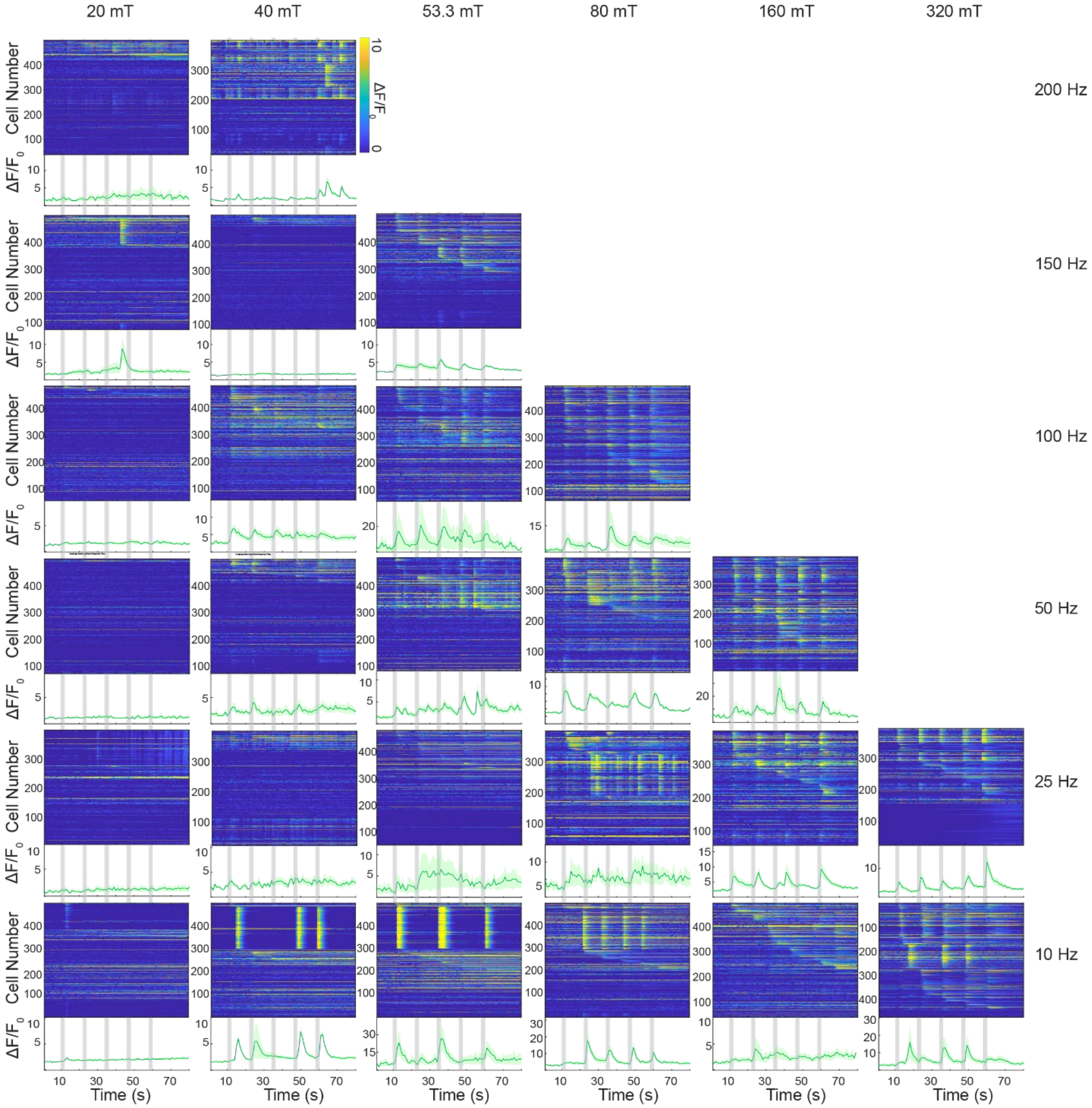


**Fig. S2 Response diagram for magnetomechanical neuromodulation conditions.** GCaMP6s relative fluorescence traces of individual cells (top panels) and their averages (bottom panels) in response to MF (epochs indicated with vertical grey bars) with different frequencies and amplitudes. In the bottom plots lines and shaded areas represent the mean and standard error of the mean (s.e.m.). Data was collected from three independent cultures.

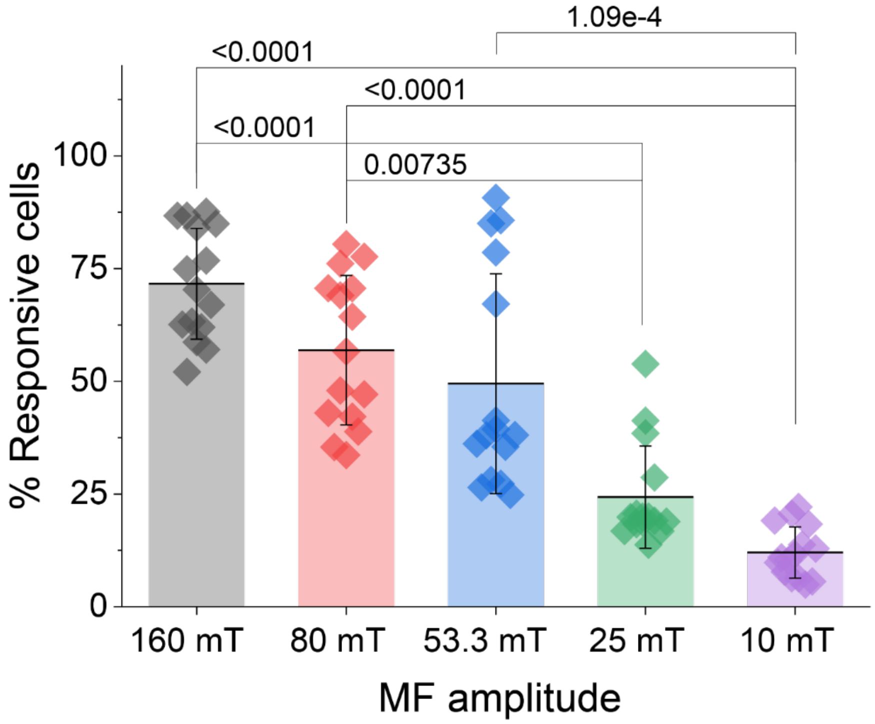


**Fig. S3 Statistical analysis on calcium imaging results from MND-decorated neurons exposed to MF with 50 Hz and different amplitudes.** As the data distribution does not fulfill the normality, statistical analysis was conducted using the Kruskall-Wallis ANOVA test followed by Dunn's post-hoc test (n=15 for each condition; each data point represents one MF application out of five applied to three different coverslips containing neurons); $\chi^2 = 53.23$, DF=4, p<0.0001. Bars and error bars represent mean ± standard deviation (s.d.).

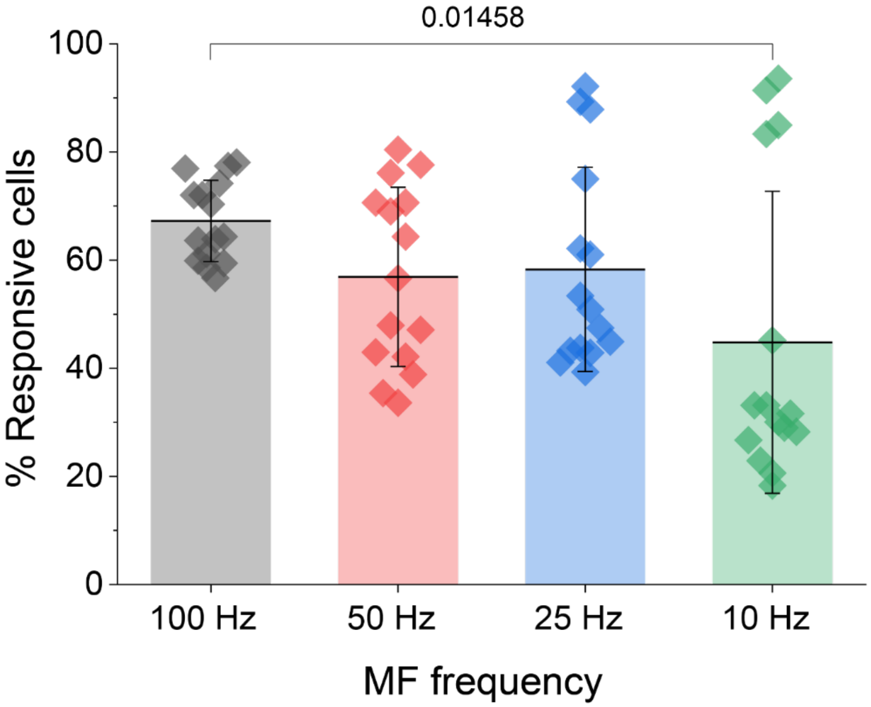


**Fig. S4 Statistical analysis on calcium imaging results from MND-decorated neurons exposed to MF of 80 mT and different frequencies.** As the data distribution does not fulfill the normality, statistical analysis was conducted using the Kruskall-Wallis ANOVA test followed by Dunn's post-hoc test (n=15 for each condition; each data point represents one MF application out of five applied to three different coverslips containing neurons); $\chi^2 =$ 9.26, DF=3, p=0.026. Bars and error bars represent mean $\pm$ s.d.

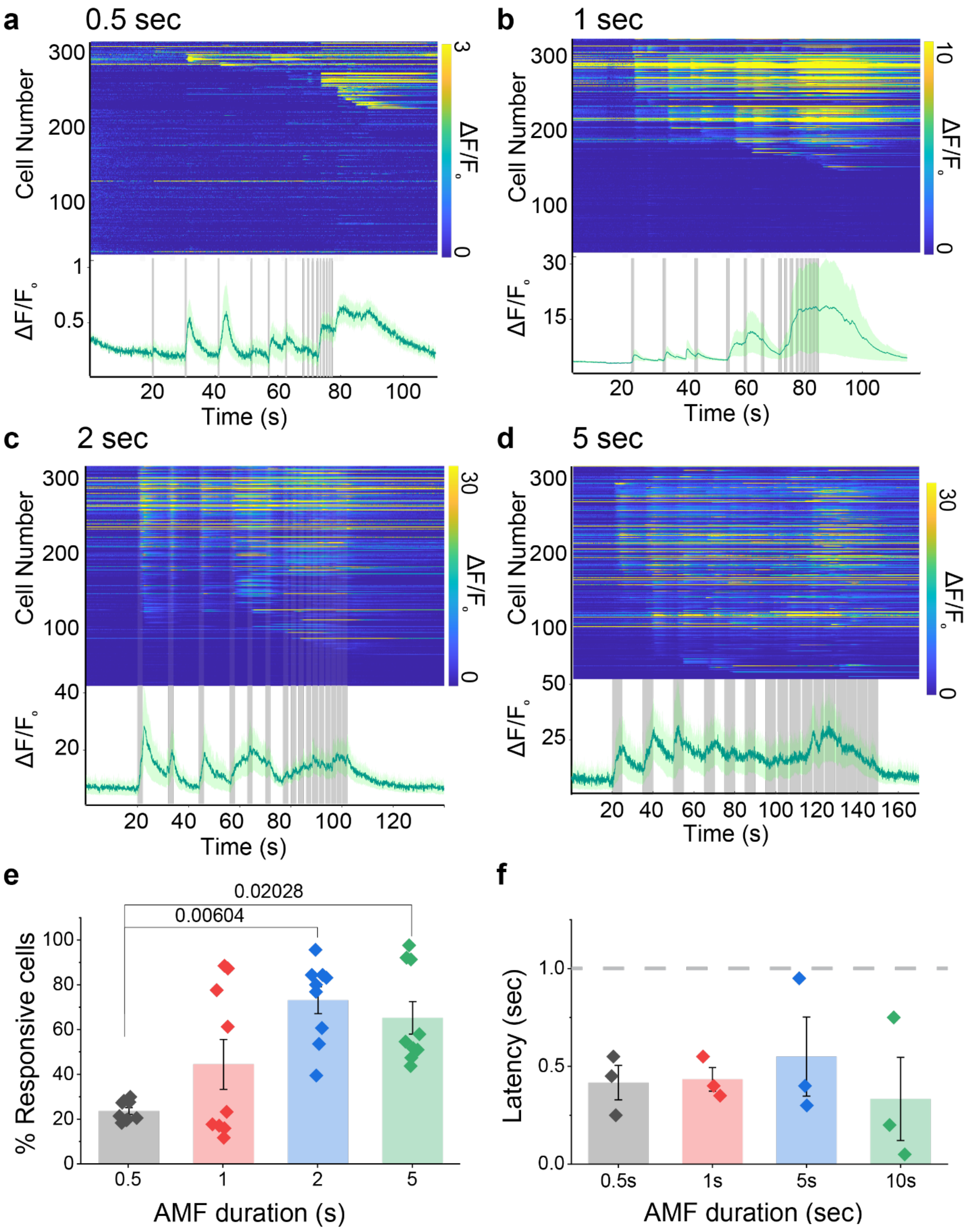


**Fig. S5 MF duration and period variation and its quantitative comparison regarding responsivity and latency.** GCaMP6s fluorescence traces of individual cells and their average under MF (80 mT 50 Hz) with various intervals between MF epochs having durations **a,** 0.5s, **b,** 1s, **c,** 2s, and **d,** 5s. **e,** Quantitative comparison of responding cell percentage when different duration of MF was applied. As the data distribution does not fulfill the normality, statistical analysis was conducted using the Kruskal-Wallis ANOVA test followed by Dunn's post-hoc test (n=9 for each condition); $\chi^2$ = 14.52, DF=3, p=0.0027. (A-D, bottom panels) Lines and shaded areas represent the mean $\pm$ s.e.m. **f,** Comparison of the latency of the neurons when different duration of MF was applied. 1 sec is indicated with grey dotted line. As the data distribution fulfill the normality, statistical analysis was conducted using the one-way ANOVA test followed by Tukey's post-hoc test (n=3 for 0.5s, n=9 for other condition); DF=3, p=0.109. Latency has been analyzed for the first three stimulations with 10-second intervals, as shorter intervals disrupt neuronal response synchronization to MF applications, making it difficult to determine the latency and response of neurons per stimulation. Each data point in the responsive cell percentage plot represents the percentage of cells, per MF epoch, that exhibit a relative fluorescence change greater than three times their own baseline (20s before the initial MF onset) standard deviation. Each data point in the latency plot indicates the location of the peak in the average relative fluorescence per coverslip per MF epoch. For 0.5s, six out of nine trials did not produce a peak detected by the peakfinder function in MATLAB. Bars and error bars represent the mean $\pm$ s.e.m.

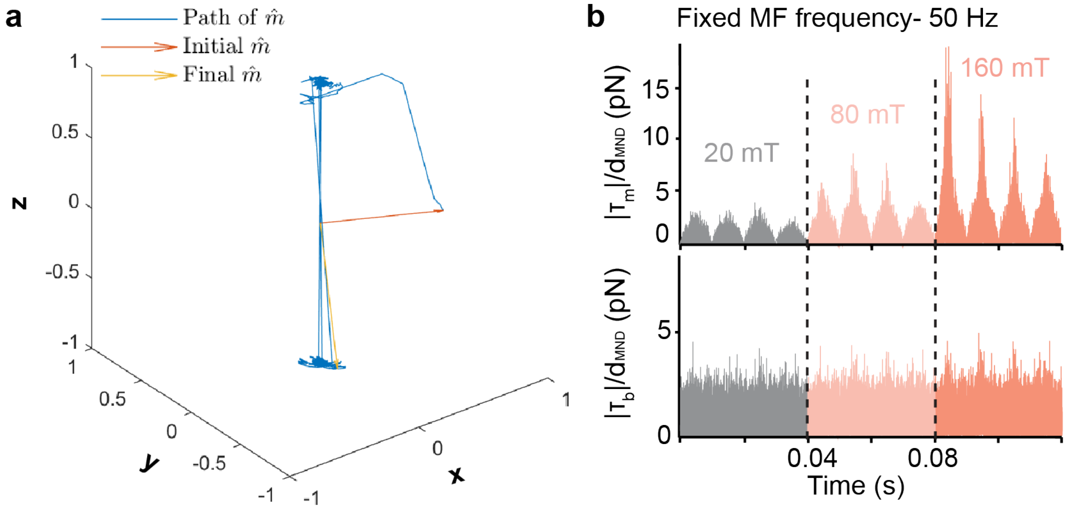


**Fig. S6 Computational simulation for mechanical force generated by MNDs. a,** Simulated traces of MNDs' magnetic moment. **b,** Simulated $\mathbf{T_m}$ and $\mathbf{T_b}$ at varying MF amplitudes with fixed frequency (50 Hz), showing frequency- and amplitude-dependent mechanical output.

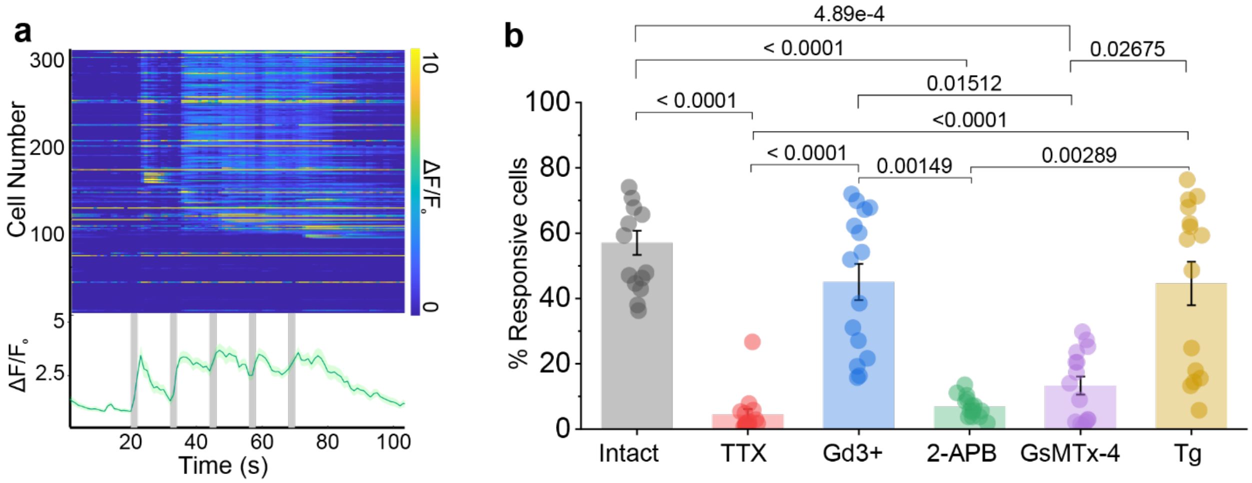


**Fig. S7 $Ca^{2+}$ imaging with pharmacological treatment. a,** GCaMP6s imaging performed with Tg treatment. Lines and shaded areas represemt the mean ± s.e.m. **b,** Quantitative comparison of the responsivity of cultured hippocampal neurons treated with different drugs, compared to intact neurons that were not exposed to any drug. As the data distribution does not fulfill the normality, statistical analysis was conducted using the Kruskall-Wallis ANOVA test followed by Dunn's post-hoc test (n=15 in each condition; each data point represents one MF application out of five applied to three different coverslips containing neurons); $\chi^2 =$ 60.23, DF=5, p<0.0001. Bars and error bars represent the mean ± s.e.m.

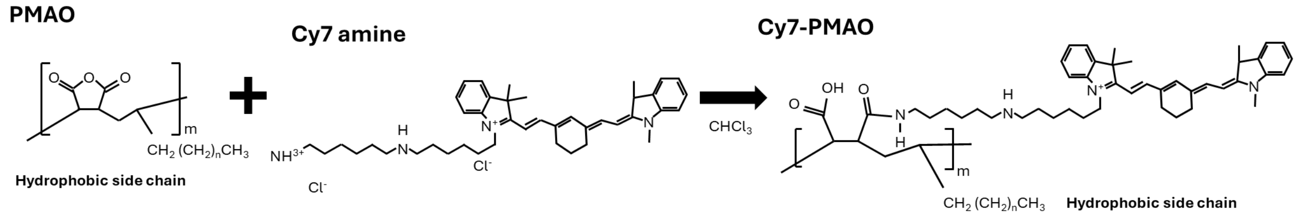


**Fig. S8 Structure of fluorescently-labeled PMA.** Schematic of a reaction of PMAO (poly(maleic anhydride-alt-1-octadecene)) with Cy7 dye.

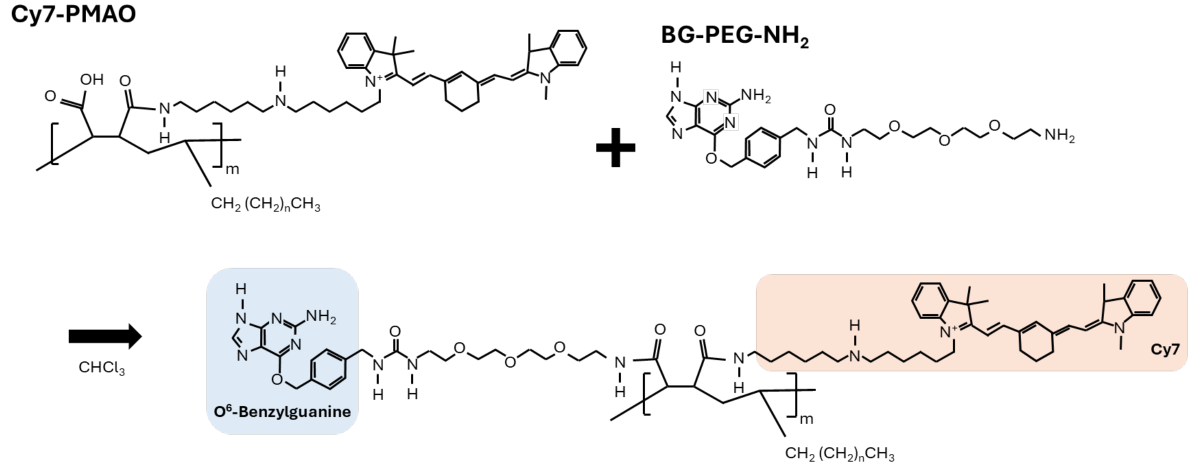


**Fig. S9 Structure of Cy7-PMAO-PEG-BG.** Reaction between Cy7-dye labeled PMAO and amine-terminated poly(ethylene glycol) (PEG) functionalized with benzyl guanine (BG).

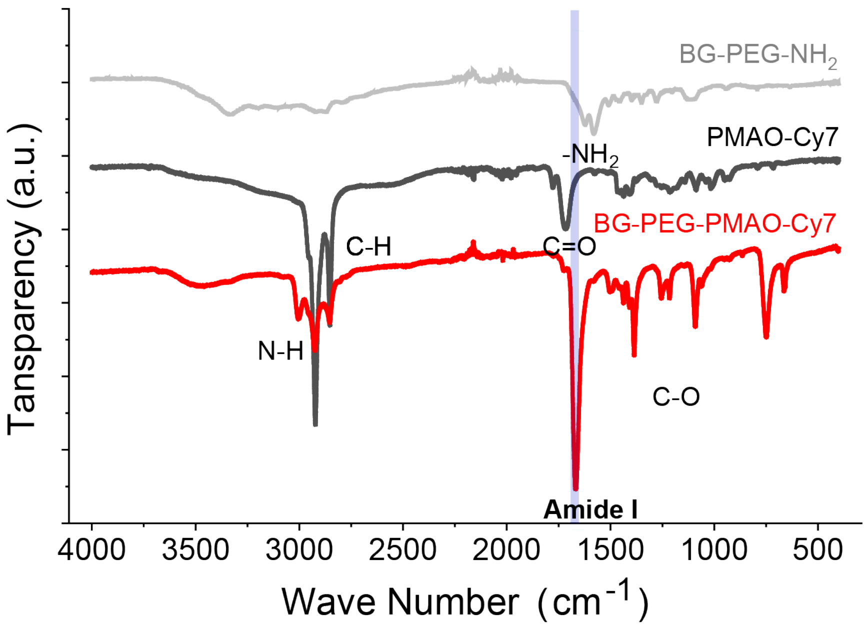


**Fig. S10 BG-PEG-PMAO-Cy7 detected in Fourier Transform Infrared (FTIR) spectroscopy.** FTIR spectra for MND ligands (i) benzylguanine (BG)-PEG-NH2, (ii) poly(maleic anhydride) 1-octadecene (PMAO)-Cy7, and (iii) BG-PEG-PMAO-Cy7. Peaks are indicated for the pendant amine in BG-PEG-NH2 (1620 $cm^{-1}$), for the anhydride carbonyl peaks in PMAO (1750 and 1820 $cm^{-1}$, with the 1750 $cm^{-1}$ peak being larger for a cyclic anhydride), and for the amide I peak arising from the fusion of PEG and PMAO in BG-PEG-PMAO-Cy7 (1650-1690 $cm^{-1}$, denoted in highlighted region). C-H and N-H stretches from the allylic carbons are also denoted in the region of 3000 $cm^{-1}$. The appearance of the amide I peak was taken as a positive indicator of the fusion of PMAO and BG-PEG.

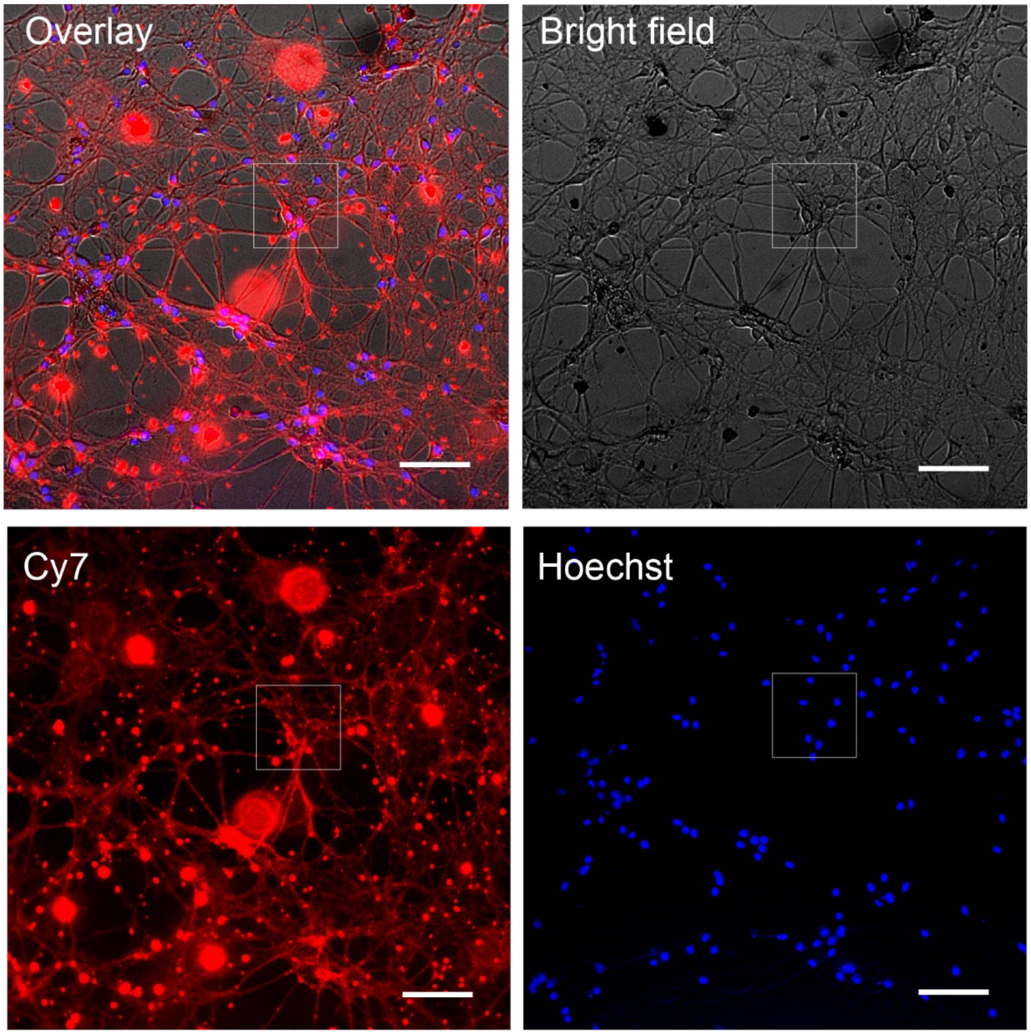


**Fig. S11 MNDs on SNAP tag-expressing hippocampal neurons.** After 1h of incubation of the cultured hippocampal neurons transduced with AAV9−Ef1α::SNAP-tag−PDGFR in (4.0 $\mu$g cm$^{-2}$) MND-dispersed Tyrode's solution. Left top: overlay, right top: bright field, left bottom: MNDs coated by BG-PEG-PMAO-Cy7. Bottom right: Hoechst. Scale bars = 100 μm.

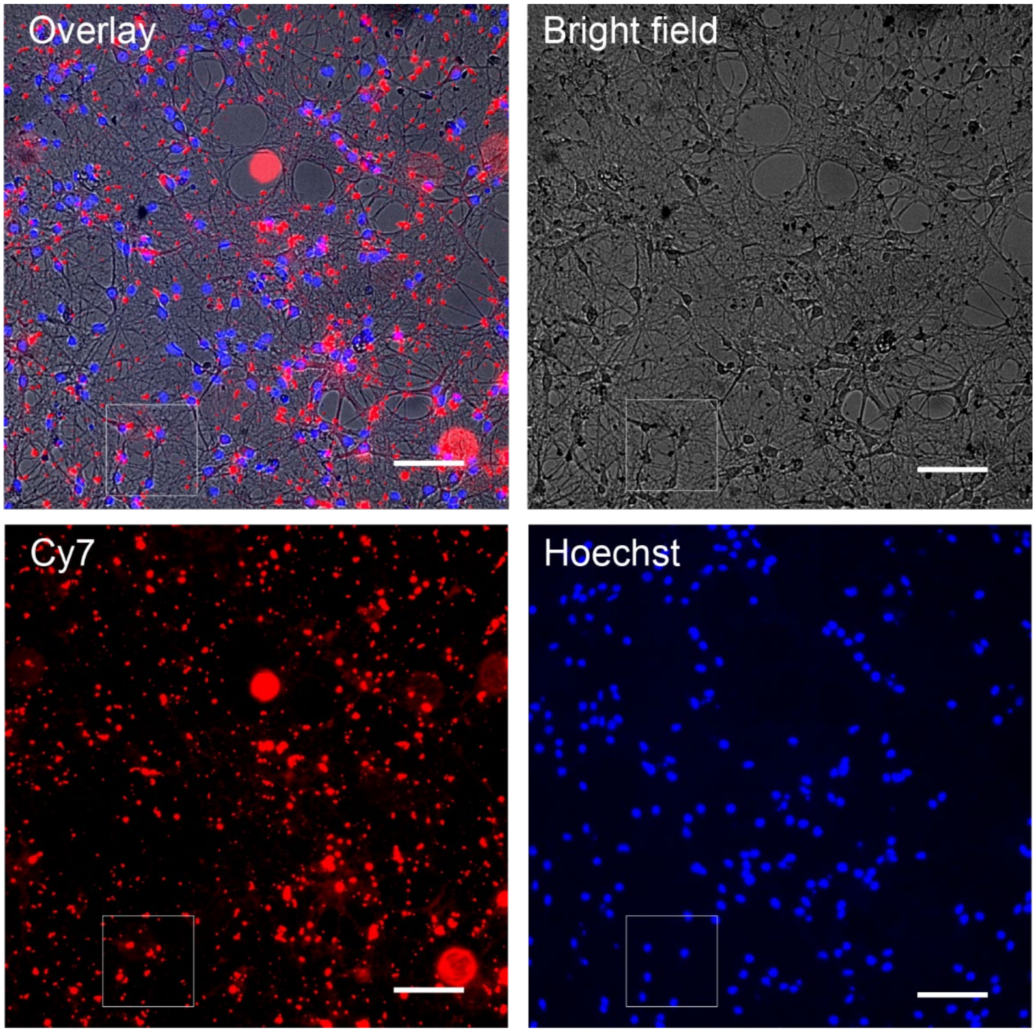


**Fig. S12 MNDs on genetically intact cultured hippocampal neurons** After 1h of incubation of the cultured hippocampal neurons in (4.0 $\mu$g cm$^{-2}$) MND-dispersed Tyrode's solution. Left top: overlay, right top: bright field, left bottom: MNDs coated by BG-PEG-PMAO-Cy7. Bottom right: Hoechst. Scale bar is 100 μm.

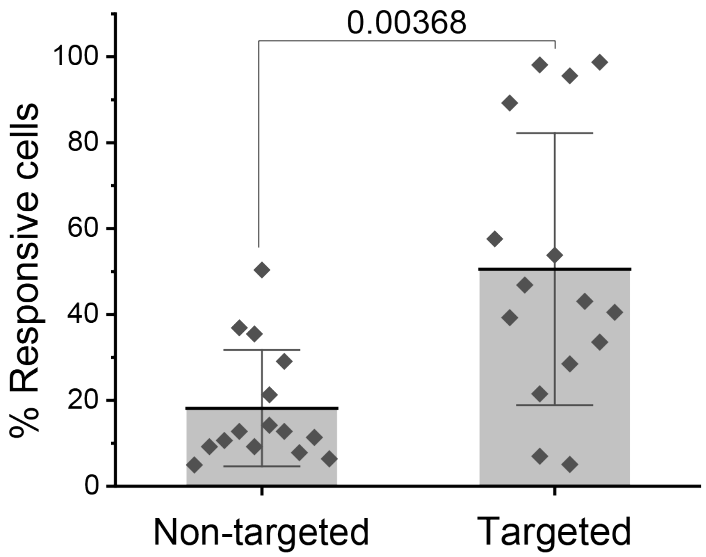


**Fig. S13 Quantified comparison of neuronal responsivity.** As the data distribution does not fulfill normality, statistical analysis was conducted using the Mann-Whitney U test to compare the two independent groups (non-target n=15, target n=15). Each data point represents one MF application out of five applied to three different coverslips containing neurons. U = 183, Z = 2.90. Bars and error bars represent the mean ± s.d.

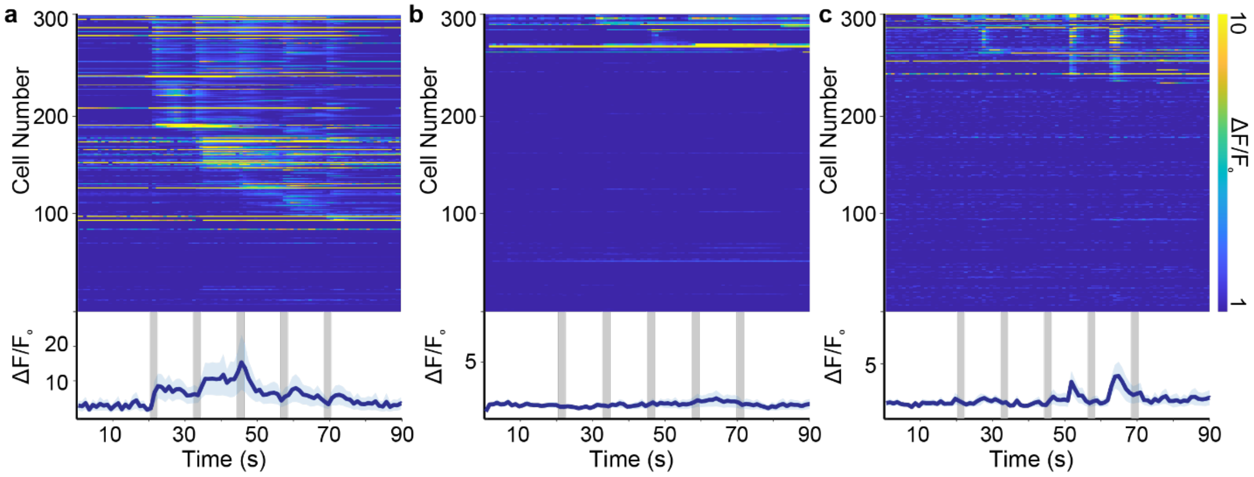


**Fig. S14 Variation in MNDs density and its effect on neuronal responses in calcium imaging.** **a,** Calcium imaging on the cultured hippocampal neurons expressing SNAP-tag with MNDs density of 7.9 $\mu$g cm$^{-2}$. **b,** Calcium imaging on the cultured hippocampal neurons not expressing SNAP-tag with MNDs density of 7.9 $\mu$g cm$^{-2}$. **c,** Calcium imaging on the cultured hippocampal neurons expressing SNAP-tag with MNDs density of 4.0 $\mu$g cm$^{-2}$. MF (80 mT, 50 Hz) application is indicated by grey bars. The data is shown with the average ± s.e.m.

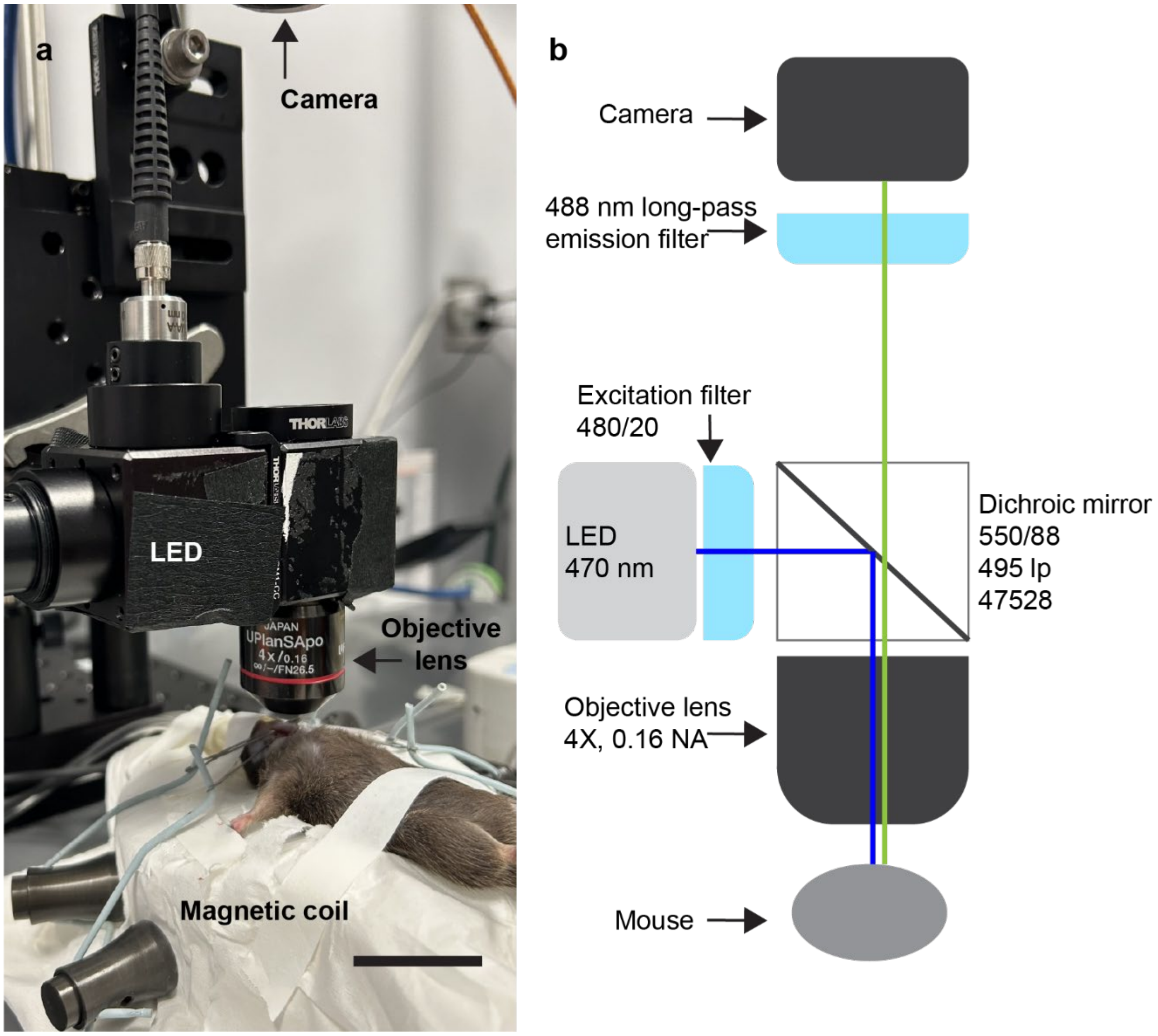


**Fig. S15 Setup of in-vivo $Ca^{2+}$ imaging combined with magnetic stimulation. a,** Photograph of the setup. Scale bar, 2 cm. **b,** A diagram of the optical setup.

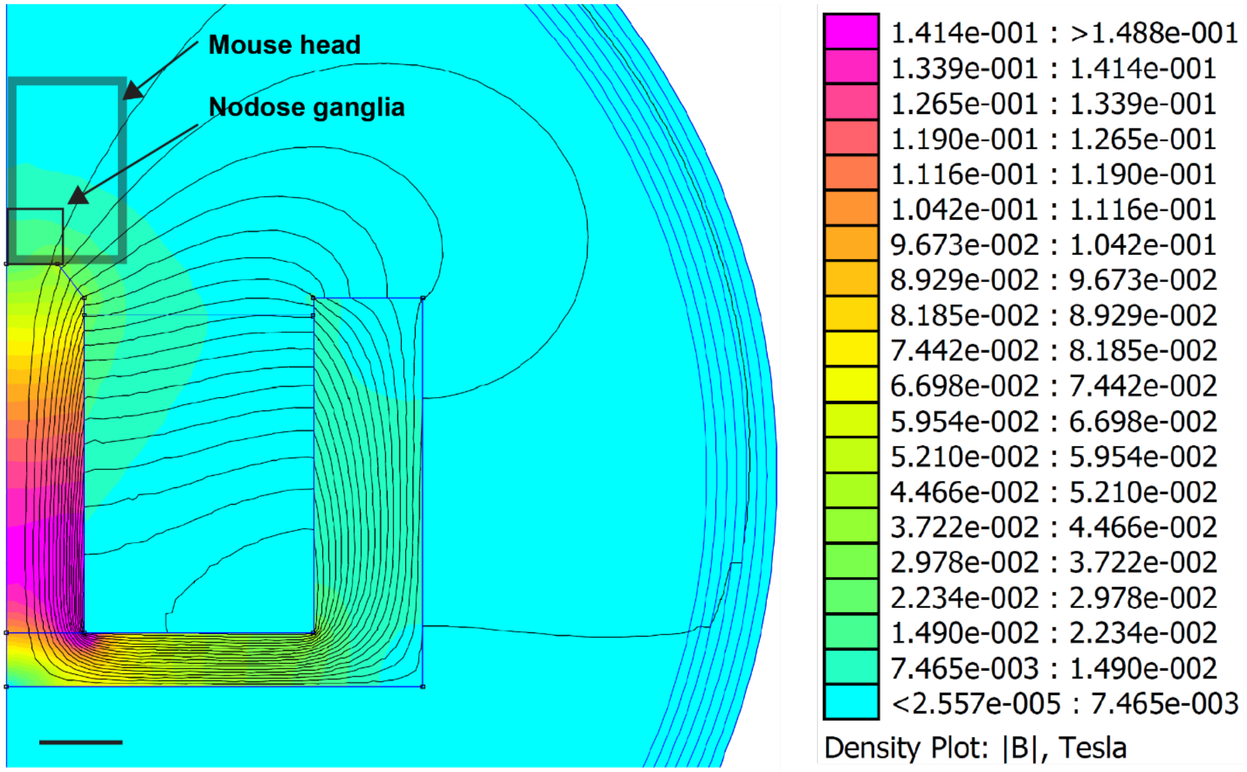


**Fig. S16 Simulation of a magnetic field profile generated by the electromagnet.** Scale bar, 1 cm.

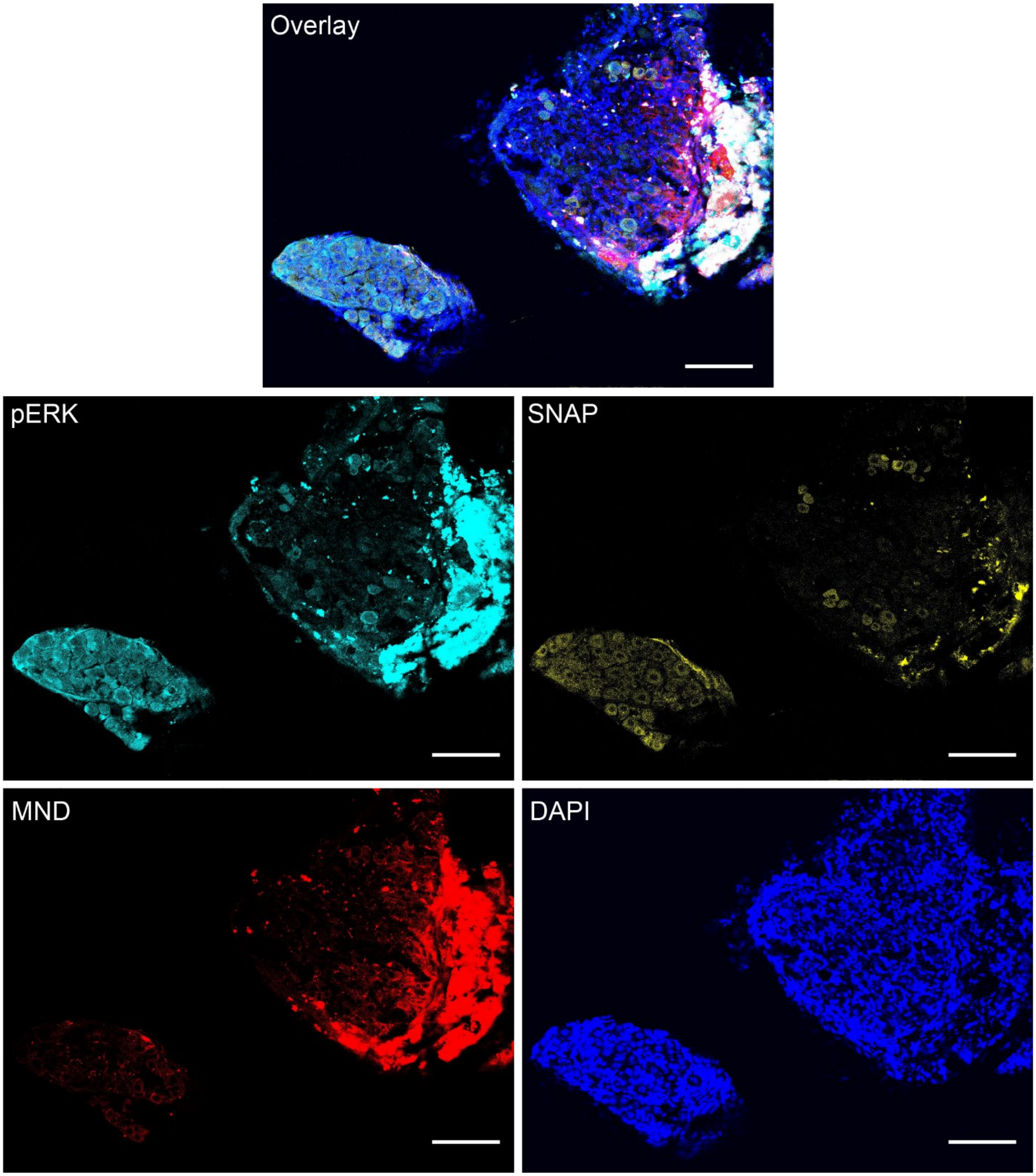


**Fig. S17 Oxtr-Cre$^{+/-}$ left NG.** Immunofluorescence image of the *left* NG in a Oxtr-Cre$^{+/-}$ mouse (control side without MND injection); **yellow indicates** SNAP-tag, cyan indicates pERK, **blue** indicates DAPI-labeled nuclei, and **red** shows Cy7 fluorescence marking MND localization. Scale bars = 100 μm.

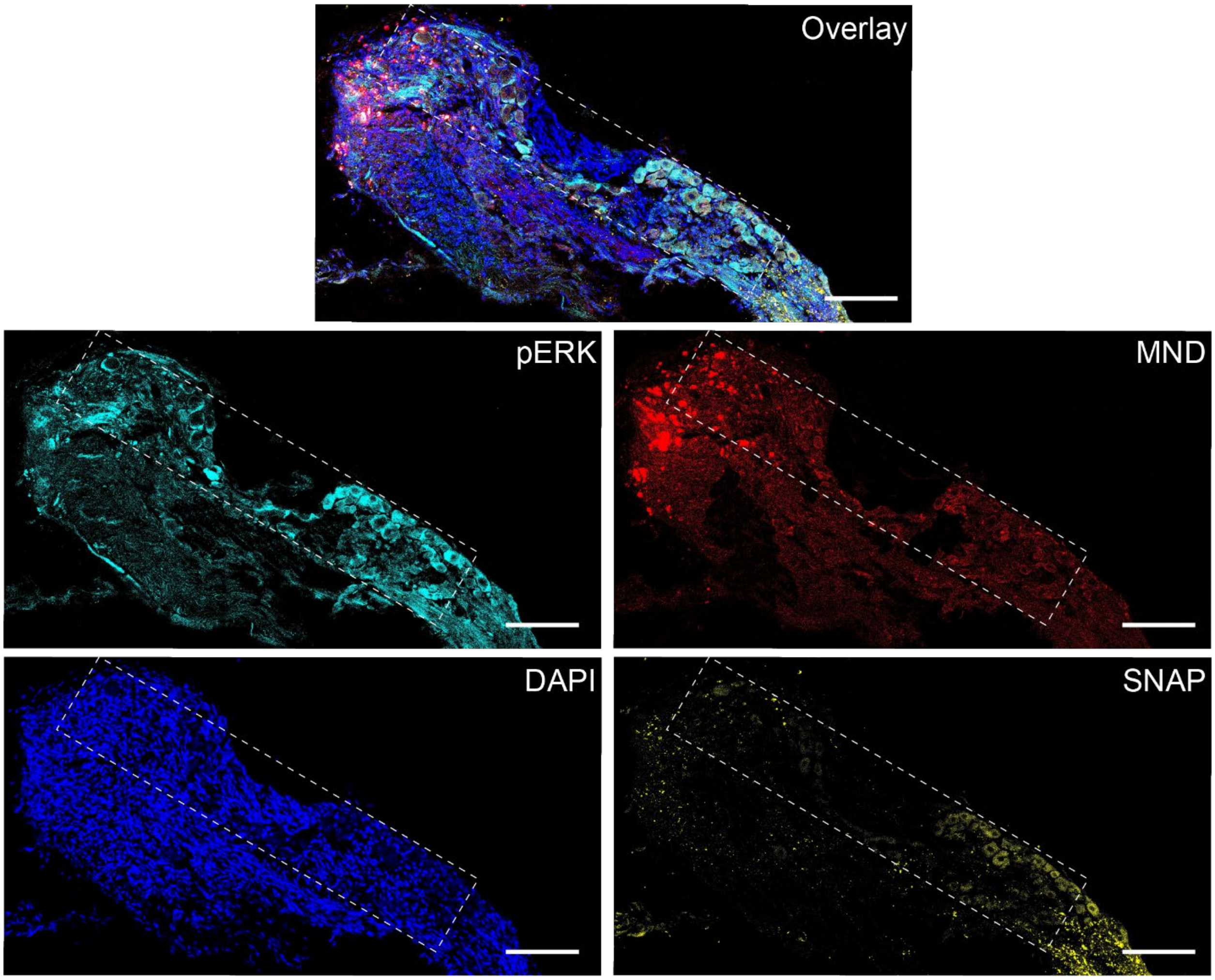


**Fig. S18 Glp1r-Cre$^{+/-}$ left NG.** Immunofluorescence image of the *left* NG in a Glp1r-Cre$^{+}$/$^{-}$ mouse (control side without MND injection); **yellow indicates** SNAP-tag, cyan indicates pERK, **blue** indicates DAPI-labeled nuclei, and **red** shows Cy7 fluorescence marking MND localization. Scale bars, 100 μm.

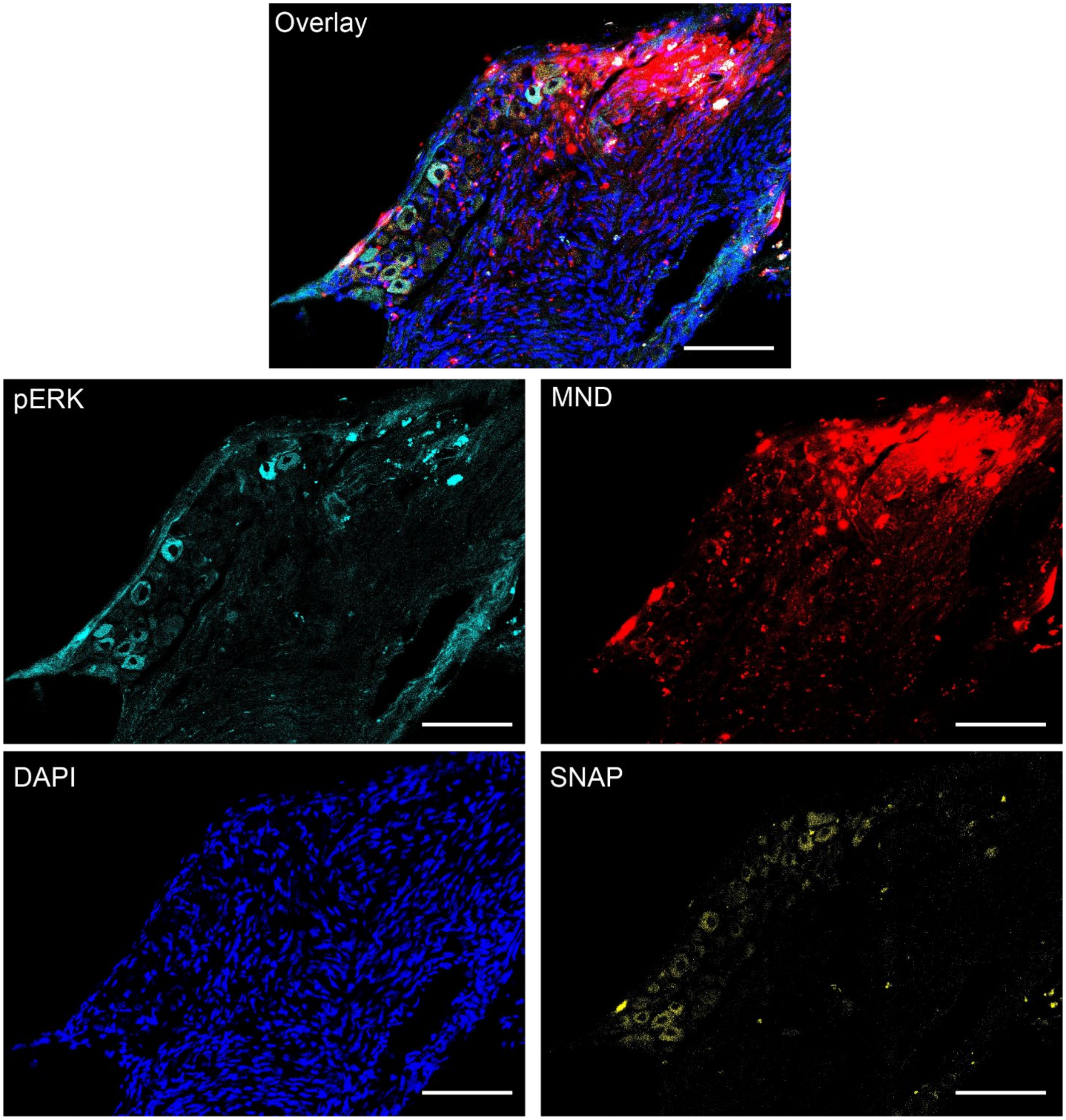


**Fig. S19 Gpr65-Cre$^{+/-}$ left NG.** Immunofluorescence image of the *left* NG in a Gpr65-Cre$^{+/-}$ mouse (control side without MND injection); **yellow indicates** SNAP-tag, cyan indicates pERK, **blue** indicates DAPI-labeled nuclei, and **red** shows Cy7 fluorescence marking MND localization. Scale bars, 100 µm.

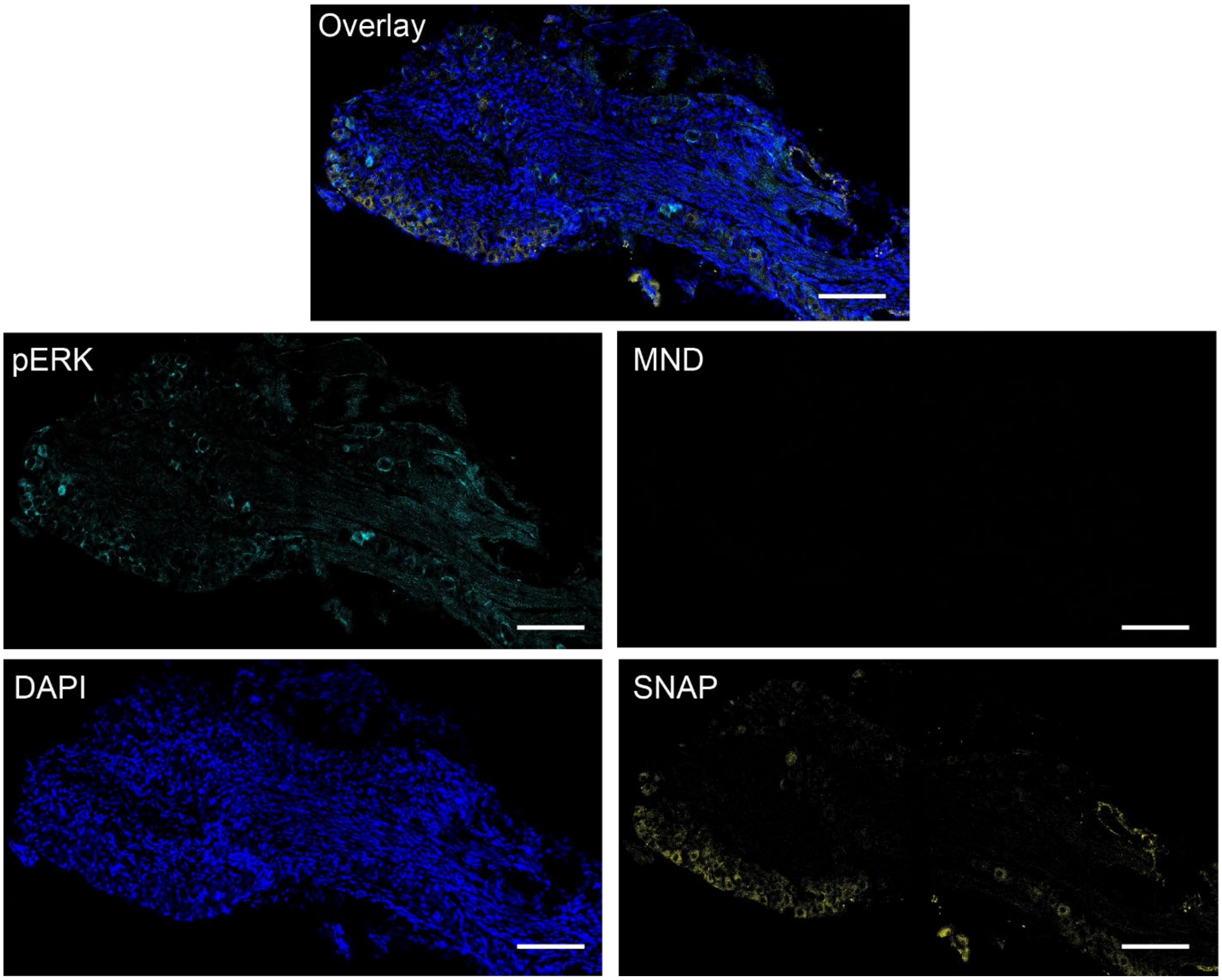


**Fig. S20 Oxtr-Cre$^{+/-}$ right NG.** Immunofluorescence image of the *right* NG in a Oxtr-Cre$^{+/-}$ mouse (control side without MND injection); **yellow indicates** SNAP-tag, cyan indicates pERK, **blue** indicates DAPI-labeled nuclei, and **red** shows Cy7 fluorescence marking MND localization. Scale bars, 100 µm.

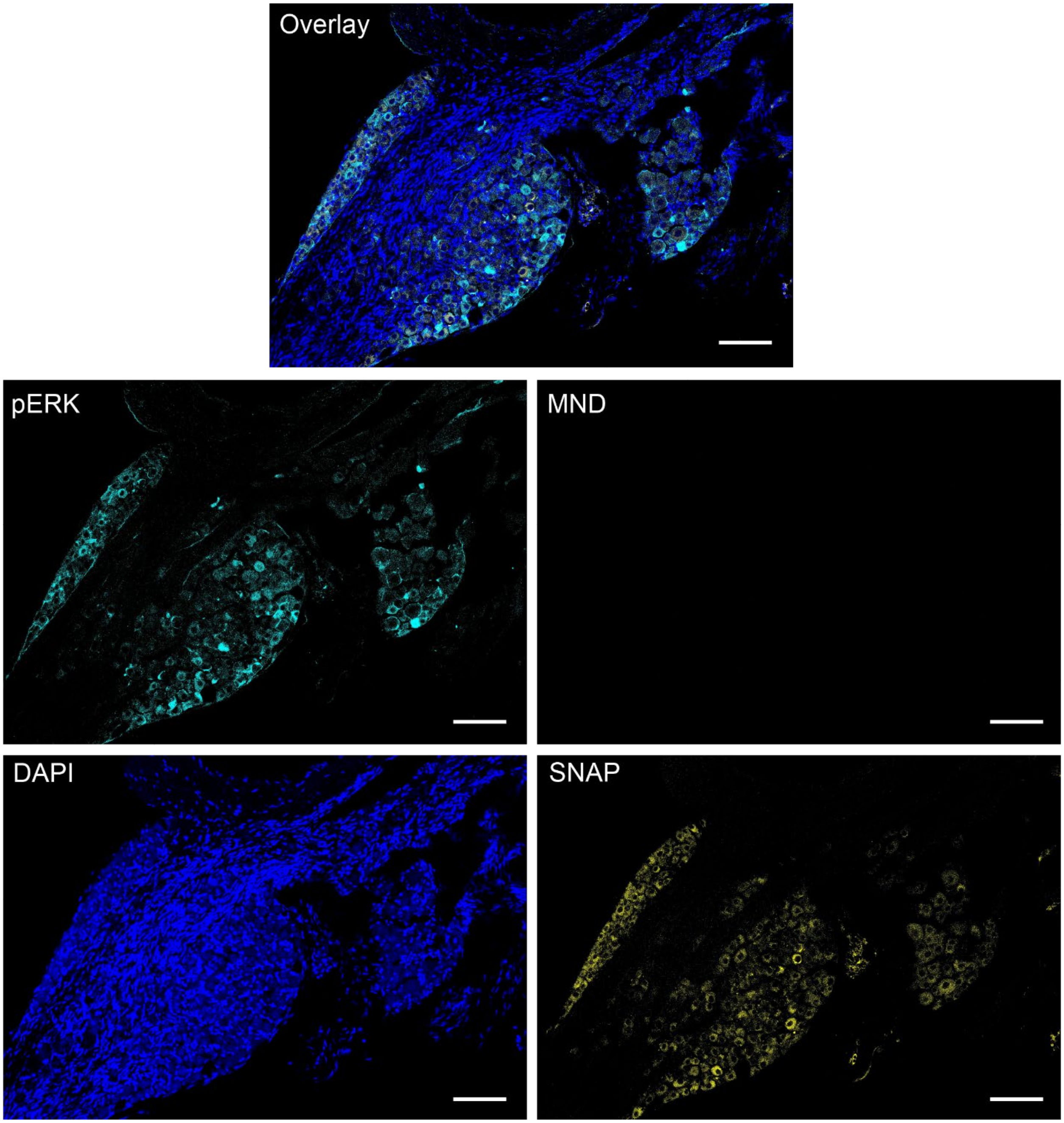


**Fig. S21 Glp1r-Cre$^{+/-}$ right NG** Immunofluorescence image of the *right* NG in a Glp1r-Cre$^{+/-}$ mouse (control side without MND injection); **yellow indicates** SNAP-tag, cyan indicates pERK, **blue** indicates DAPI-labeled nuclei, and **red** shows Cy7 fluorescence marking MND localization. Scale bars, 100 µm.

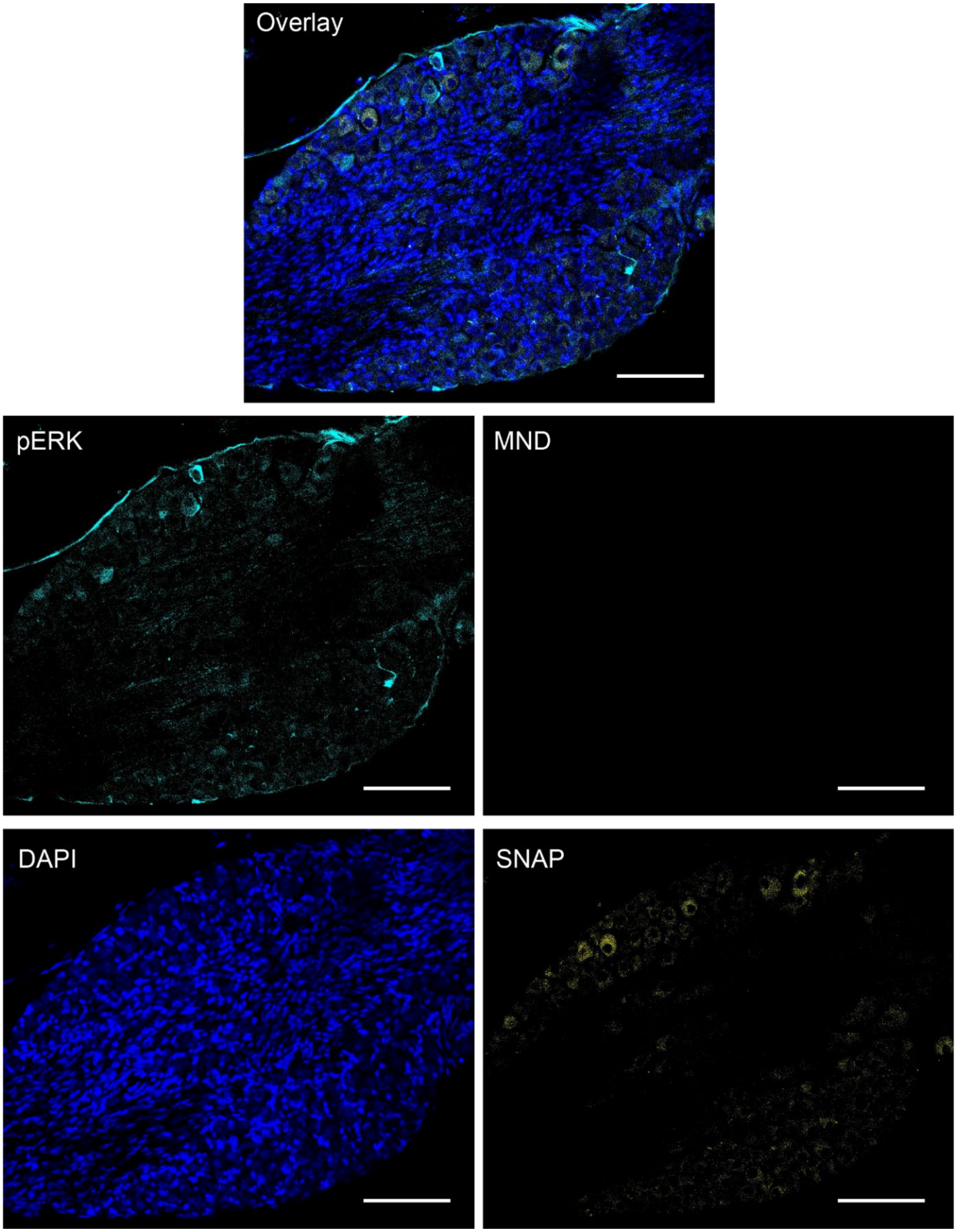


**Fig. S22 Gpr65-Cre$^{+/-}$ right NG.** Immunofluorescence image of the *right* NG in a Gpr65-Cre$^{+/-}$ mouse (control side without MND injection); **yellow indicates** SNAP-tag, cyan indicates pERK, **blue** indicates DAPI-labeled nuclei, and **red** shows Cy7 fluorescence marking MND localization. Scale bars, 100 μm.

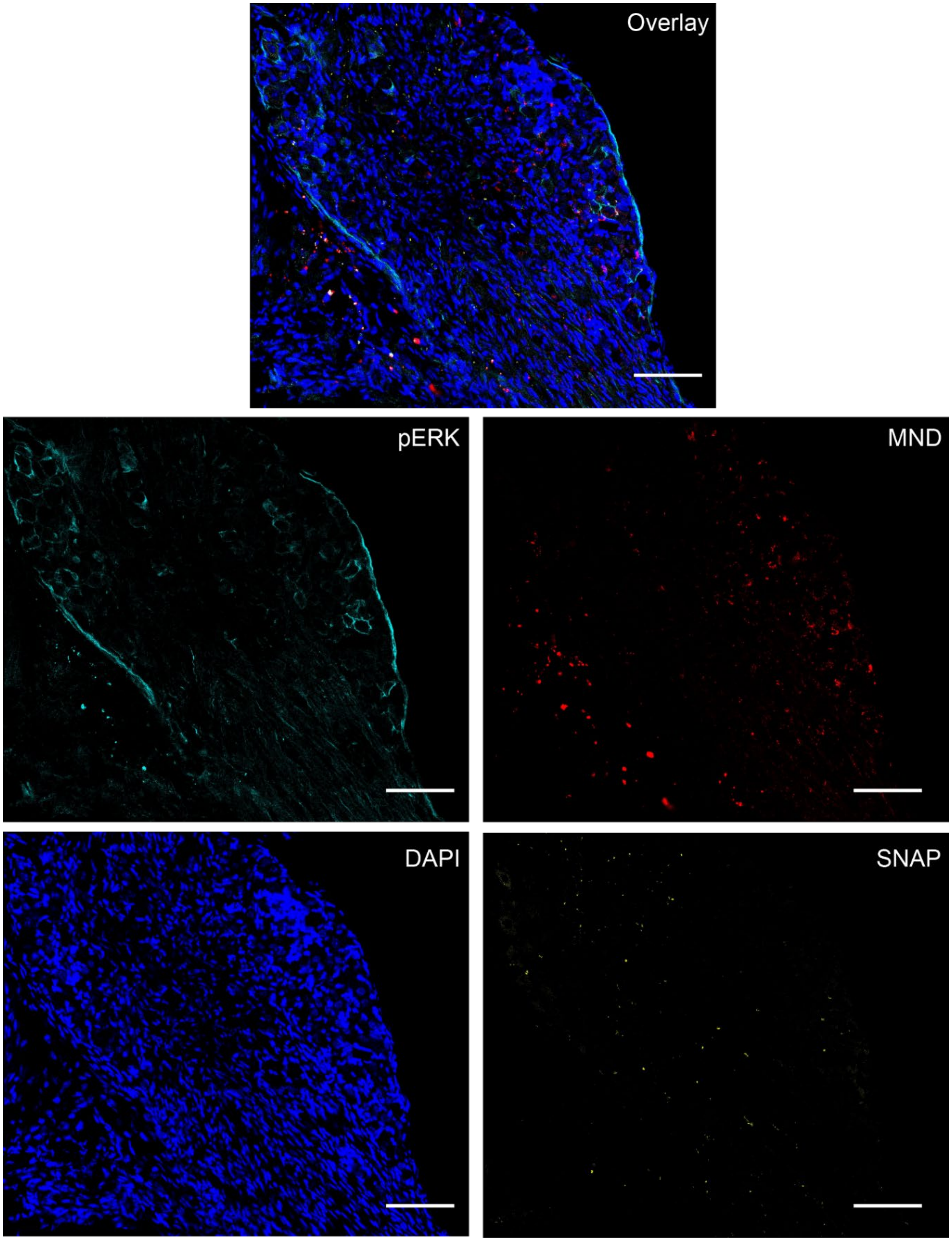


**Fig. S23 Oxtr-Cre$^{-/-}$ left NG.** Immunofluorescence image of the *left* NG in a Oxtr-Cre$^{-/-}$ mouse (control side without MND injection); **yellow indicates** SNAP-tag, cyan indicates pERK, **blue** indicates DAPI-labeled nuclei, and **red** shows Cy7 fluorescence marking MND localization. Scale bars, 100 μm.

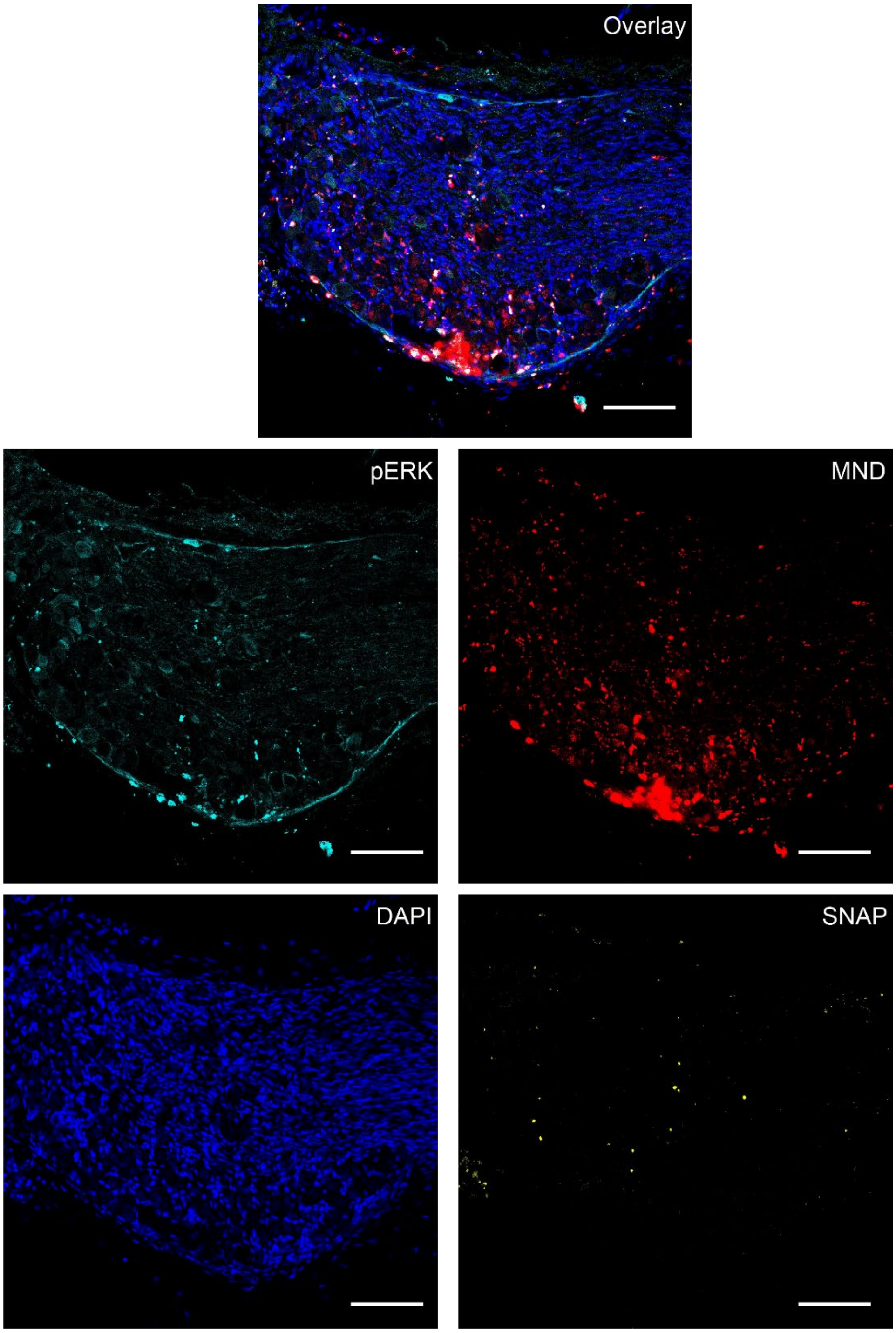


**Fig. S24 Glp1r-Cre$^{-/-}$ left NG.** Immunofluorescence image of the *left* NG in a Glp1r-Cre$^{-/-}$ mouse (control side without MND injection); **yellow indicates** SNAP-tag, cyan indicates pERK, **blue** indicates DAPI-labeled nuclei, and **red** shows Cy7 fluorescence marking MND localization. Scale bars, 100 µm.

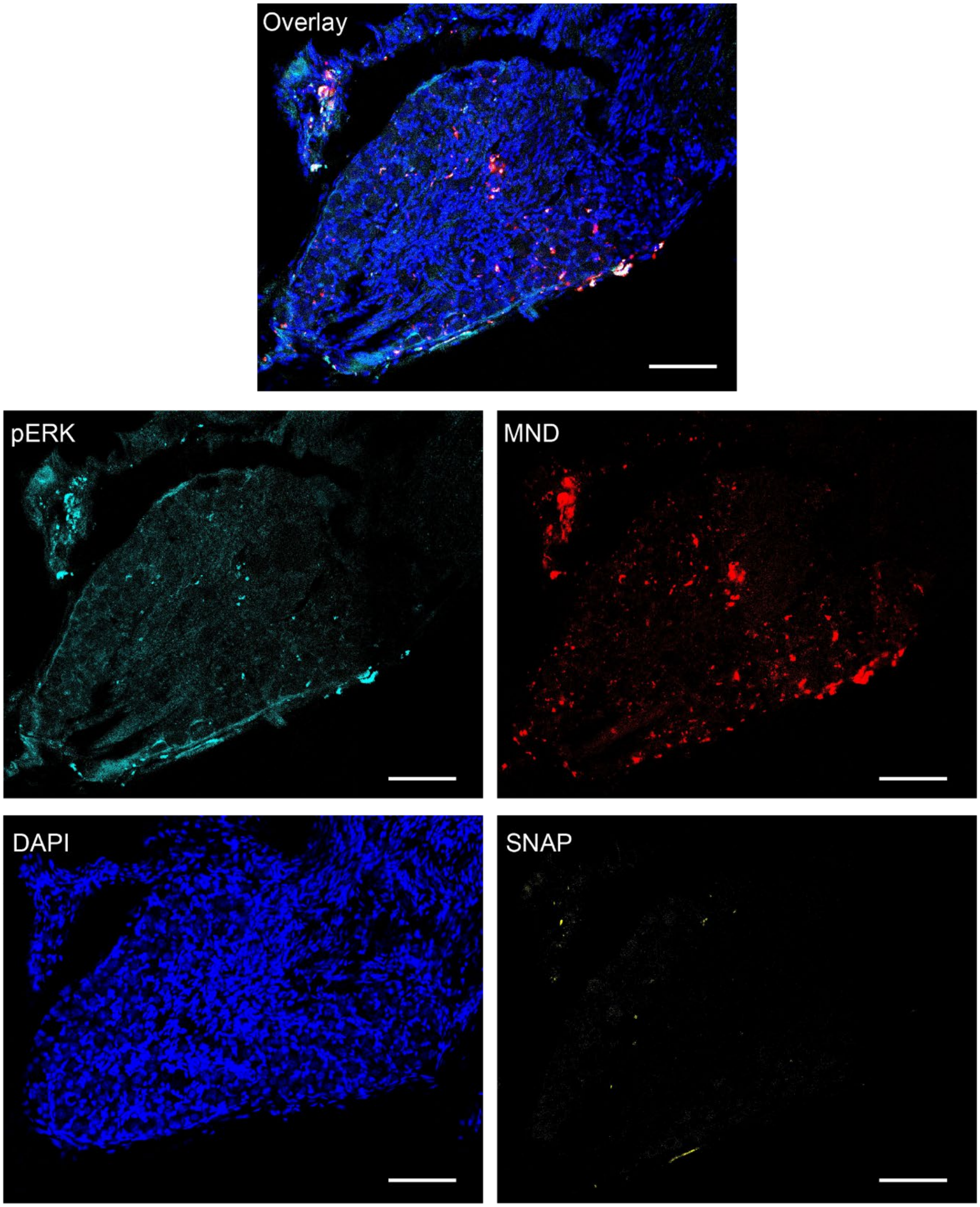


**Fig. S25 Gpr65-Cre$^{-/-}$ left NG.** Immunofluorescence image of the *left* NG in a Gpr65-Cre$^{-/-}$ mouse (control side without MND injection); **yellow indicates** SNAP-tag, cyan indicates pERK, **blue** indicates DAPI-labeled nuclei, and **red** shows Cy7 fluorescence marking MND localization. Scale bars, 100 µm.

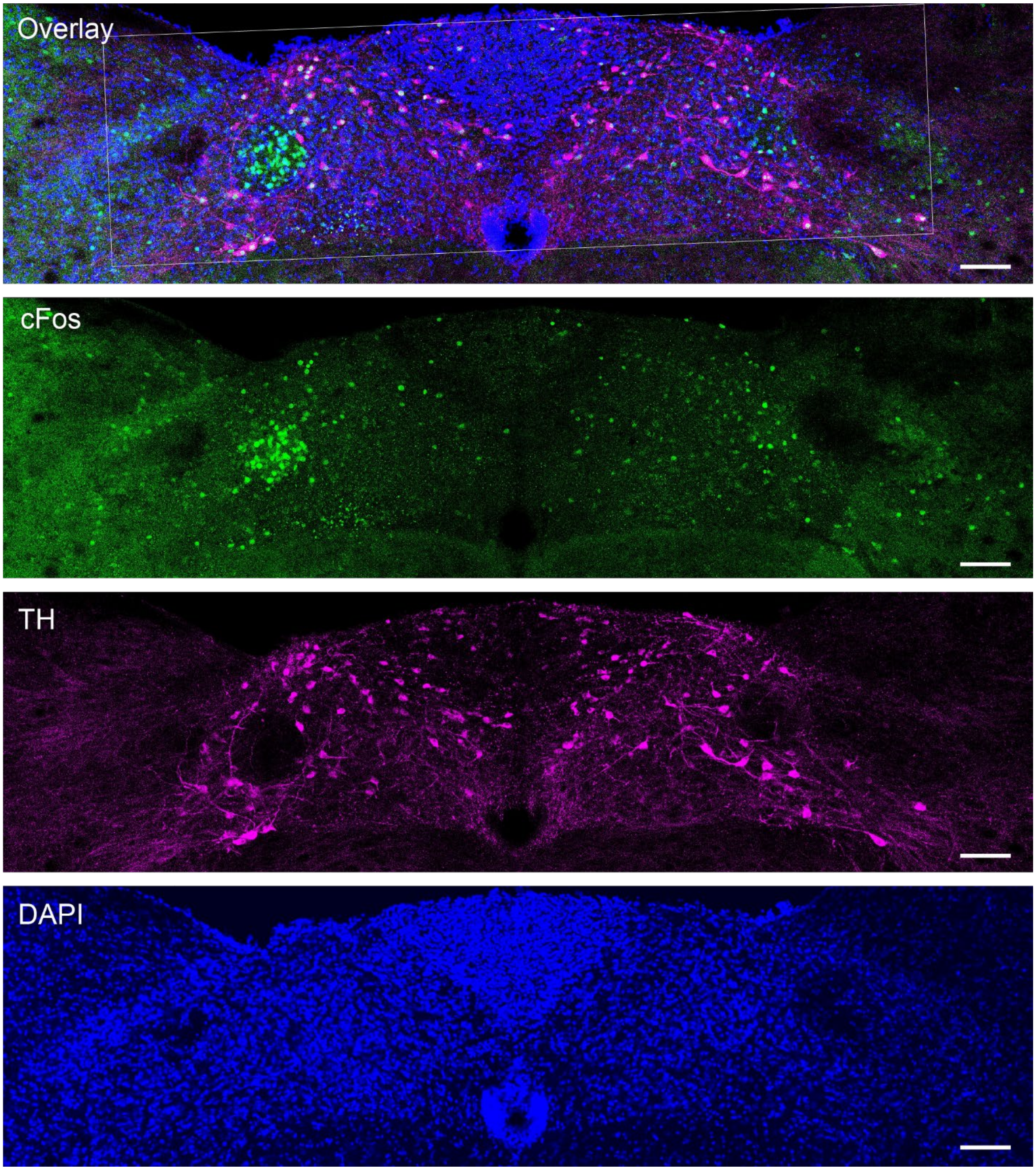


**Fig. S26 Oxtr-Cre$^{+/-}$ NTS.** Immunofluorescence images of NTS sections from Oxtr-Cre$^{+/-}$ mouse; **Green**, c-Fos; **magenta**, Th; **blue**, DAPI. The white square indicates the portion in Fig, 4. Scale bars, 100 µm.

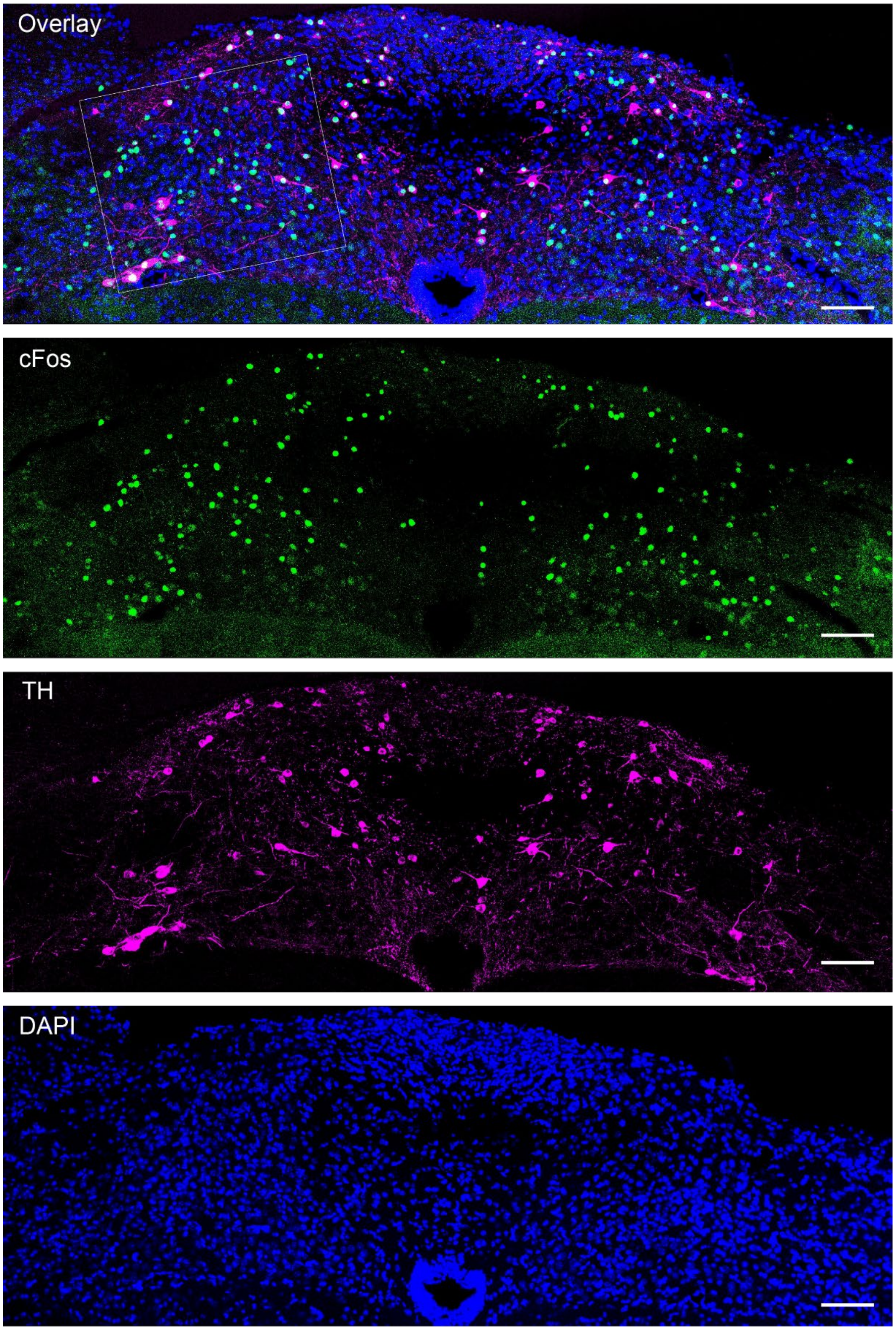


**Fig. S27 Glp1r-Cre$^{+/-}$ NTS.** Immunofluorescence images of NTS sections from Glp1r-Cre$^{+/-}$ mouse; **Green**, c-Fos; **magenta**, Th; **blue**, DAPI. The white square indicates the portion in Fig, 4. Scale bars, 100 μm.

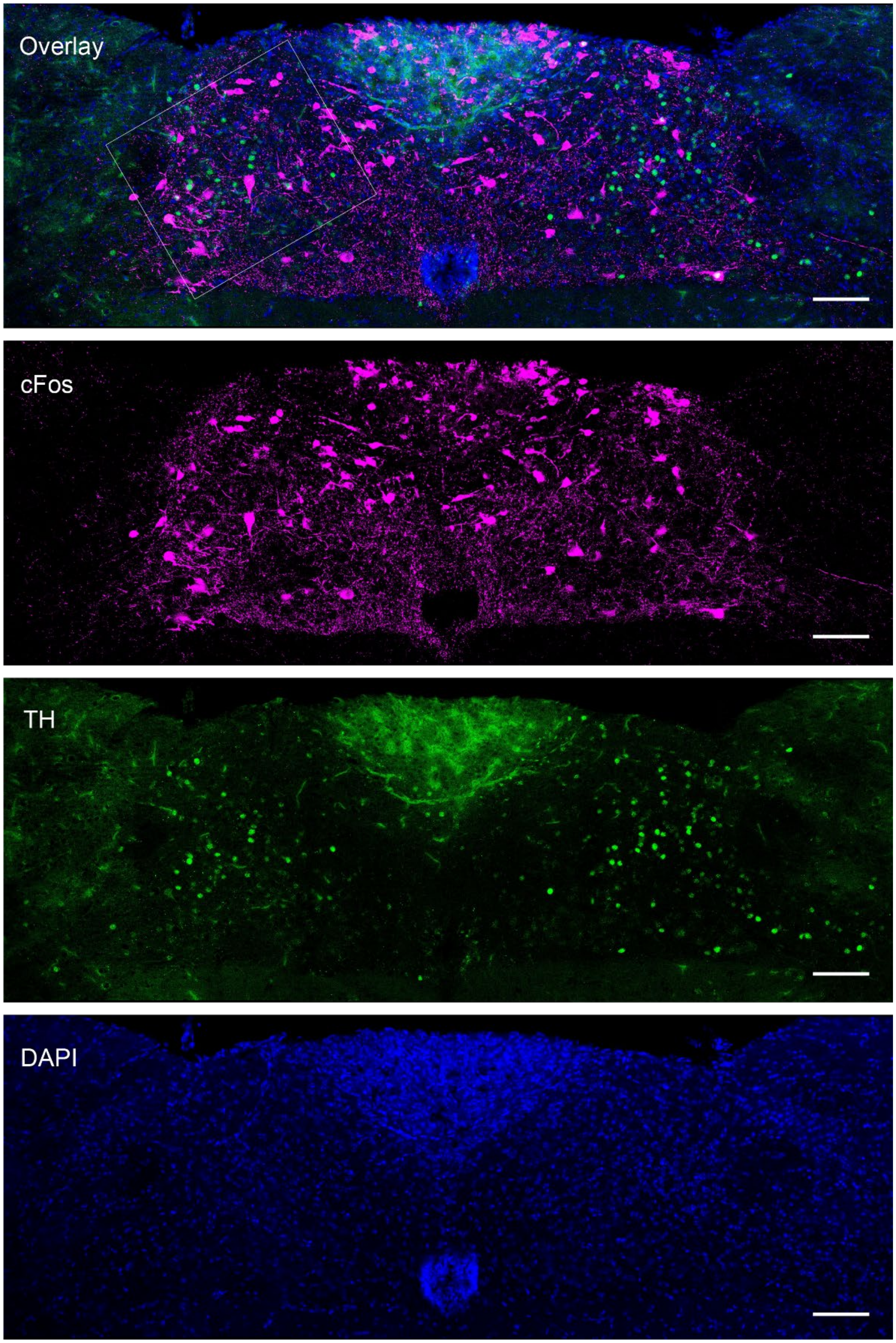


**Fig. S28 Gpr65-Cre$^{+/-}$ NTS.** Immunofluorescence images of NTS sections from Gpr65-Cre$^{+/-}$ mouse; **Green**, c-Fos; **magenta**, Th; **blue**, DAPI. The white square indicates the portion in Fig, 4. Scale bars, 100 μm.

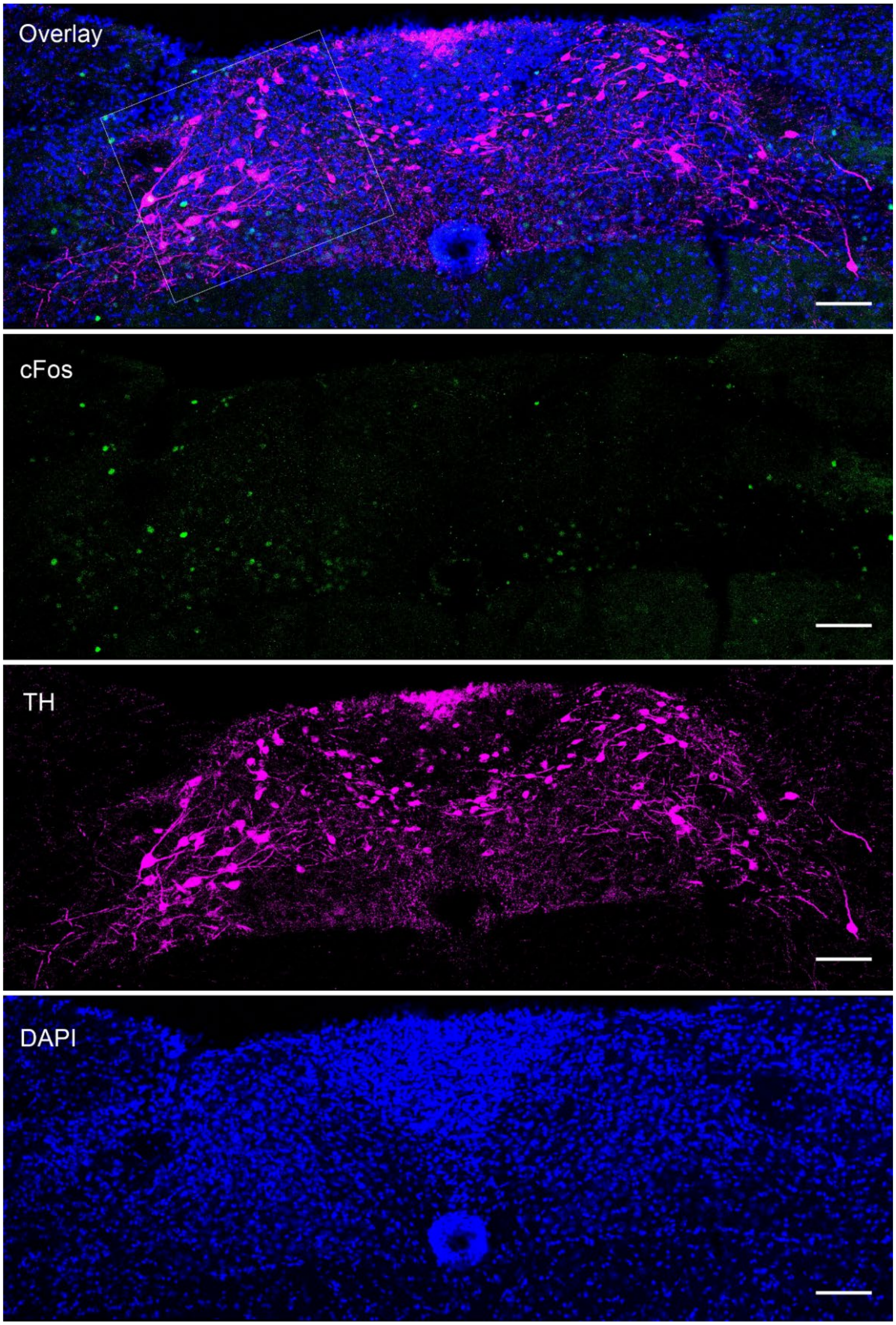


**Fig. S29 Oxtr-Cre$^{-/-}$ NTS.** Immunofluorescence images of NTS sections from Oxtr-Cre$^{-/-}$ mouse; **Green**, c-Fos; **magenta**, Th; **blue**, DAPI. The white square indicates the portion in Fig, 4. Scale bars, 100 µm.

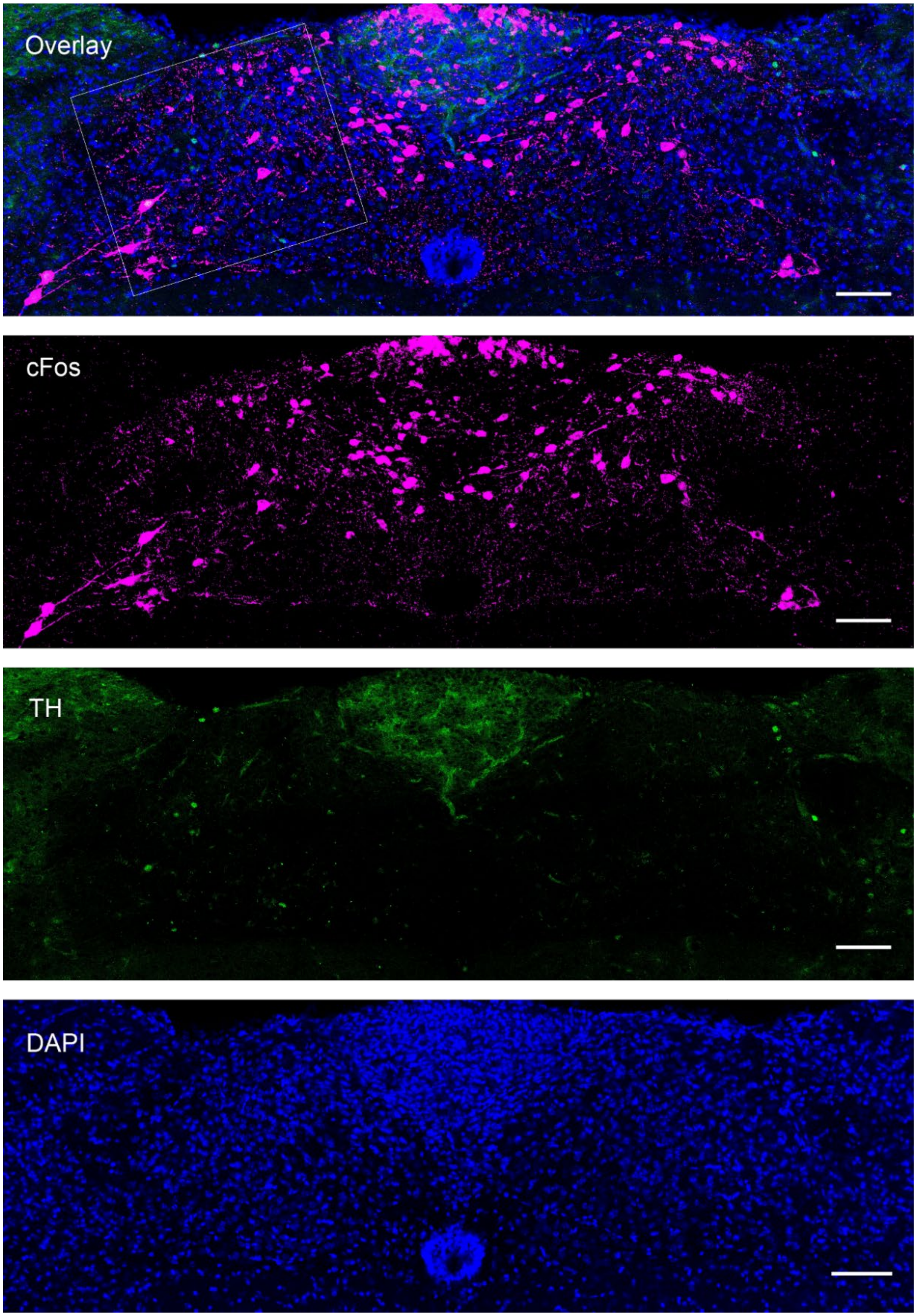


**Fig. S30 Glp1r-Cre -/- NTS.** Immunofluorescence images of NTS sections from Glp1r-Cre$^{-/-}$ mouse; **Green**, c-Fos; **magenta**, Th; **blue**, DAPI. The white square marks the section presented in Fig. 4. Scale bars, 100 µm.

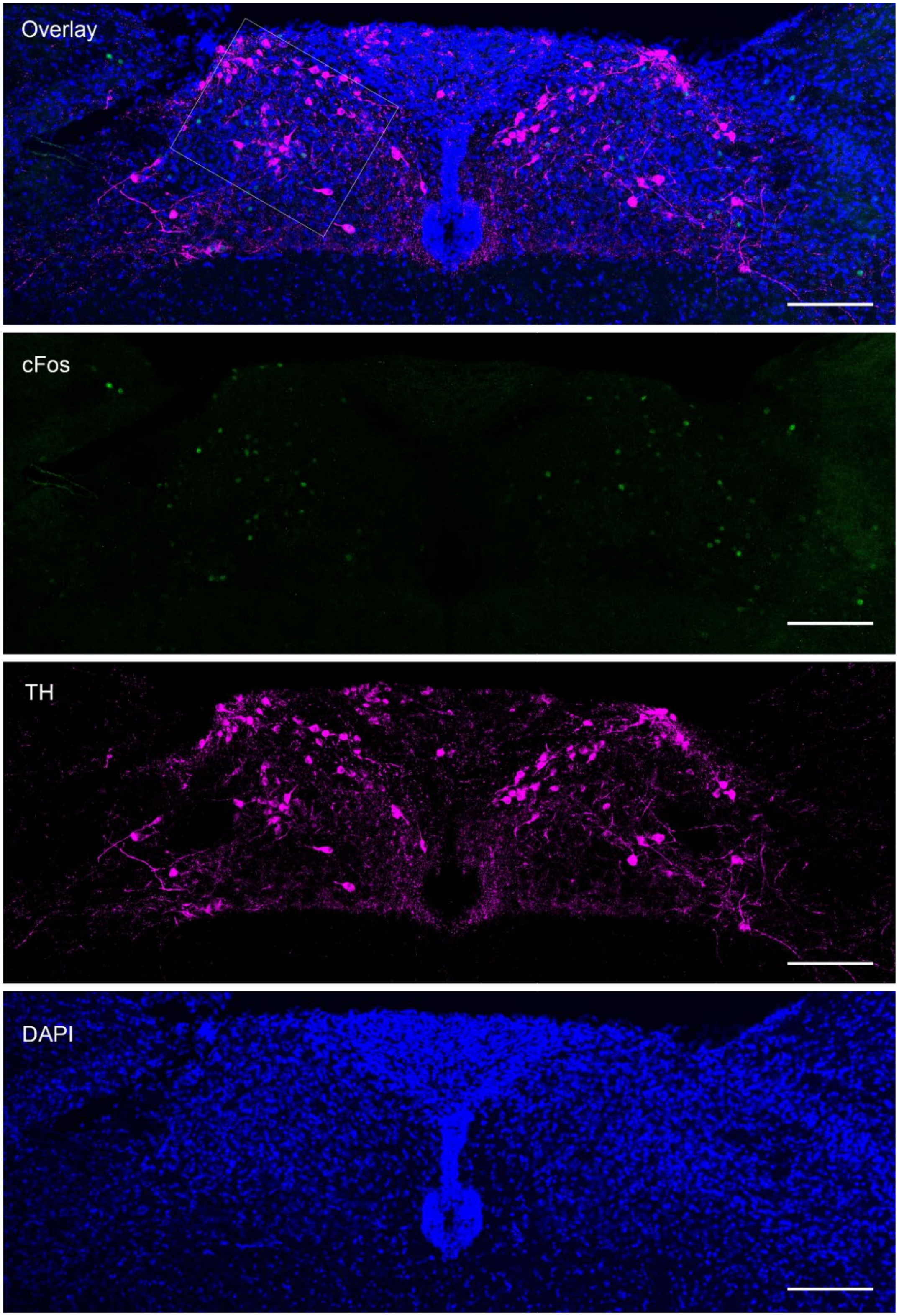


**Fig. S31 Gpr65-Cre -/- NTS** Immunofluorescence images of NTS sections from Gpr65-Cre$^{-/-}$ mouse; **Green**, c-Fos; **magenta**, Th; **blue**, DAPI. The white square marks the section presented in Fig. 4. Scale bars, 100 μm.

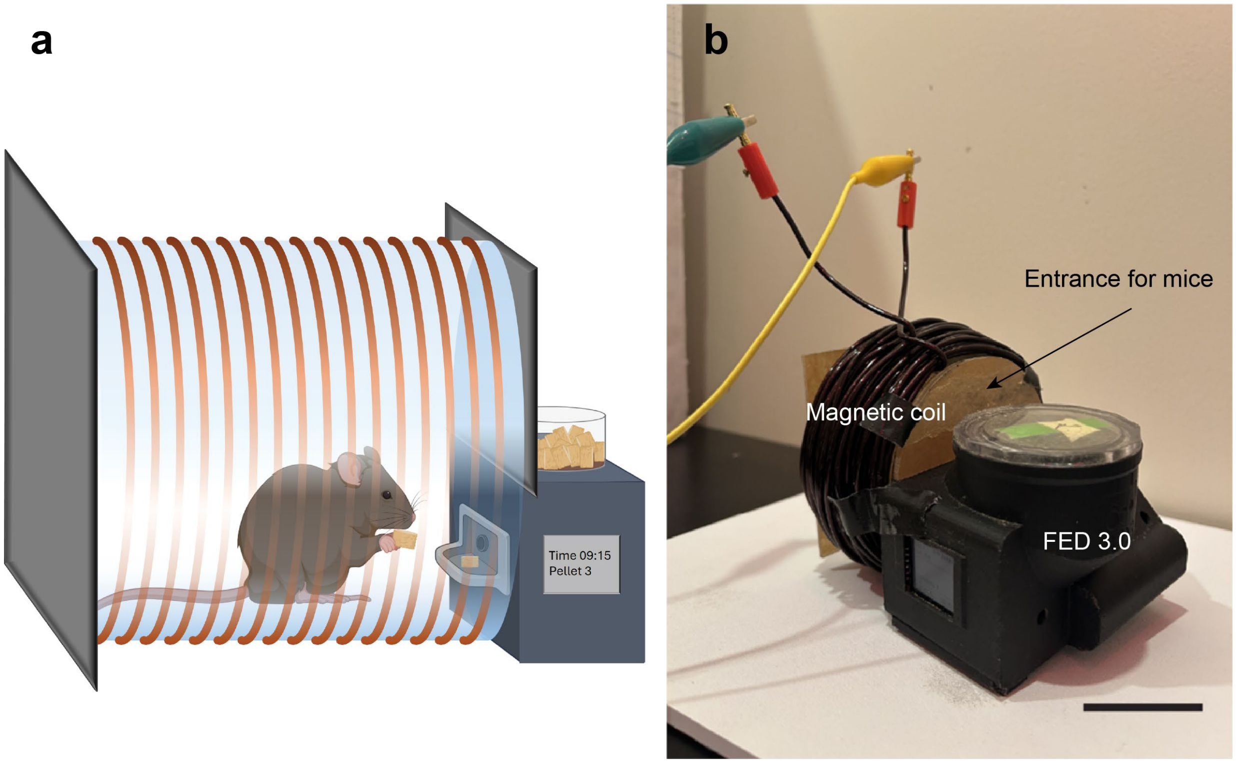


**Fig. S32 Food intake assay. a,** Scheme illustrating a mouse positioned within a magnetic coil for alternating magnetic field (MF) generation, which is integrated with the FED3.0 system. Upon pellet retrieval from the food-dispensing port by the mouse, the system delivers the next pellet within 3 s, thereby marking the time of pellet acquisition. **b,** Picture of the system. Scale bar, 5 cm.

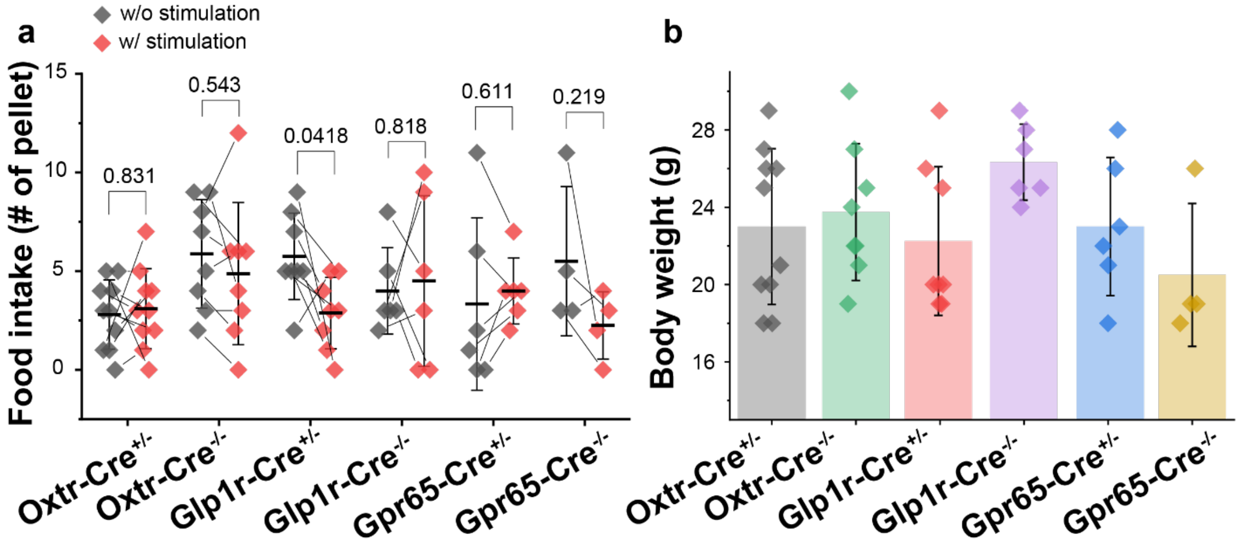


**Fig. S33 Food intake after 30 min MF application and body weight. a,** Number of pellets consumed during the 30-min MF stimulation period only. As the data distribution follows normality, paired sample t-test was performed; Oxtr-Cre$^{+/-}$ t=-0.22, DF=9; Glp1r-Cre$^{+/-}$ t=2.49, DF=7; Gpr65-Cre$^{+/-}$ t=-0.54, DF=5; Oxtr-Cre$^{-/-}$ t=1.04, DF=7; Glp1r-Cre$^{-/-}$ t=-0.242, DF=5; Gpr65-Cre$^{-/-}$ t=1.55, DF=3. **b,** Body weight. As the data did not meet normality assumptions, group comparisons were performed using a Kruskal–Wallis test with Dunn's post hoc correction; $\chi^2$ = 6.98, DF=5, p=0.222. Data shown as the mean ± s.d.

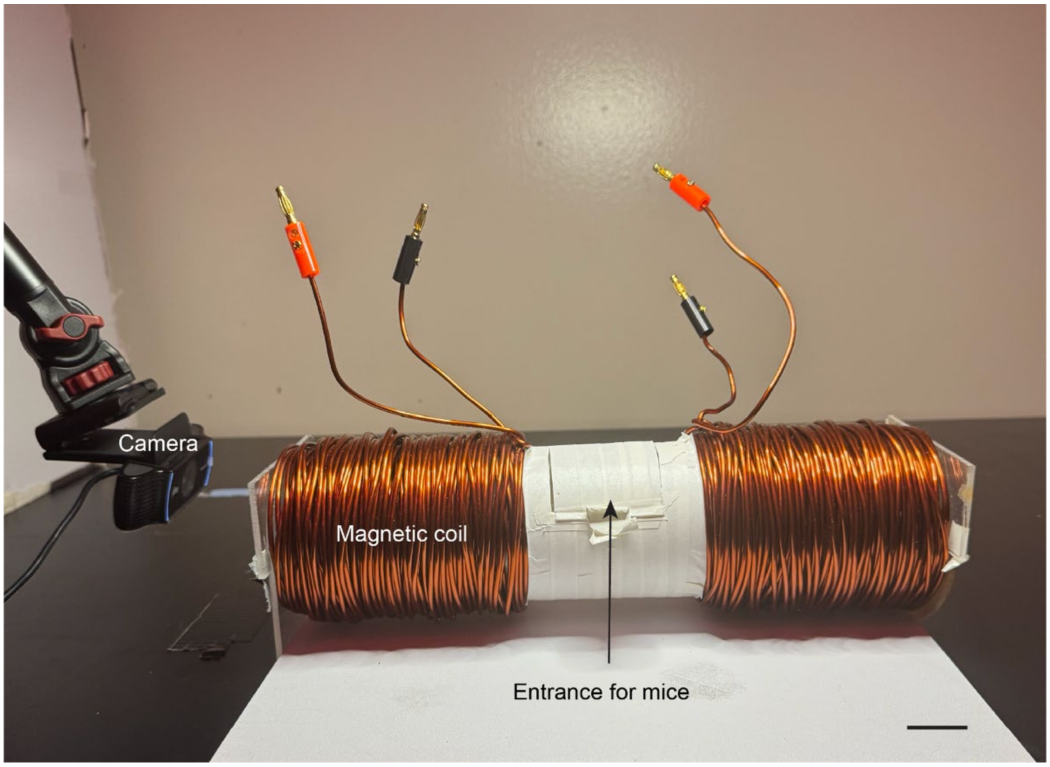


**Fig. S34 Place preference assay.** Scale bar, 2 cm.

**Supplementary Video S1.**

Simulation of magnetization ($\boldsymbol{m}$) and the instantaneous angular velocity $\boldsymbol{\omega_r}$ of MNDs for the two cycles of applied magnetic field (B, 80 mT 50 Hz). Frame rate is equal to 0.001 of the normalized phase (angular velocity ($\boldsymbol{\omega}$) × time (t)) / 2$\boldsymbol{\pi}$.

**Supplementary Video S2.**

GCaMP6s fluorescence dynamics (5× speed, recorded at a rate of 20 frames per second) in primary hippocampal neurons with (left) and without (right) SNAP-tag expression decorated with BG-functionalized MNDs during a 30 s baseline and during five 2 s magnetic field (MF) epochs (80 mT, 50 Hz) separated by 10 s rest periods.

**Supplementary Video S3.**

GCaMP6s fluorescence changes (10× speed) in neurons in left nodose ganglion injected with BG-functionalized MNDs with (right) and without (left) SNAP-tag expression in $Oxtr^+$ neurons at baseline and following four 4 sec and four 10 sec MF epochs (80 mT, 50 Hz), recorded at 5 frames per second.
